# Design of novel synthetic promoters to tune gene expression in T cells

**DOI:** 10.1101/2025.01.09.632034

**Authors:** Giuliano Bonfá, Giovanna Martino, Assunta Sellitto, Antonio Rinaldi, Fabiana Tedeschi, Fabio Caliendo, Loris Melchiorri, Daniela Perna, Freddie Starkey, Evangelos Nikolados, Filippo Menolascina, Velia Siciliano

## Abstract

Transcriptional control of transgene expression can be linked to dynamic changes in cellular states if this is accompanied by differential expression of transcription factors (TFs). Synthetic promoters (SPs) designed to respond to the desired TFs can provide this regulation with compactness, specificity, and orthogonality. T cells display differentially expressed TFs according to the functional state. In solid tumors, the highly immunosuppressive TME and the chronic exposure to antigens lead to a progression of T cells from a functional to a dysfunctional state known as exhaustion (Tex), in which their power against cancer cells is strongly compromised. Importantly, this transition is accompanied by a marked increase of several TFs, among other factors, that drive targeted genetic programs. Strategies to detect and mitigate Tex are extremely needed. Here, we design SPs that respond to TFs differentially expressed in activated and exhausted T cells to enable new classifiers of the functional/dysfunctional states. We developed a library of over 80 SPs responsive to 7 TFs. The SPs showed broad strength of activation of reporter genes or immunomodulatory molecules in HEK293 and Jurkat T cell lines. Moreover, using a transfer learning strategy we show SPs strength predictability. By combining SPs responding to different TFs, we created Boolean logic gates and implemented a feed-forward design that was previously shown to reduce noise in the OFF-state. Finally, as proof of principle, we demonstrate the dynamic activation of the NR4A2-responsive SP according to the T cell state in primary human CD8+ T cells. Collectively we present a sensing platform that provides a versatile tool to study and monitor the dynamic changes occurring in T cells. In perspective the biosensors coupled to therapeutic genes can be used to reprogram the TME and reinvigorate the T cell anti-tumoral functionality, preventing or reverting the exhausted phenotype.

**Graphical Abstract:** Created with BioRender.com.

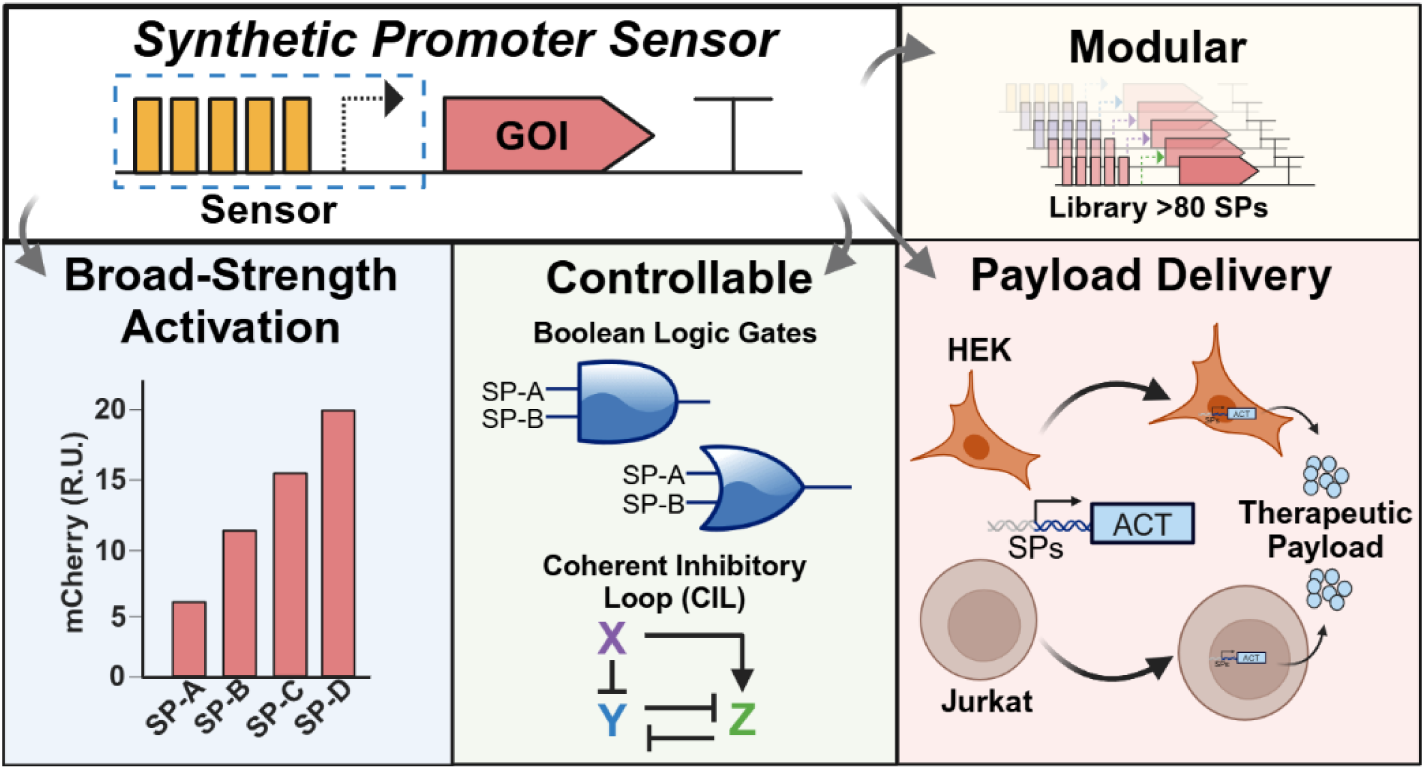

## INTRODUCTION

Realizing the potential of mammalian synthetic biology for biomedical applications requires the construction of complex genetic circuits with predictable behaviors ^1,2^, and the ability to control their expression with high robustness to prevent off-target activations ^3,4^. Over the years, synthetic biology has introduced innovative tools and strategies to achieve cell-specific and spatiotemporal control of therapeutic gene expression. These include transcriptional and post-transcriptional systems, chemogenetics, optogenetics, and feedback-based circuit topologies ^5,6^.

Transcriptional regulation remains the most widely employed strategy, particularly when integrating genetic payloads into target cells. However, the use of strong constitutive promoters can lead to gene expression levels that risk dose-dependent side effects, raising significant safety concerns ^7^. Alternatively, genetic circuits can be regulated by artificial transcriptional systems (ATS) that employ either engineered transcription factor (TF) and cognate synthetic promoters, or Zinc finger, Talen and CRISPR-based TFs to target endogenous promoters. The amplitude of promoter activation can be tuned using activation domains (AD) of varying strength, offering a safer and more precise approach to gene regulation ^8–11^.

Transcriptional control of transgene expression can be also linked to dynamic changes in cellular states if this is accompanied by differential expression of TFs ^12,13^. In this case the synthetic promoters (SP) are designed to be highly specific for the TF families ^14^, with designs considerations that enable compactness, specificity, and orthogonality ^15–17^. This places SPs in a privileged position compared to endogenous promoters, which often exhibit crosstalk due to the presence of enhancer or binding sites responding to several different TFs. The potential of SPs for cancer immunotherapy had already demonstrated to enhance immunomodulatory circuit selectivity in a tumor-specific fashion to enhance safety of output control for clinical applications. Indeed, NFAT- and NR4A-based promoters that respond to TFs induced by antigen stimulation of T cells have shown to control cytokine expression in engineered T cells ^14,17^. Thus, CAR-T cells that are armored with therapeutic payloads such as inflammatory cytokines to revamp the immunosuppressive tumor microenvironment (TME), could benefit from the use of SP ^7,18^. T cells display differentially expressed TFs according to the functional state. In solid tumors the highly immunosuppressive TME supported by the M2 tumor associated macrophages (M2 TAMs), and the chronic exposure to antigens lead to a progression of T cells from a functional state, characterized by high proliferation and cytotoxic capabilities to a dysfunctional state known as exhaustion (Tex), in which their power against cancer cells is strongly compromised. Tex cells exhibit a different epigenetic profile, have a distinct phenotype, reduced functionality and cytotoxicity ^19–21^. Importantly, the transition from functional to exhausted T cells is also accompanied by a marked increase of several TFs, that drive targeted genetic programs ^22–27^.

Here, we designed a library of synthetic promoters that respond to transcription factors differentially expressed in activated and exhausted T cells to enable new classifiers of the functional/dysfunctional states. We first carried out a bioinformatics analysis to define TFs that exhibits different levels of expression in activated and exhausted T cells, using expression data collected from pioneer studies in the field ^22,28,29^. We then created a modular platform for the rapid engineering of a library of roughly 80 SPs responding to 7 TFs. These sensors demonstrated to be modular, tunable and showed broad strength activation with potential to release a therapeutic payload in engineered cells. To have compelling information on the ON-OFF state, we evaluated leakiness and absolute induction in HEK293 cells and in Jurkat T cells. The latter is a T-derived cell line that works as the best approximation for the promoter screening prior using them in primary T cells. Further, we integrated the SPs in engineering architectures that improve the ON-OFF state ^30^, and built AND and OR gates to compute TFs- specific information processing. Using a transfer learning strategy we show SPs strength predictability. Finally, as proof of principle, we demonstrated the NR4A SP functionality across T cells states in an *in vitro* model of exhaustion.

## RESULTS

### Design of a modular architecture for rapid construction of synthetic promoters

To create transcriptional classifiers of T cells functional states, we first carried out a bioinformatics analysis to identify TFs that are robustly upregulated at different stages of activation and exhaustion (**Fig. 1A**). To this end, by exploring publicly available gene expression data, we generated a list of 197 TFs always upregulated in exhausted vs effector/memory T cells across chronic infections and cancer ^28,29,31^. We then refined our analysis using RNA-Seq data from functional and exhausted human T cells from positional papers in the field of T cell and CAR-T cell immunology ^31–34^. Among the differentially expressed TFs we selected a subset of seven TFs (IRF4, BATF, MAF, GATA3, NR4A2, EOMES and IKZF2) based on: (i) expression in the early stages of activation, functional effector/memory and in dysfunctional cells, and (ii) on their well characterized role in promoting or augmenting dysfunctional states ^22–24,27,35^ (**Fig. 1A and Supplementary Table S1**).

**Figure 1.**
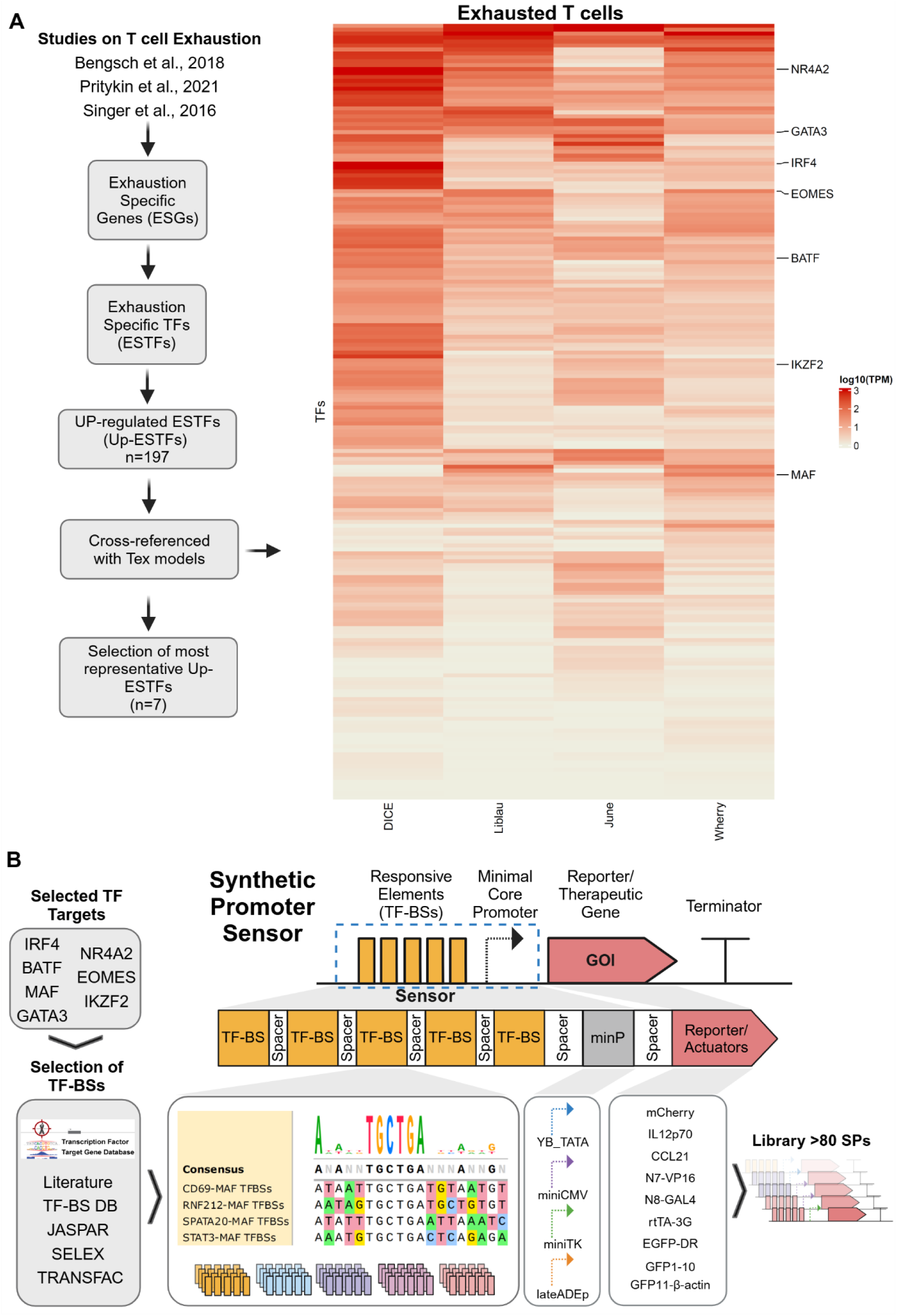
Bioinformatics selection of transcription factors (TFs) differentially expressed in activated and exhausted T cells. (**A**) Workflow strategy to select targetable TFs from Exhaustion-Specific Genes (ESGs) to most representative Upregulated Exhaustion-Specific TFs (Up-ESTFs (**left**). Heatmap showing the median centered expression values (log10TPM- transcript per million) of 197 dysfunction-associated TFs in activated and exhausted T cells. Euclidean distance has been applied to cluster genes (**right**). (**B**) General structure and design workflow of synthetic promoter (SPs) to sense the seven pre-selected TFs (IRF4, BATF, MAF, GATA3, NR4A2, EOMES and IKZF2). The sensor is composed by a responsive region where TF Binding Sites (TF-BSs) are in 5x tandem repetition upstream of a minimal core promoter (minP), following specific spacers between each repetition (10-15 bps) and from last BS to minP (20 bps). TF-specific binding sites (BSs) were selected from literature and TF-BSs repositories (such as JASPAR, SELEX and TRANSFAC, http://tfbsdb.systemsbiology.net/). Four minP were used (YB_TATA, miniCMV, miniTK and lateADEp) to drive the expression of fluorescent reporter genes (mCherry, EGFP-DR and splited EGFP), inducible and split proteins for logic circuits compositions (rtTA, N7-VP16, N8-GAL4) or therapeutic proteins (IL12p70 and CCL21), generating a library of over 80 SPs. Created with BioRender.com.

The SPs are composed of three main modules: the transcription factors binding sites (TF-BSs), a minimal core promoter (minP) and a reporter gene (**Fig. 1B**). TF-BSs) were identified from TFBS Data Bank (TFBS-DB) and selected based either on reported binding of the TF, or on the likelihood to be targeted by the TF. TF-BSs were cloned in 5x tandem repetition, upstream the minP module, alternating each TF-BSs with spacers of 10-20bps **(Supplementary Tables S2A and B**) ^14,36^.

To investigate the impact of minP on SP leakiness and activation we initially combined TF-BSs with four different minP (YB-TATA, miniCMV, miniTK, LateADEp). A mCherry fluorescent protein was inserted downstream the SPs as output. Since each TF can recognize different DNA responsive elements, by using this combinatorial modular design we generated a library of over than 80 SPs sensors (**Fig. 1B and Supplementary Tables S2A and B**).

### Basal activity of synthetic sensors depends on minimal core promoters and TF-BSs

Leakiness of inducible promoters hampers the fine control of genetic circuits. This issue is particularly critical in applications such as gene and cell therapy, where stringent regulation is essential to mitigate adverse effects ^36–38^. Since basal activity may be associated with the nature of the responsive elements, including number of repetitions, or the characteristics of the minimal promoters that initiate transcription, we first characterized the leakiness of the minP alone or combined with the TF-BS of IRF4/BATF (that regulate the promoters of endogenous genes in T cells cooperatively ^39,40^, MAF and GATA3 by mCherry expression in HEK293 and Jurkat cells (**Fig. 2A**).

**Figure 2.**
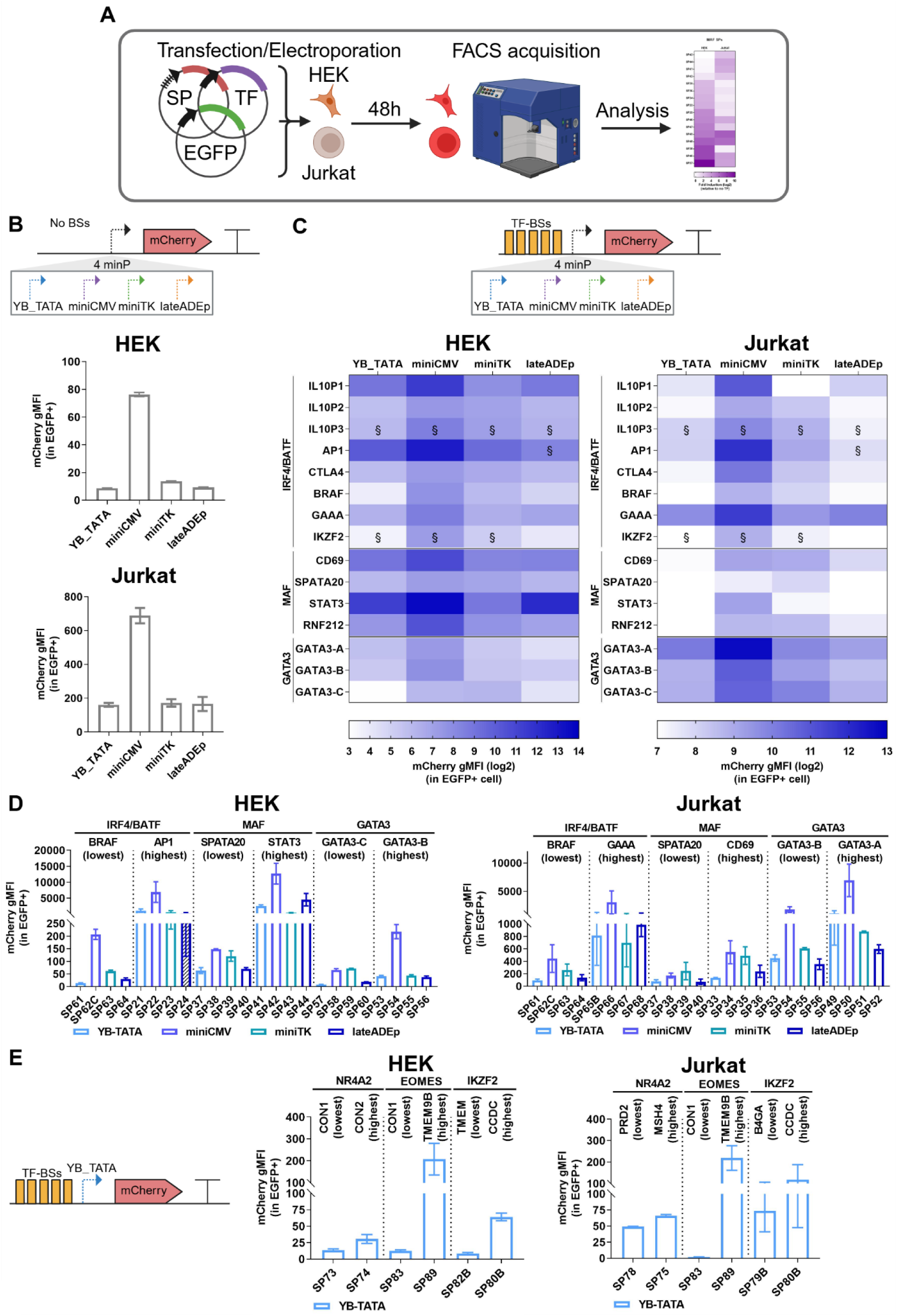
Investigation of parameters affecting the background noise of the SPs in the OFF state. (**A**) General experimental workflow of the screening process using the SPs library. SP sensors were characterized by transient transfection or electroporation in HEK293 and Jurkat cell lines, respectively. Three constructs are introduced into the cells: the SP, the cognate TF, and a constitutively expressed EGFP as transfection control. After 48 hours, we evaluated by flow cytometry (FACS) the expression of the mCherry fluorescent reporter. A scramble control in which the TF-BSs were not inserted were used to check basal activities of the minimal promoter. (**B**) Leakiness of the four minP (miniTK, lateADE, YB_TATA, mCMV), reported by mCherry gMFI 48h post-transfection/electroporation in HEK293 and Jurkat cells (**C-D**) Leakiness of minP combined to TF-BS of IRF4/BATF, MAF, GATA3, reported by mCherry gMFI 48h post-transfection/electroporation in HEK293 and Jurkat cells. (**E**) Leakiness of YB_TATA minP combined to TF-BS of NR4A2, EOMES, IKZF2 reported by mCherry gMFI 48h post-transfection/electroporation in HEK293 and Jurkat cells. (**D-E**) SPs showing the highest and lowest leakiness were grouped by cell type and expressed in bar- plots. §: SPs containing less than 5x TF-BSs repetition. N=2 biological replicates. Created with BioRender.com.

All minP except miniCMV showed low basal activity both in HEK293 and Jurkat cells in the absence of TF-BSs (**Fig. 2B**). Conversely, when combined to the TF-BSs we observed a broad range of background noise that correlated the TF-BS (**Fig. 2C**), and the YB_TATA emerged as the least leaky in combination with the responsive elements particularly in Jurkat cells (**Fig. 2C-E and Supplementary Figure S2.1A-C**).

Among the SPs targeting IRF4/BATF, we observed low background noise with TF-BS derived from BRAF endogenous promoters (SP61-SP64, **Fig. 2D**). Of these, the lowest noise was achieved in combination with YB_TATA (SP61), while coupling with miniCMV (SP62C) resulted in higher leakiness in both HEK293 and Jurkat cells. Notably, high background noise was associated with two distinct groups of SPs in HEK293 and Jurkat cells: AP1-derived BS (SP21-24) in HEK293 and GAAA-derived BS (SP65B-68) in Jurkat cells (**Fig. 2D, Supplementary Figure S2.1B**).

Also for MAF SPs, both HEK293 and Jurkat showed minimal basal activity with SPATA20 promoter-derived BS (SP37-40), and the combination with YB_TATA and LateADEp exhibited the lowest noise levels. Conversely, highest background was associated to two distinct groups of SPs in HEK293 and Jurkat cells: STATA-3-derived BS (SP41-44) in HEK293, and CD69-derived BS (SP33-36) in Jurkat cells (**Fig. 2D, Supplementary Figure S2.1C**). However, it was interesting to observe that in Jurkat cells the leakiness was not as high as the other SPs. Last, all SPs targeting GATA3 exhibited minimal noise in HEK293 cells, whereas in Jurkat cells the background activity was observed in all combinations, albeit to a different extent (**Fig.2C-D**).

Collectively our data indicate that both YB_TATA and lateADEp are suitable to build SPs with low background. Since long-term expression in engineered cells is crucial to exploit therapeutic interventions, it is critical to minimize viral genetic elements in the delivery vectors to reduce the risk of silencing. As lateADEp has adenoviral origins ^41^, we then opted for YB_TATA to build the next set of SPs responding to NR4A2, EOMES and IKZF2.

NR4A2-responsive SPs showed low basal activity in HEK293 and Jurkat cells except for SP77A (NEG1, 4 Repetition) that exhibited high leakiness in Jurkat cells (**Fig. 2E, Supplementary Figure S2.2A-B**). Similar to NR4A2, also EOMES- and IKZF2-responsive SPs showed low basal activity in general for both cell lines tested (**Fig. 2E, Supplementary Figure S2.2A-D**).

Altogether, this initial characterization enabled the design of transcriptional sensors with low leakiness when coupled to YB_TATA. In contrast, miniCMV exhibited particularly strong basal activity, regardless of the specificity of the TF-BSs, generating high noise in the OFF state when combined with certain TF-BSs (e.g., SP22, SP42, SP50, SP54 and SP66) (**Fig. 2D**). Notably, background noise was cell-dependent, underscoring the importance of conducting preliminary screening designs in bona fide in vitro setups.

### The synthetic promoter sensors are inducible and show a wide range of activation

We next evaluated activation of the SPs by transient co-transfection/electroporation with and without the cognate TF in HEK293 and Jurkat cell lines (**Fig. 2A**). In general, IRF4/BATF, MAF, NR4A2 and EOMES SPs demonstrated to induce the expression levels of reporter mCherry by the TFs as compared to the control no-TF in both cell lines (**Fig. 3A, Supplementary Figures S3.2-3.7**). In contrast, GATA3-SPs reduced reporter expression in the presence of the TF in HEK293 cells, whereas the repression was milder in Jurkat cells. (**Supplementary Figures S3.6B-3.7C**). Indeed, GATA3 TF regulates histone modifications and can work as inducer or repressor, depending on the cell context ^42^. The **IRF4/BATF SPs** were tested in the presence of either the single TF or both, resulting in different induction pattern. In some designs (SP05-5X_IL10P1_BS and SP69A-3X_IKZF2_BS), BATF alone did not activate the SP, IRF4 did, and co-expression of both resulted in increased mCherry levels (**Supplementary Figures S3.2A-3.5A**). In other designs (e.g. SP61-5X_BRAF_BS) we observed mild activation with individual TFs and a synergistic effect with co-expression of both (**Fig.3A**). Some SPs, such as SP13A (3X_IL10P3_BS) and SP65B (GAAA), showed induction only by IRF4, especially in HEK293 cells (**Supplementary Figures S3.2A-3.5A**). Indeed, most of the TF-BSs selected to sense IRF4/BATF are composite of responsive elements to sense both TFs, whereas GAAA BS was selected as an IRF4 single motif control, explaining this induction pattern. We selected also a single responsive element for BATF, but it demonstrated high basal level with no differential induction, as such for SP21 (AP1) (**Supplementary Figures S3.2A**). This effect could be explained by the fact that this motif is shared and can be sensed by other TFs in the same AP1 TF family. Instead, SP25 (CTLA-4) showed a low induction activity in the presence of the single TFs and a synergistic expression when they were both together, (**Supplementary Figures S3.2A**). In general, the minimal core promoter showing lower leakiness and higher performance in these SPs targeting IRF4/BATF were YB_TATA and lateADEp (**Fig. 3A, Supplementary Figures S3.2-3.**).

**Figure 3.**
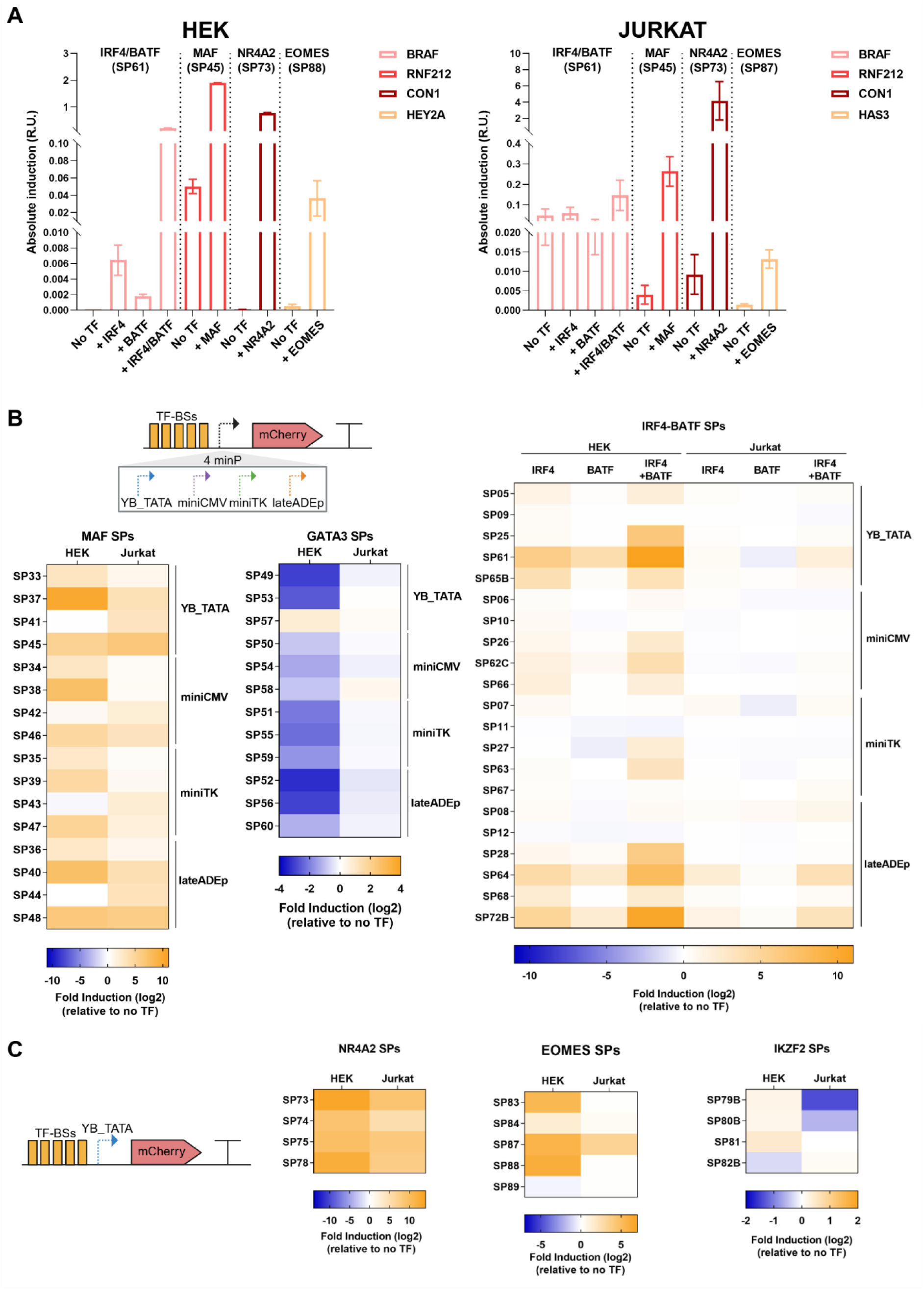
Evaluation of SPs activation. (**A**) Absolute induction of SPs showing the highest induction grouped by cell type expressed in bar plots. (**B-C**) Heat maps show fold induction of the SPs across different TF-BSs and across different minimal promoters or on just YB_TATA. The fold induction was calculated on normalized conditions using the mCherry and EGFP geometric means and percentage to calculate the absolute induction (**see Methods**) which was then referenced to the condition no-TF. The SPs best candidates were evaluated based on the ability to exhibit superior fold induction, which means high inducible activity in the presence of the correlated TF and minimal background (basal activity in the absence of the TF). N=2 biological replicates. Statistical test: one-way ANOVA with Tukey’s multiple comparison test (for IRF4/BATF SPs) or unpaired T test. p value: *<0.05; **<0.005; ***<0.0005; ****<0.0001. Created with BioRender.com.

The SPs responsive to MAF showed a more consistent activation both in HEK293 and Jurkat cells, in particular with SPATA20 (SP37-40) and RNF212 (SP45-48) BSs, in combination to all core promoters tested (**Supplementary Figure S3.6A**). The minP showing lower leakiness and higher activation was observed when coupled to YB_TATA and lateADEp (**Supplementary Figure S3.6A**). Highest reporter activation was observed for SP45 (RNF212) (**Fig. 3A**). SP33 (CD69) and SP37 (SPATA20) showed a moderate activation in HEK293 cells while SP41 (STAT3) was active in the absence of MAF. Different from HEK293 cells, SP33 and SP37 did not show induction in Jurkat, whereas SP41 exhibited mild activation, highlighting the differences the SPs could perform in different cell lines (**Supplementary Figure S3.6A**).

We confirmed that YB_TATA had lowest basal activities with most consistent induction across the SPs and was chosen to drive new SPs targeting NR4A2, EOMES and IKZF2 TFs. All NR4A2 SPs showed an ample range of activation and SP73 (CON1) induced highest reporter levels in both HEK293 and Jurkat T cells (**Fig. 3A and Supplementary Figure S3.7A**). The SPs targeting EOMES did not show high activation and among them the best performing were SP83 (CON1) and SP84 (CON2) in HEK293 cells and SP87 (HAS3) in Jurkat cells. Interestingly, SP88 (HEY2A) was effectively induced in HEK293 cells whereas in Jurkat cells the basal activity was already too high to show differences in the ON state (**Fig. 3A and Supplementary Figure S3.7B**). IKZF2 SPs did not show functional response, due in most cases to the high noise in the OFF state (**Supplementary Figure S3.7C**).

Given the deep characterization of noise at the OFF state and induction to the ON state, we were able to identify best compromises to achieve low leakiness and high induction. In the high leakiness scenario, we also propose a network topology to reduce the background.

To quantify the fold induction we performed the ratio of reporter expression in TF vs noTF controls (**Supplementary Figures S3.8-3.12**). In general, we observed a broad range of fold induction for almost all TFs targeted (**Fig. 3B and C and Supplementary Figures S3.8-3.12)**, with GATA3 SPs showing fold-repression instead (**Fig. 3B and C**). IRF4/BATF- responsive SP61 showed the highest fold induction in HEK293 cells and moderate fold induction in Jurkat cells. MAF-responsive SP37 and SP45 exhibited 1,134- and 39-fold activation respectively in HEK293 cells *versus* 11- and 86-fold respectively in Jurkat cells. Despite SP37 showed higher fold induction in HEK293 cells, SP45 showed the highest absolute activation in both cell lines (**Fig. 3A**). NR4A2-responsive SP73 showed 9,795-fold induction in HEK293 cells and 448-fold in Jurkat cells, proving the most consistent across experimental setting. EOMES-responsive SP88 exhibited 69-fold induction in HEK293 cells, but interestingly it was not activated in Jurkat cells. Conversely, EOMES-responsive SP87 showed 9.7-fold activation in Jurkat cells with 50-fold activation in HEK293. Our results confirm that even though some SPs showed elevated reporter activity, low to absent fold-induction was appreciated due to the high basal levels (no TF) they showed (**Fig. 3B and C and Supplementary Figures S3.8-3.12**).

Altogether, the designed SPs targeting IRF4/BATF, MAF, NR4A2 and EOMES are inducible, and the several SP configurations allowed us to achieve a broad range of fold induction of the reporter gene among the candidates, enabling a versatile tool to be adapted to sensing incrementing levels of the selected TF. This broader functionality will be essential during the choice of SPs to guide the expression of a therapeutic molecule.

We then sought to model promoter strength to capture relevant features for predicting gene expression given the SPs sequences. Due to the cardinality of the experimental dataset (n=82) we used a transfer learning strategy, a technique where a model pre-trained on one task is adapted to a different but related task (**Fig. 4A**). To apply transfer learning to our SPs task we use a dataset of approximately 25,000 human promoters from the Eukaryotic Promoter Database (EPD) ^43^. Our secondary EPD dataset contains characterization information for each promoter in the form of the number of ChIP-seq tags matching the 250bp region to each side of the transcription start site. Our approach consisted in training a deep neural network taking as input the EPD promoter sequences and predicting as output the expression in the form of ChIP-seq tag count (**Supplementary Note 1**).

**Figure 4.**
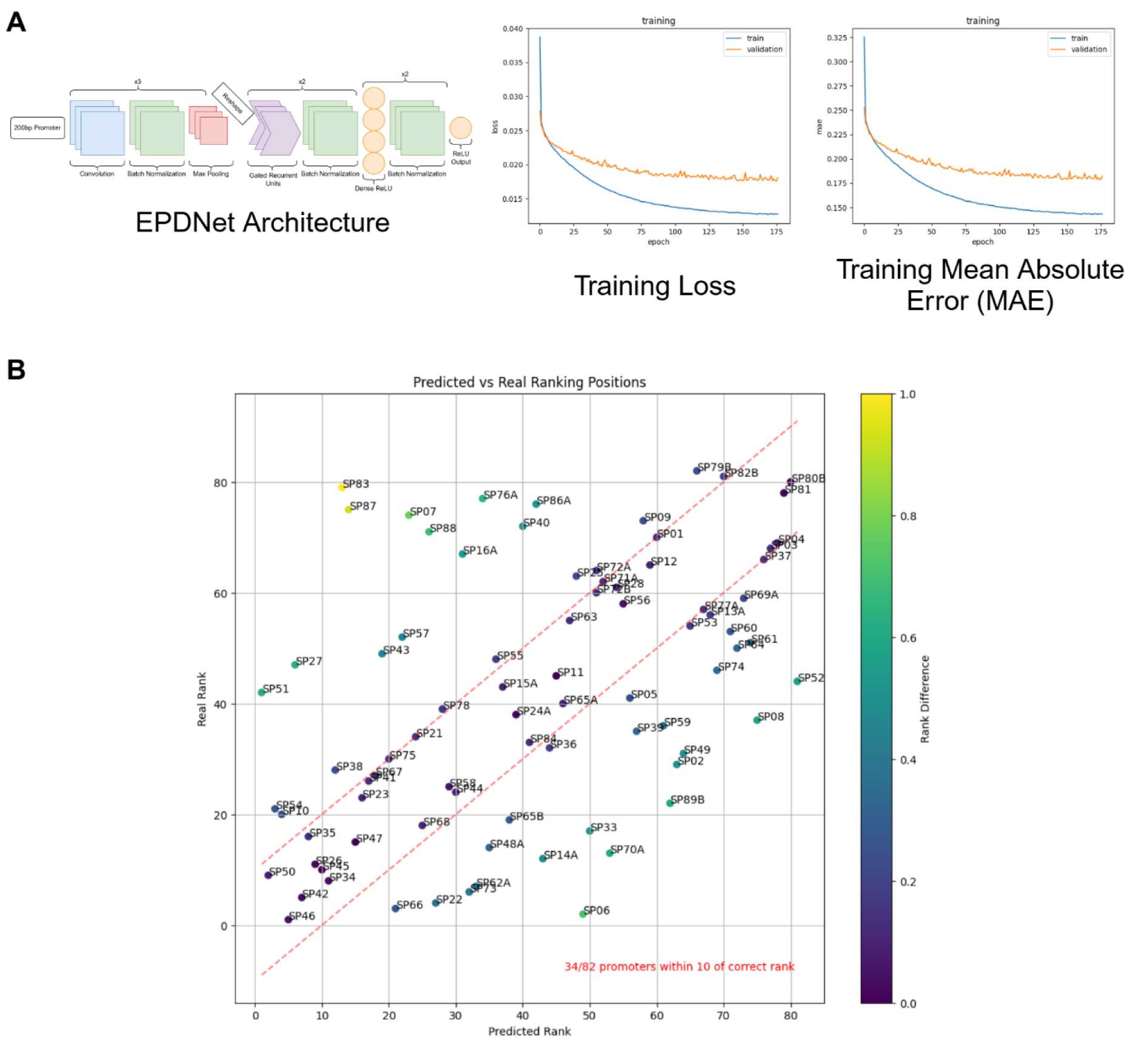
Transfer learning model performance showing measured vs predicted ranking. (**A**) Model architecture and training procedure. (**B)** The model produced a predicted ranking of the sequences based on their expected ’strength’. The mean absolute induction when the promoter is at its strongest expression was used to build the ranking to highlight the exact differences in the correct and predicted ranking for each individual promoter. Created with BioRender.com.

Having trained a network on the EPD promoter task we then applied it to the SPs which were ranked based on the fluorescence of the reporter gene. After training it on the EPD dataset, the neural network was applied to the 82 synthetic promoters. The fluorescence-based absolute mean induction of these promoters was Box-Cox transformed, and the model produced a predicted ranking of the sequences based on their expected strength. In each case we use the mean absolute induction when the promoter is at its strongest expression to build the ranking. The correlation between the predicted and actual rankings of the SPs was 0.49, indicating a moderate but significant agreement **(Fig. 4B)**. This result demonstrates that the transfer learning approach, leveraging a large dataset of human promoters, effectively captures relevant features for predicting gene expression in a distinct focused SP context despite the limited dataset size.

### Boolean logic gates and network topologies by synthetic promoters

Genetic circuits to regulate cell functionality require tight control to generate a response only if strictly defined conditions are fulfilled. We tested the robustness of two SP to implement multi-input synthetic sensors, by designing Boolean Logic gates that respond to MAF and NR4A2 TFs. By combining novel SP45 and SP73 engineered to respond to MAF and NR4A2 respectively, we developed an AND and an OR gate to induce the output when both TFs (AND) or at least one TFs is present (OR) (**Fig. 5A and D**).

**Figure 5.**
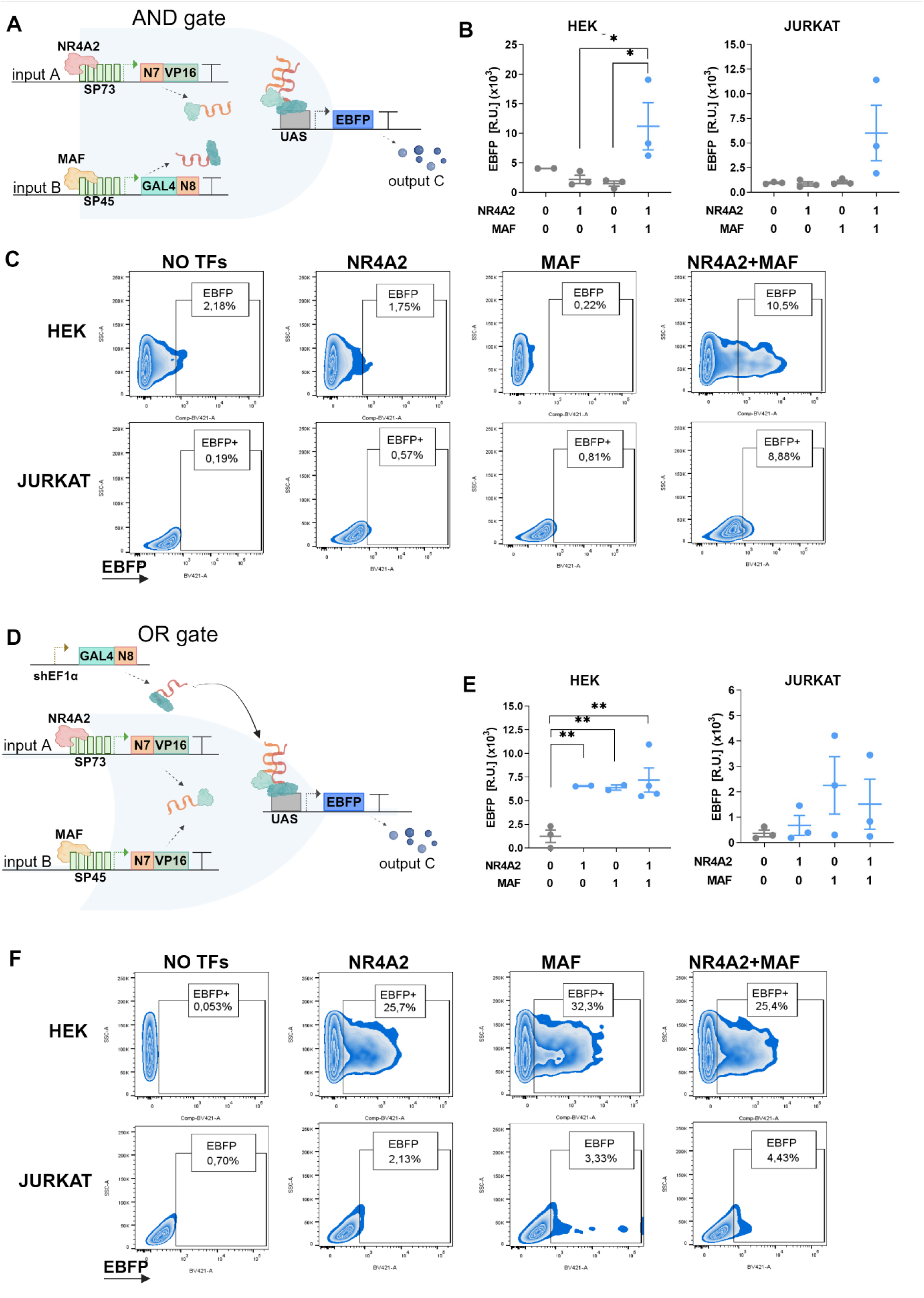
Implementation of Boolean Logic Gates and network topologies using MAF and NR4A2 SPs. (**A-C**) AND gate was composed by a split GAL4-VP16. GAL4 fused to the dimerization domain N8 is induced by MAF-responsive SP45 (input B), whereas VP16 fused to the dimerization domain N7 is controlled by NR4A2-responsive SP73 (input A). When both TFs are present, the two halves reconstitute to make a functional GAL4-VP16 that binds and activates the UAS promoter to induce EBFP expression (output C). (**D-F**) In the OR gate both input A and B (SP73 and SP45) induce the N7-VP16, whereas the N8-GAL4 is constitutively expressed. EBFP expression was evaluated by FACS 48h post transfection/electroporation. N=2-4 biological replicates. Statistical test: unpaired T test. p value: *<0.05; **<0.005. Created with BioRender.com.

In the AND gate configuration, we used a split synthetic transcription factor (GAL4- VP16) whereby each of the parts (VP16 and GAL4) are fused to N7 and N8 dimerizing tags respectively ^44^. VP16 is a strong transcriptional activation domain derived from the herpes simplex virus (HSV). When fused to GAL4 (forming GAL4-VP16 complex), it significantly enhances the ability of the protein to binds to UAS enhancer regions and activate gene expression. VP16-N7 was placed downstream of the NR4A2-SP, whereas the GAL4-N8 was placed downstream of the MAF-SP. Thus, both NR4A2 and MAF must be upregulated to activate the circuit and trigger the output. When reassembled, the GAL4-VP16 activates the UAS promoter triggering EBFP expression (**Fig. 5A**). The circuit showed that only in the presence of both inputs NR4A2 and MAF, EBFP levels increased both in HEK293 and Jurkat cells (**Fig. 5A-C**). The same behavior was also demonstrated in HEK293 cells using a split GFP reconstitution. Here, the GFP1-10 fragment was placed downstream NR4A2-SP, while the GFP-11 helix was fused to β-actin to improve stability and was placed downstream MAF-SP. Thus, we observed that only when both NR4A2 and MAF are expressed the split parts reassemble to form the full fluorescent GFP protein (**Supplementary Figure S5.1**).

In the OR gate configuration, we used the same split synthetic transcription factor (GAL4-VP16), with VP16-N7 placed downstream of both NR4A2- and MAF-SP, while GAL4- N8 was placed downstream of a constitutive promoter, shEF1α (**Fig. 5D-F**). Thus, the full GAL4-VP16 complex is formed when either NR4A2 or MAF is present, resulting in EBFP expression (**Fig. 5D**). We confirmed OR functions in both HEK293 and Jurkat cells, with EBFP expression as function of either of the two TFs upregulation (**Fig. 5D-F**). The design of logic gates by our synthetic sensors demonstrate that they can be exploited for programmable and precise regulation of genetic circuits with higher therapeutic potential.

Next, having observed that several SPs show a basal level of activation (**Fig. 2**), we sought to implement the CASwitch, a synthetic gene circuit based on combining two well-known network motifs: the Coherent Feed-Forward Loop (CFFL) and the Mutual Inhibition (MI), namely Coherent Inhibitory Loop (CIL). CIL was previously demonstrated to reduce leakiness while maintaining high expression, without modifying neither the TF nor the promoter^30^. The CASwitch is based on the activity of CasRx endoribonuclease that binds and cleaves the direct repeats (DR) in the 3’UTR of the mRNA output of interest causing mRNA degradation and therefore controlling the basal expression ^45^. The CasRx is regulated by a rtTA-3G-controlled pCMV/TO promoter repressed by doxycycline (Dox) (X-Y repression in **Fig. 6A**). rtTA-3G also activates the reporter gene that include the DR. Since the CasRx irreversibly binds the DR, this configuration could implement a mutual inhibition between CasRx and reporter (Y-Z in **Fig. 6A**).

**Figure 6.**
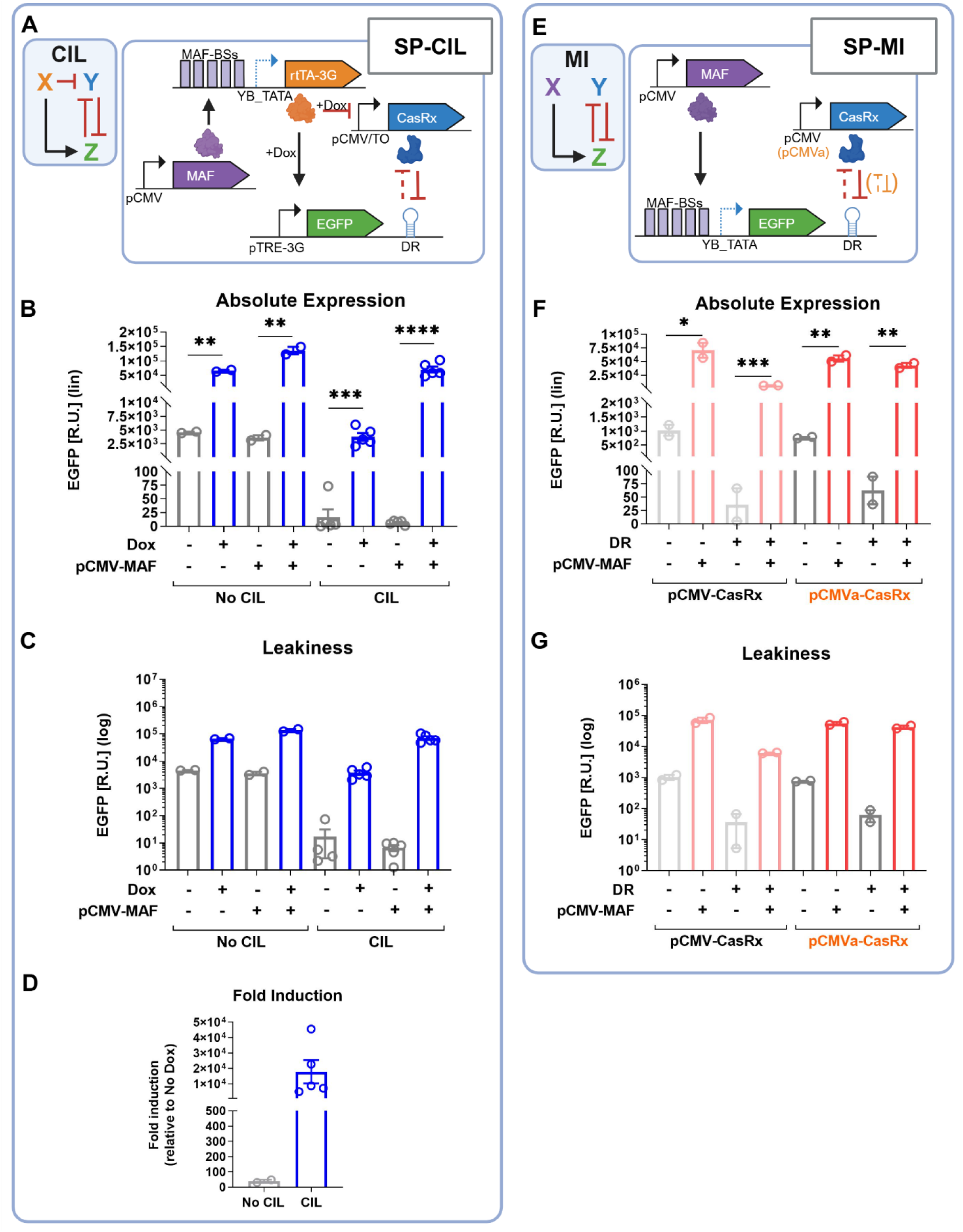
Implementation of network topologies using MAF SP. Network designs to reduce leakiness and increase fold activation by inducible promoters. (**A-D**) In the first design, MAF SP45 controls the expression of the rtTA, which activates the output in the presence of Dox, and simultaneously represses the CasRx, which is controlled by a pCMV/TO promoter (T-REx™ System, Invitrogen). CasRx in turn represses the output by irreversibly binding to direct repeats (DR) in the target mRNA, resulting in its degradation. (**E-G**) In an alternative design we restored mutual inhibition (MI) topology with SP45 driving EGFP output, while CasRx is induced by a pCMV. To tune CasRx we also developed a weaker pCMV (pCMVa). EGFP expression was quantified by FACS 48h after transfection. N=2-5 biological replicates. Statistical test: unpaired T test. p value: *<0.05; **<0.005; ***<0.0005; ****<0.0001. Created with BioRender.com.

By readapting the original architecture of the CASwitch, we developed two designs that include MAF-responsive SP45, which exhibited basal activity along with high fold induction (**Supplementary Figure S3.4A**). In the first design (SP-CIL), SP45 induces the expression of the rtTA-3G initiating the regulatory cascade (**Fig. 6A**). In the second design (SP-MI), we removed the X-Y repression to obtain a Mutual Inhibitor (MI) topology, by placing CasRx under a constitutive CMV promoter, while the reporter EGFP-DR was controlled by MAF SP45. To tune the repression by CasRx we also developed a weaker CMV promoter (pCMVa in **Fig. 6B**).

The circuit components were co-transfected in HEK293 cells in the presence or absence of the related TF, and EGFP expression was evaluated by flow cytometry 48h post transfection. As positive control we used the original system with a constitutive promoter guiding the rtTA-3G, confirming the ability of the circuit to decrease basal levels of activation generated by the TRE promoter (**Supplementary Fig. S6.1B**). When MAF SP45 was included to control rtTA-3G expression (SP-CIL design), similar reduction of the leakiness and an increase of fold induction of 459x was observed in the presence of MAF and Dox, as compared to the system without the regulation of CasRx (**Fig. 6D**).

In the SP-MI circuit, we observed high leakiness in the absence of DR, computing for a 68-fold induction when MAF was added, whereas in the presence of DR the leakiness were reduce, showing higher fold induction (650-fold) in the presence of MAF in the system (**Fig. 6F-G, Supplementary Figure S6.1C**). Moreover, by replacing the strong pCMV with a weaker version (pCMVa) obtained by adding inert spacers (2x 16bps) downstream the TATA box sequence, we tuned CasRx expression to mitigate the repression of the target EGFP mRNA, resulting in an effective compromise between basal level inhibition and activation (788-fold activation) (**Fig. 6F-G, Supplementary Figure S6.1C-D**). Overall, these results demonstrated that the synthetic sensors can be exploited to different genetic designs, and by applying rational network topology we can improve ON-OFF ratio for tight control in therapeutic interventions.

### Controlling therapeutic payload by synthetic promoters

Immunosuppressive TME can prevent the success of immunotherapies by disruption of cell-mediated and cytokine-mediated immunological functions extremely necessary for the anti-tumor immune response ^46^. To demonstrate the capacity of our SPs to induce the production of a therapeutic molecule we use as proof of concept the strongest NR4A2- responsive promoter (SP73) to guide the express of IL12p70, a cytokine that enhances T cell activation and function, and CCL21, a chemokine that promotes T cell trafficking to tumor sites47,48.

We transiently expressed in HEK293 and Jurkat cells the SP-payload construct in the presence or absence of NR4A2 and we evaluated cytokine secretion by ELISA immunoassay 48h later. Using a constitutive promoter guiding mCherry, IL-12p70 or CCL21 as controls, we show low basal levels in the mCherry negative control, and moderate expression in constitutively expressed cytokines (**Supplementary Figure S7**). SP73 cytokine control resulted in a low basal production in the absence of NR4A2 (**Fig. 7A-B**) and high release in the presence of the TF (**Fig. 7A**). We also observed CCL21 increase in HEK293 cells, but not in Jurkat, despite good electroporation efficiency. However, we observed low absolute levels of CCL21 in both cell lines in the ON state, and we speculate that technical problems in the immunoassay determined these results (**Fig. 7B**).

**Figure 7.**
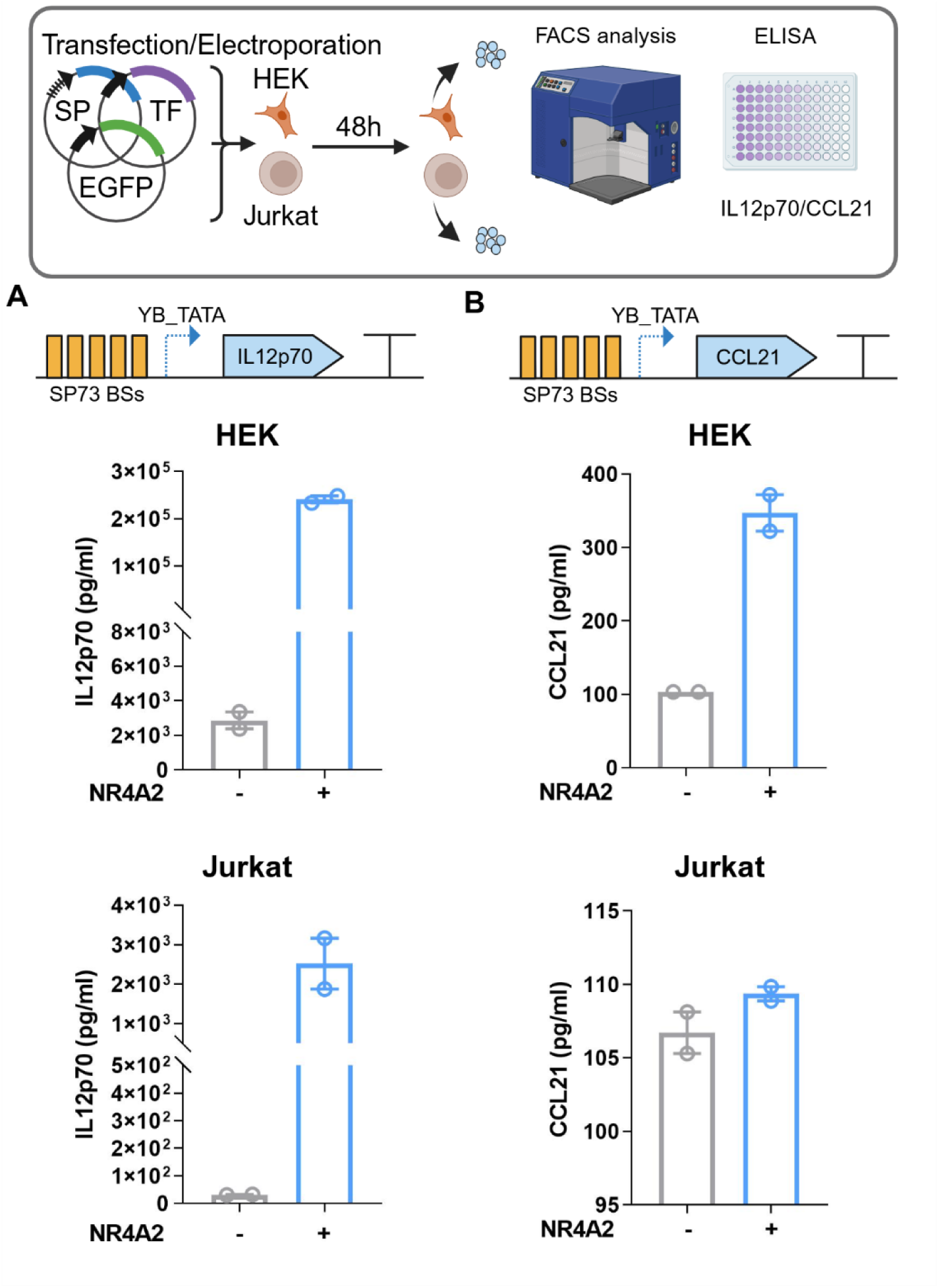
Release of therapeutic molecules by NR4A2 SP73. (**A**) The NR4A2-responsive SP73 that showed best performance was designed to trigger the expression of a cytokine IL12p70 or a chemokine CCL21. HEK293 or Jurkat cells were co-transfected/electroporated with the SP and TF plasmids along with a transfection marker, and evaluated by FACS after 48h to probe transfection efficiency. (**B**) ELISA assay were performed on supernatants to assess IL-12p70 and CCL21 production. N=2 biological replicates. Created with BioRender.com.

Overall, these data indicate that our novel synthetic promoter sensors can be engineered to deliver immunomodulatory molecules with potential to stimulate the immune system and improve immunotherapy.

### NR4A2 SP73 is activated in primary human CD8 T cells undergoing exhaustion

To evaluate the potential of SPs in therapeutic settings, we designed a lentiviral vector with the NR4A2-responsive SP73 driving the mCherry protein and engineered primary human HER2CARCD8+T cells targeting ovarian cancer cells SKOV3 (**Fig. 8A**). To correlate SP73 activation to NR4A2 upregulation, we chronically stimulated transduced T cells *ex vivo* with human CD3/CD28 antibody and evaluated classical makers of activation and exhaustion including PD1, CTLA4, LAG3 and CD39, confirming the different cellular state at the end of the assay (day 9) (**Fig. 8B**). We also assessed that over the chronic stimulation T cells lose the ability to kill SKOV3 cells (**Fig. 8C)**, suggesting the emergence of the exhaustion phenotype. Using flow cytometry we confirmed that upon exposure to antigens, NR4A2 progressively increase (**Fig. 8D)**, and the mCherry fluorescence correlates with NR4A2 levels (**Fig. 8E**), demonstrating that the SP73 responds in a specific and sensitive manner to the emergence of the dysfunctional phenotype. In perspective this proof-of concept demonstrated that our transcriptional sensors can be effectively used in ex vivo engineered T cells, opening new venues for controlled therapeutic release according to the T cell states.

**Figure 8.**
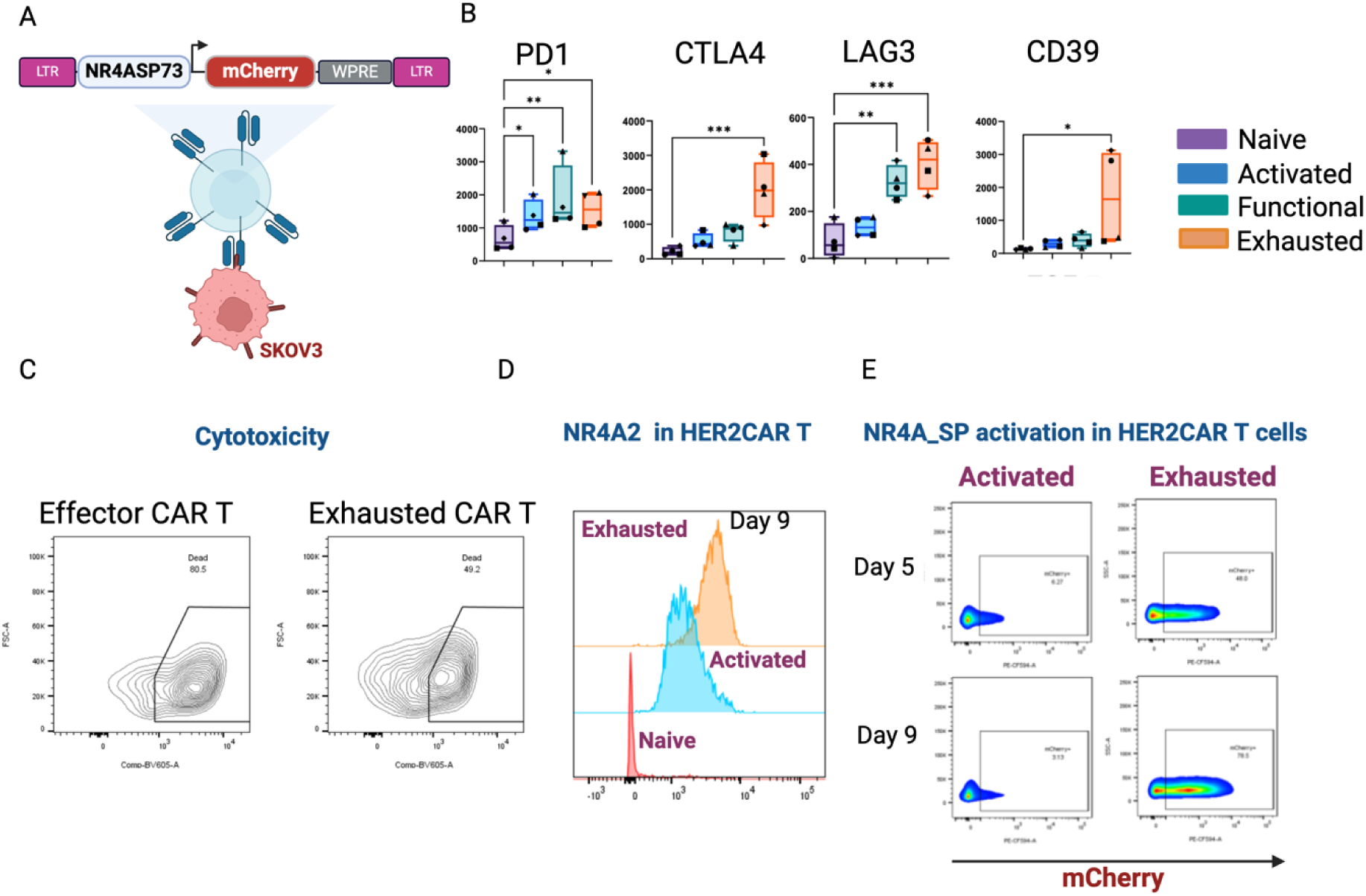
Test of NR4A2-responsive SP73 in human primary CD8 T cells. **(A)** CD8+T cells are co-transduced with SP73 driving the expression of mCherry and anti-HER2 CAR. **(B)** Engineered CD8+T cells chronically stimulated exhibit phenotypic markers typical of the emergence of exhaustion. **(C)** Engineered CD8+T cells chronically stimulated lose the cytotoxic capability when co-cultured with SKOV3 ovarian cancer cells. **(D)** NR4A2 levels progressively increase from activation to exhaustion. **(E)** NR4A2 accumulation results in the activation of the SP73 in a sensitive and specific fashion. N=5 biological replicates. Statistical test: Two-way ANOVA. p value: *<0.05; **<0,005; ***<0,0005. Created with BioRender.com.

## DISCUSSION

In this study, we show that target-directed design of synthetic promoters is a versatile approach to achieve highly specific inducible activity for T cell-specific TFs. Achieving tunable, and controlled activation of gene expression in response to differentially expressed TFs requires two main features. The first is the specificity of activation; the second is the low noise in the OFF-state. We thus focused on the deep characterization of design considerations to reduce background noise along with evaluation of best fold inductions. To ensure specificity and evaluate leakiness, we inferred for each TF the reported and predicted binding sites and engineered a simple architecture of the SPs that was meant to be compact and short. We were able to observe a response for several of the SPs engineered, whereas others did not function as expected. This may be due to unknown protein-protein interactions that serve for the transcriptional regulation and that were thus missing in our screening. This hypothesis can be investigated in the future performing Chip-seq analysis.

To investigate more sophisticated information processing, we chose two of best performing SPs (responding to MAF and NR4A2) and engineered Boolean logic gates. Moreover, we implemented an engineering design to reduce the background of leaky promoters and increase the fold activation in the ON-state (e.g. SP45 responding to MAF). Our modular design approach facilitates straightforward optimization, paving the way for future clinical translation and the targeting of additional dysfunctional cell types.

The major advantage of creating for each TF a library of SPs based on different BSs is the remarkable diverse strength of activation offering a tool for precise and programmable signal adaptation, crucial for advancing gene circuit design with finely tuned control over cellular behavior.

Several challenges must be addressed to progress this approach toward therapeutic applications. Circuit design and testing will likely require high-throughput cloning, automation steps, and machine learning (ML) analysis for quicker evaluation of a higher number of sensors candidates. These approaches are being addressed in another studies ^49,50^. We also attempted a ML-based approach to infer the parameters that affect promoter strength. Our neural network-based modeling approach, trained on the EPD dataset, achieved a moderate correlation (r = 0.49), which was a good first result considering the limited number of data provided, and the fact that the reference dataset originates from yeast promoters. Together, these findings validate our rational design strategy and modeling framework for engineering SPs to optimize therapeutic output and improve immunotherapy.

We finally show first tests in human primary CD8 T cells of SP functionality. Further screenings are currently ongoing to understand the correlation with results obtained in Jurkat cells, a T cell line widely used in pre-screening phases. Indeed, our data strongly indicate the cell-context affect the functionality and general parameters of the SPs.

Our approach, which allows cell context-specific secretion of immunomodulators, offers a valuable framework for investigating the immunological mechanisms underlying the tumor microenvironment. Synthetic gene circuits have demonstrated potential for treating immunological disorders by detecting extracellular disease markers ^51^. Similarly, our strategy of sensing internal cell states using multiple artificial promoters could be adapted to address other dysfunctional cells. With straightforward modifications to the synthetic promoter inputs and immunomodulatory outputs, this platform enables precise and multifactorial programming of immunological functions.

In summary, our findings represent a significant advancement in synthetic biology. By engineering Synthetic Promoters that respond to specific transcription factors, we provide aflexible and powerful platform for controlling cellular behavior, providing exciting prospects for novel therapeutics aimed at tuning T cell exhaustion and optimizing therapeutic outcomes.

## MATERIAL AND METHODS

### Transcription Factors (TFs) and TF-Binding Sites (TF-BSs) selection

A universal epigenetic and transcriptional gene signature of T cell dysfunction in mouse and human has been identified by exploring deposited gene expression data (GSE41867, GSE86042) and ATAC-Seq data (GSE164978 GSE97646 GSE86881); the analysis yield 2844 transcripts of genes, up-regulated in dysfunctional vs effector or memory CD8+ cells in chronic infections and cancer, confirmed at chromatin level. By categorizing all these genes, we identified 197 TFs, whose expression has been further investigated in T cell and CAR-T cells datasets of exhaustion (GSE240851, GSE179613, GSE160174, and https://dice-database.org). By ranking the most relevant TFs in these datasets, we selected a subset of seven TFs (IRF4, BATF, MAF, GATA3, NR4A2, EOMES and IKZF2) to sense by designing novel Synthetic Promoters (SPs). The TFs selected were based on their expression in the early stages of activation, functional effector/memory and in dysfunctional cells, and on their well characterized role in promoting or augmenting dysfunctional states ^22–24,27,35^. To design SPs, we selected transcription factors binding sites (TF-BSs) from literature and TFBS Data Bank (TFBS-DB) web tool (http://tfbsdb.systemsbiology.net/) ^52^ which use robust ATACseq data and motif repositories such as JASPAR, SELEX and TRANSFAC.

### Plasmids library generation

<u>Design and Generation of Synthetic Promoters for Screening</u> - The SPs are composed of three main modules: the TF-BSs, a minimal core promoter (minP) and a reporter gene. TF- BSs were cloned in 5x tandem repetition, upstream the minP module, alternating each TF- BSs with spacers of 10-20bps. We combined TF-BSs with four different minP (YB-TATA, miniCMV, miniTK, LateADEp). Constructs for each minP where no TF-BSs were inserted was also created as minP basal activity control. A mCherry fluorescent protein was inserted downstream the SPs as output.

These modules composing the SP and reporter gene were cloned into pAI274 plasmid backbone (pT2A-MCS-pPGK-H2B-mCherry-pA). Originally, the AI274 plasmid encodes for nuclear mCherry under the regulation of the constitutive promoter PGK followed by SV40 polyA signal. We switched the PGK and the H2B sequences from the AI274 backbone with the synthetic promoter sequence. Each module was flanked by two restriction sites, allowing the easy combination of all parts (NotI-5xTF-BSs+spacers-HindIII-minP-PacI). Each module was assembled via oligo annealing reactions and ligated into the linearized vector. First, constructs control for each minP (no TF-BSs) were created. From this last, all the others SPs were created by replacing the TF-BSs region via the NotI and HindIII sites with new TF-BSs by digestion-oligo annealing-ligation. All general structures and sequences are listed in Supplementary Tables S2A-B, S3 and S4.

<u>Generation of SPs Driving Circuits and Payload Release</u> - The circuits designed for Boolean Logic Gates, Feed-Forward Loop and payload release were created by implementing the combinatorial modular design MTK (Mammalian Tool Kit, ^53^), a Golden Gate-based cloning toolkit designed for the versatile, reversible and fast assembly of DNA vectors and their implementation in mammalian models. Modular parts can be assembled into transcriptional units and further integrated into complex circuits ^54^. To adapt the SPs to the MTK, we split the promoter into two different parts: MTK2a (5xTF-BSs+spacers) and 2b (minP YB_TATA), keeping as much as possible original sequences and adapting it to match the original MTK overhangs. Some parts of the MTK library were ordered from Addgene (https://www.addgene.org/browse/article/28197510/) or generated from oligos and PCR fragments. Domestication of new parts was done by Esp3I Golden Gate assembly into a chloramphenicol resistant destination vector and by selection of white colonies. From the library of parts, transcriptional units (TUs) were built by assembling Parts (1-5) with a BsaI-HF Golden Gate reaction and selection of ampicillin resistant white colonies, which generates plasmids ready for introduction into mammalian cells. MTK2a parts from MAF-SP45 and NR4A2-SP73 were created and combined to other modules in order to create new SP constructs.

All DNA fragments were either chemically synthesized or PCR-amplified from existing plasmids. All oligos or DNA syntheses were performed by Sigma-Aldrich or Integrated DNA Technologies (Coralville, IA), respectively. Plasmids were sequence-verified by Sanger Sequencing (Genewiz, Germany). Since each TF can recognize different DNA responsive elements, by using this combinatorial modular design we generated a library of over than 80 SPs sensors (**Supplementary Tables S2A and B**). Plasmids for transfection and electroporation were prepared by high-quality standard procedures and purified using a Plasmid Midi/Maxiprep Kit (Qiagen) according to manufacturer’s protocol.

### Cell lines maintenance

HEK 293FT and Jurkat Clone E-6 cells were obtained from ATCC (Manassas, VA). HEK 293FT cells were cultured in high-glucose DMEM (Life Technologies, Grand Island, NY) and Jurkat cells were maintained in RPMI1640 (Life Technologies). All media were supplemented with 10% FBS (Life Technologies), 1% penicillin/streptomycin (Gibco), 2mM L-Glutamine (Sigma-Aldrich) and 1% Non-Essential Amino Acids (HyClone). Cells were incubated at 37°C under conditions of 100% humidity and 5% CO2.

### Cell transfection and electroporation

Transfections in HEK 293FT cells were performed in a 24-well plate for flow cytometry analysis. 7×10^5^ cells per well were seeded approximately 24 h before transfection in complete media. Cells were transfected with PEI transfection reagent according to manufacturer’s instructions using 200-400 ng DNA diluted in Opti-MEM reduced serum media (Gibco) before being mixed and incubated for 15 minutes prior to addition to the cells. Cells were detached and prepared for flow cytometry analysis 48h after transfection. For SP screening, the synthetic promoters were transfected together with an expression plasmids encoding for the SP specific TF plus a plasmid encoding for GFP using the following concentrations: 100 ng pSP, 50 ng pCMV-TF, 20ng pCMV-EGFP. Jurkat cells were electroporated in a 96-well plate with Neon Transfection System 10 μl Neon Tip (Life Technologies). A total of 1.3 μg of DNA was prepared in a 0.5 mL tube. Meanwhile 5×10^4^ cells/well were centrifuged in PBS at 200xg for 5 min at room temperature. Cells were suspended in suspension buffer R, added to the DNA tube and gently mixed. The DNA and cell mixtures were picked with the appropriate Neon Tip, electroporated (pulse voltage 1422 v, pulse width 10 ms, pulse number 3) and cultured in RPMI medium without antibiotics. Cells were collected and prepared for flow cytometry analysis 48h after transfection.

### Flow Cytometry

All cells were analyzed with a BD CELESTA™ cell analyzer (BD Biosciences) 48h after transfection/electroporation. HEK 293FT cells were detached with 50 μl of Trypsin and resuspended in 70 μl of PBS, while Jurkat cells were analyzed directly. For each sample, >10000 singlet events were collected, a compensation matrix created using unstained cells (wild type) and single-color controls (EBFP only, EGFP only and mCherry only). Population of live cells and single cells were selected according to FSC/SSC parameters, followed by a gate strategy selecting the fluorescent cells sequentially. Data containing percentage and Geometric Mean were extrapolated from this analysis and used for calculation of SP basal and inducible activity. Data analysis was performed with FlowJo™ v10.9.0 Software (BD Life Sciences).

### Basal activity, absolute and fold induction calculation of SPs characterization

Background activity of SPs were represented by extrapolating mCherry Geometric Mean Fluorescence Intensity (gMFI) after sequential gate analysis (FSC/SSC>>Single cells>>EGFP+ cells>>mCherry+ cells) performed in FlowJo™. Absolute induction of SPs were represented by a normalization between reporter mCherry and transfection control EGFP, calculated for each condition using the following formula:

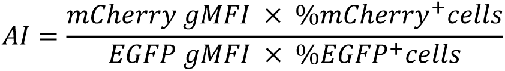

Here, mCherry gMFI is extracted from double EGFP+mCherry+ cell population, %mCherry+ cells is extracted from EGFP+ population, EGFP gMFI is extracted from single EGFP+ population and %EGFP+ cells is extracted from Single cells. The calculation of Fold-induction was calculated by dividing the AI of inducible conditions (+ TF) by non-inducible (no TF).

### ELISA assays

ELISAs were performed with the Human IL12p70 Lumit (Promega) or Human Secondary Lymphoid-tissue Chemokine (CCL21) (SLC) Uncoated ELISA Kit (ThermoFisher Scientifics). Supernatant were harvested from 24 well plates with transfected HEK 293FT or electroporated Jurkat cells 48 hours after transfection/electroporation and IL12p70 or CCL21 was quantified according to manufacturer instructions.

### Primary human CD8 T cells isolation and transduction

Human CD8^+^ T cells were isolated from buffy coats of blood donors by magnetic sorting, activated with Dynabeads™ Human T-Activator CD3/CD28 at 2:1 beads:cells ratio, and cultured in R10 medium in the presence of IL-2 (30 U/ml). Lentivirus transduction were carried out in 6-well plates in stimulated T cells after 24-48h from stimulus. Cells were co-transduced with lentivirus for SPs and CAR-Her2 at a MOI of 5 and 15, respectively, and kept at a concentration of one million cells/ml by adding new fresh medium plus IL-7 and IL-15 (1:1000) to allow expansion for 10-14 days.

### Immunostaining and Cytotoxicity Assay

For experiments in T cells, CD8+ T cells were collected at specific time points, washed twice with PBS and then incubated with LIVE/DEAD™ Fixable Yellow Dead Cell Stain (L34959, Life Technologies Inc.) for 20 min at RT for dead cell exclusion. CD8+ T cells were then washed twice with PBS and stained for 20 min at RT with the following antibodies: anti- PD1 (2249750, Sony Biotechnology Inc.), anti-LAG3 (565718, BD Horizon) and anti-CD39 (2241070, Sony Biotechnology Inc.). Cells were fixed and permeabilized using the Transcription factor buffer set (562574, BD Pharmingen Inc.), following the manufacturer’s instruction, then stained for 30 min at 4°C with anti- CTLA4 (555854, BD Pharmingen Inc.) and anti-NR4A2 (orb464231, Biorbyt ltd). Stained cells were then acquired by FACS as described above.

For the cytotoxicity assay, functional and exhausted CAR-Her2-CD8+ T cells were co-cultured for 24h in a 3:1 ratio with SKOV3 (Her2+) ovarian cancer cells, previously stained with cell tracker (C2927, Thermo Fisher Scientific). Cells were then collected and incubated with LIVE/DEAD™ Fixable Yellow Dead Cell Stain (L34959, Life Technologies Inc.) for 20 min at RT for live/dead cells discrimination. The percentage of dead SKOV3 in each condition was determined by FACS.

### Transfer Learning and Promoter Ranking

Detailed description can be found in **Supplementary Note 1**.

## Statistical Analysis

Each experiment was repeated independently at least three times with similar results. To compare multiple connected conditions, statistical analysis was performed using one-way ANOVA with Tukey’s multiple comparison analysis. For comparisons between two conditions, a one-tailed unpaired t-test was used. The threshold for significance was set to 0.05 for the global ANOVA and t-test p-values.

## Acknowledgments

Author contributions: V.S. conceived the project and coordinated research activities. VS and GB designed experiments. GB performed experiments and analysis. GM designed and tested Boolean logic gates. AS and AR performed bioinformatics analysis. GB, GM and AS designed in vitro experiments for T cell exhaustion and performed the assay. GB, FC, DP, FT, LM designed SPs, FM, FS, NE developed the transfer learning model. GB and VS wrote the manuscript. GM, AS, FM, provided input and edited the manuscript.

## FUNDING

ERC Starting grant Synthetic T-rex [852012]; NextGenerationEU PNRR MUR [M4C2]; National Center for Gene Therapy and Drugs based on RNA Technology [CN00000041].

## Conflict of interest statement

IP have been filed for the MAF and NR4A2 SPs.

## Notes

### Competing Interest Statement

The authors have declared no competing interest.

