## Supplementary Figures for "Design of novel synthetic promoters to tune gene expression in T cells"

### FIGURE'S CAPTION

**Supplementary Figure S1. Bioinformatics selection of transcription factors (TFs) differentially expressed in activated and exhausted T cells.** Different studies utilized to select the TFs showed in individual heat maps.

**Supplementary Figure S2.1. Background noise of the SPs in the OFF state.** (**A**) General experimental workflow of the screening process using the SPs library. SP sensors were characterized by transient transfection or electroporation in HEK293 and Jurkat cell lines, respectively. Three constructs are introduced into the cells: the SP, the cognate TF, and a constitutively expressed EGFP as transfection control. After 48 hours, we evaluated by flow cytometry (FACS) the expression of the mCherry fluorescent reporter. A scramble control in which the TF-BSs were not inserted were used to check basal activities of the minimal promoter. (**B**) Leakiness of miniP combined to TF-BS of IRF4/BATF reported by mCherry gMFI 48h post-transfection/electroporation in HEK293 and Jurkat cells. (**C**) Leakiness of miniP combined to TF-BS of MAF, GATA3, reported by mCherry gMFI 48h post-transfection/electroporation in HEK293 and Jurkat cells. N=2 biological replicates. Created with BioRender.com.

**Supplementary Figure S2.2. Background noise of the SPs in the OFF state.** (**A**) Leakiness Heat Map of YB_TATA combined to TF-BS of NR4A2, EOMES, IKZF2, reported by mCherry gMFI 48h post-transfection/electroporation in HEK293 and Jurkat cells. (**B-D**) Leakiness of YB_TATA minP combined to TF-BS of NR4A2, EOMES, IKZF2 reported by mCherry gMFI 48h post-transfection/electroporation in HEK293 and Jurkat cells. §: SPs containing less than 5x TF-BSs repetition. N=2 biological replicates. Created with BioRender.com.

**Supplementary Figure S3.1. Gate strategy analysis performed to characterize the SP activation.** (**A**) Gate strategy hierarchy used for transfected HEK293 cells. (**B**) Gate strategy hierarchy used for electroporated Jurkat cells.

**Supplementary Figure S3.2. Evaluation of IRF4/BATF YB_TATA SPs activation.** (**A**) Absolute induction of SPs showing IRF4/BATF SPs in YB_TATA configuration in HEK293 or (**B**) Jurkat cells. N=2 biological replicates. Statistical test: one-way ANOVA with Tukey's multiple comparison test (for IRF4/BATF SPs). p value: *<0.05; **<0.005; ***<0.0005; ****<0.0001. Hatched bars: SPs containing less than 5xTF-BSs. Created with BioRender.com.

**Supplementary Figure S3.3. Evaluation of IRF4/BATF miniCMV SPs activation.** (**A**) Absolute induction of SPs showing IRF4/BATF SPs in miniCMV configuration in HEK293 or (**B**) Jurkat cells. N=2 biological replicates. Statistical test: one-way ANOVA with Tukey's multiple comparison test (for IRF4/BATF SPs). p value: *<0.05; **<0.005; ***<0.0005; ****<0.0001. Hatched bars: SPs containing less than 5xTF-BSs. Created with BioRender.com.

**Supplementary Figure S3.4. Evaluation of IRF4/BATF miniTK SPs activation.** (**A**) Absolute induction of SPs showing IRF4/BATF SPs in miniTK configuration in HEK293 or (**B**) Jurkat cells. N=2 biological replicates. Statistical test: one-way ANOVA with Tukey's multiple comparison test (for IRF4/BATF SPs). p value: *<0.05; **<0.005; ***<0.0005; ****<0.0001. Hatched bars: SPs containing less than 5xTF-BSs. Created with BioRender.com.

**Supplementary Figure S3.5. Evaluation of IRF4/BATF lateADEp SPs activation.** (**A**) Absolute induction of SPs showing IRF4/BATF SPs in lateADEp configuration in HEK293 or (**B**) Jurkat cells. N=2 biological replicates. Statistical test: one-way ANOVA with Tukey's multiple comparison test (for IRF4/BATF SPs). p value: *<0.05; **<0.005; ***<0.0005; ****<0.0001. Hatched bars: SPs containing less than 5xTF-BSs. Created with BioRender.com.

**Supplementary Figure S3.6. Evaluation of MAF and GATA3 SPs activation.** (**A**) Absolute induction of SPs showing MAF SPs in all minP configuration in HEK293 and Jurkat cells. (**B**) Absolute induction of SPs showing GATA3 SPs in all minP configuration in HEK293 and Jurkat cells. N=2 biological replicates. Statistical test:  unpaired T test. p value: *<0.05; **<0.005; ***<0.0005; ****<0.0001. Hatched bars: SPs containing less than 5xTF-BSs. Created with BioRender.com.

**Supplementary Figure S3.7. Evaluation of NR4A2, EOMES and IKZF2 SPs activation.** (**A**) Absolute induction of SPs showing NR4A2 SPs in YB_TATA minP configuration in HEK293 and Jurkat cells. (**B**) Absolute induction of SPs showing EOMES SPs in YB_TATA minP configuration in HEK293 and Jurkat cells. (**C**) Absolute induction of SPs showing IKZF2 SPs in YB_TATA minP configuration in HEK293 and Jurkat cells. N=2 biological replicates. Statistical test:  unpaired T test. p value: *<0.05; **<0.005; ***<0.0005; ****<0.0001. Hatched bars: SPs containing less than 5xTF-BSs. Created with BioRender.com.

**Supplementary Figure S3.8. Evaluation of IRF4/BATF YB_TATA SPs activation by fold induction in bar graphs.** (**A**) Fold Induction of SPs showing IRF4/BATF SPs in YB_TATA configuration in HEK293 or (**B**) Jurkat cells. N=2 biological replicates. Hatched bars: SPs containing less than 5xTF-BSs. Created with BioRender.com.

**Supplementary Figure S3.9. Evaluation of IRF4/BATF miniCMV SPs activation by fold induction in bar graphs.** (**A**) Fold Induction of SPs showing IRF4/BATF SPs in miniCMV configuration in HEK293 or (**B**) Jurkat cells. N=2 biological replicates. Hatched bars: SPs containing less than 5xTF-BSs. Created with BioRender.com.

**Supplementary Figure S3.10. Evaluation of IRF4/BATF miniTK SPs activation by fold induction in bar graphs.** (**A**) Fold Induction of SPs showing IRF4/BATF SPs in miniTK configuration in HEK293 or (**B**) Jurkat cells. N=2 biological replicates. Hatched bars: SPs containing less than 5xTF-BSs. Created with BioRender.com.

**Supplementary Figure S3.11. Evaluation of IRF4/BATF lateADEp SPs activation by fold induction in bar graphs.** (**A**) Fold Induction of SPs showing IRF4/BATF SPs in lateADEp configuration in HEK293 or (**B**) Jurkat cells. N=2 biological replicates. Hatched bars: SPs containing less than 5xTF-BSs. Created with BioRender.com.

**Supplementary Figure S3.12. Evaluation of MAF, GATA3, NR4A2, EOMES and IKZF2 SPs activation by fold induction in bar graphs.** (**A-B**) Fold Induction of SPs showing MAF or GATA3 SPs in all minP configuration in HEK293 and Jurkat cells. (**C-E**) Fold Induction of SPs showing NR4A2, EOMES or IKZF2 SPs in YB_TATA minP configuration in HEK293 and Jurkat cells. N=2 biological replicates. Hatched bars: SPs containing less than 5xTF-BSs. Created with BioRender.com.

**Supplementary Figure S5. Implementation of AND Boolean Logic Gates using MAF and NR4A2 SPs for split GFP.** AND gate was composed by a split GFP. GFP1-10 is controlled by NR4A2-responsive SP73 (input A) whereas GFP11-βactin is induced by MAF-responsive SP45 (input B). When both TFs are present, the two halves reconstitute to make a fluorescent GFP (output C). GFP expression was evaluated by FACS 48h post transfection in HEK293 cells. N=3 biological replicates. Statistical test:  unpaired T test. p value: *<0.05; **<0.005. Created with BioRender.com.

**Supplementary Figure S6.1. Implementation of network topologies using MAF SP.** Network designs to reduce leakiness and increase fold activation by inducible promoters. (**A**) Original design and tests for the CASwitch system. (**B**) Histogram showing MAF SP45 basal activity (no TF) and activation (+MAF). (**C**) Fold Induction of configuration SP-MI. (**D**) Dot-plots of mScarlet expression for configuration SP-MI showing decreased level of expression in engineered altered pCMV promoter (pCMVa) which correlates with CasRx expression as it is connected by a P2A sequence (CasRx::P2A::mScarlet). Created with BioRender.com.

**Supplementary Figure S6.2. Implementation of network topologies using MAF SP.** Network designs to reduce leakiness and increase fold activation by inducible promoters. (**A**) Dot-plots of EGFP reporter gene expression for configuration SP-CIL showing decreased level of leakiness in the presence of CasRx (+CIL). (**B**) Dot-plots of EGFP reporter gene expression for configuration SP-MI showing decreased level of leakiness and increased level of expression in the presence of DR domain and using altered pCMV promoter (pCMVa).

**Supplementary Figure S7. Release of therapeutic molecules by NR4A2 SP73.** ELISA control conditions for NR4A2-responsive SP73 designed to trigger the expression of a (**A**) cytokine IL12p70 or (**B**) a chemokine CCL21 in HEK293 or Jurkat cells when co-transfected/electroporated with constitutive promoter guiding expression of IL12p70, CCL21 or a fluorescent protein (mCherry), along with a transfection marker. Supernatants were evaluated by ELISA assay after 48h of transfection/electroporation. N=2 biological replicates. Created with BioRender.com.

### SUPPLEMENTARY FIGURES

#### Supplementary Figure S1

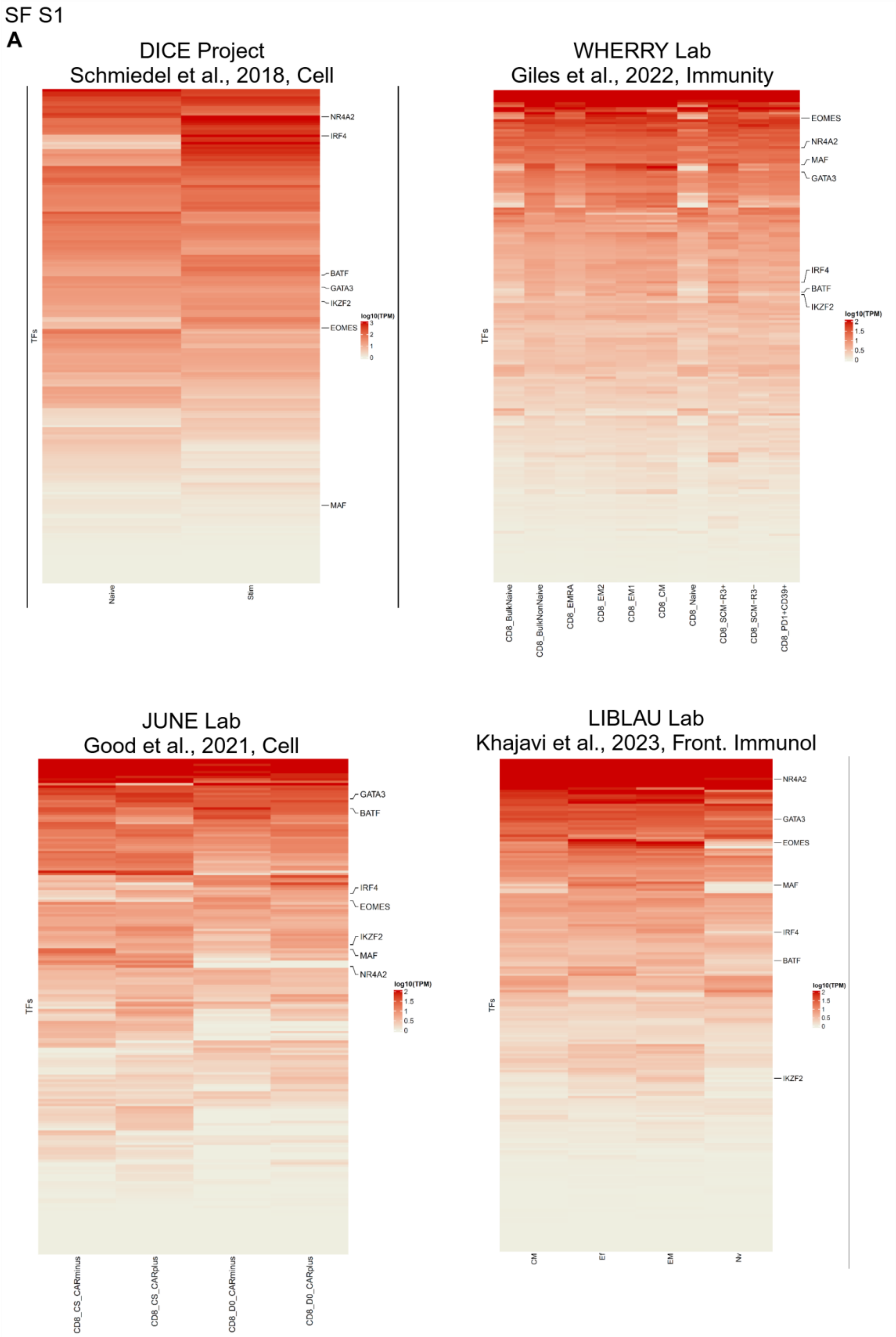

### Supplementary Figure S2.1

SF S2.1 - Synthetic Sensor Characterization

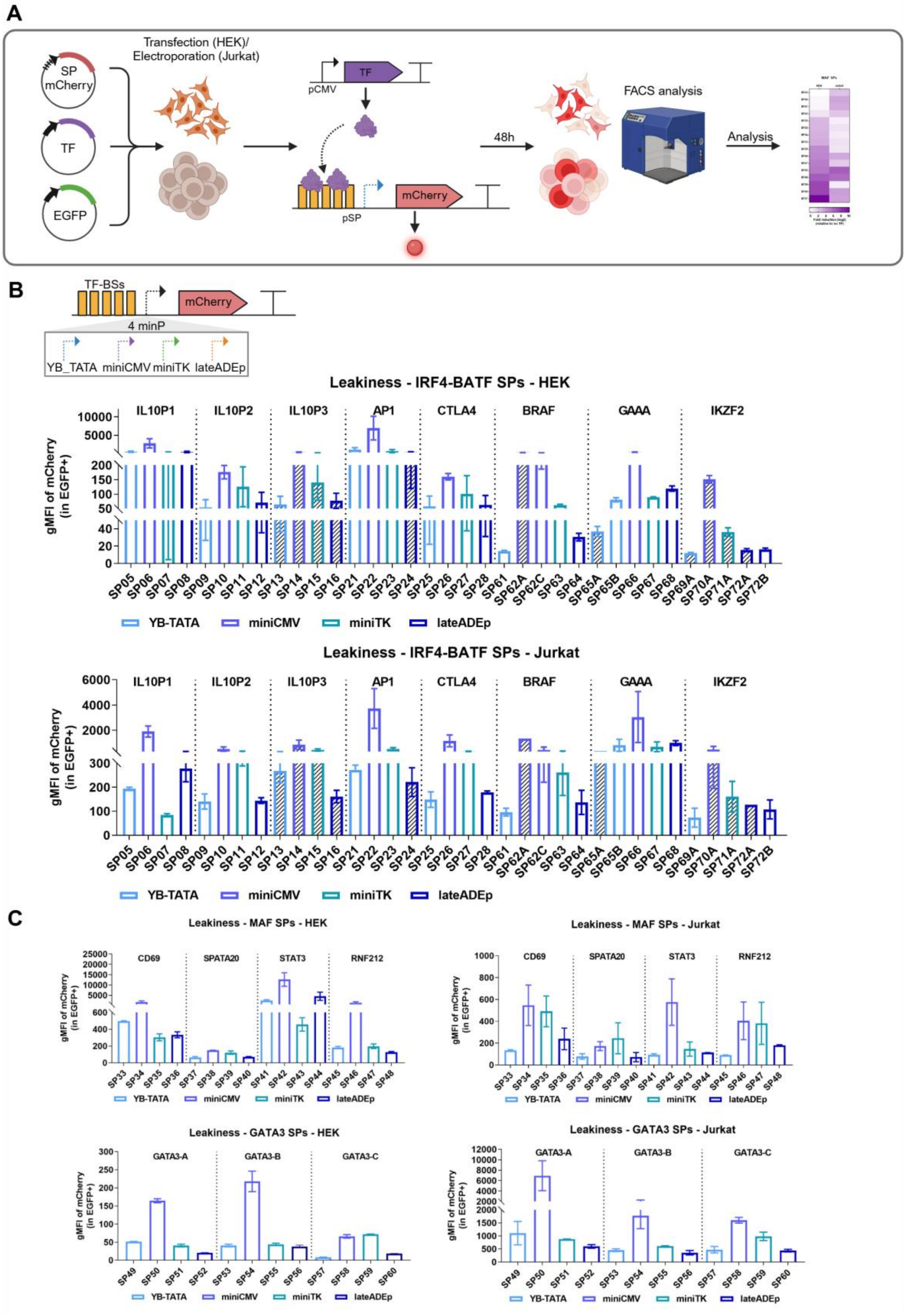

### Supplementary Figure S2.2

SF S2.2 - Synthetic Sensor Characterization

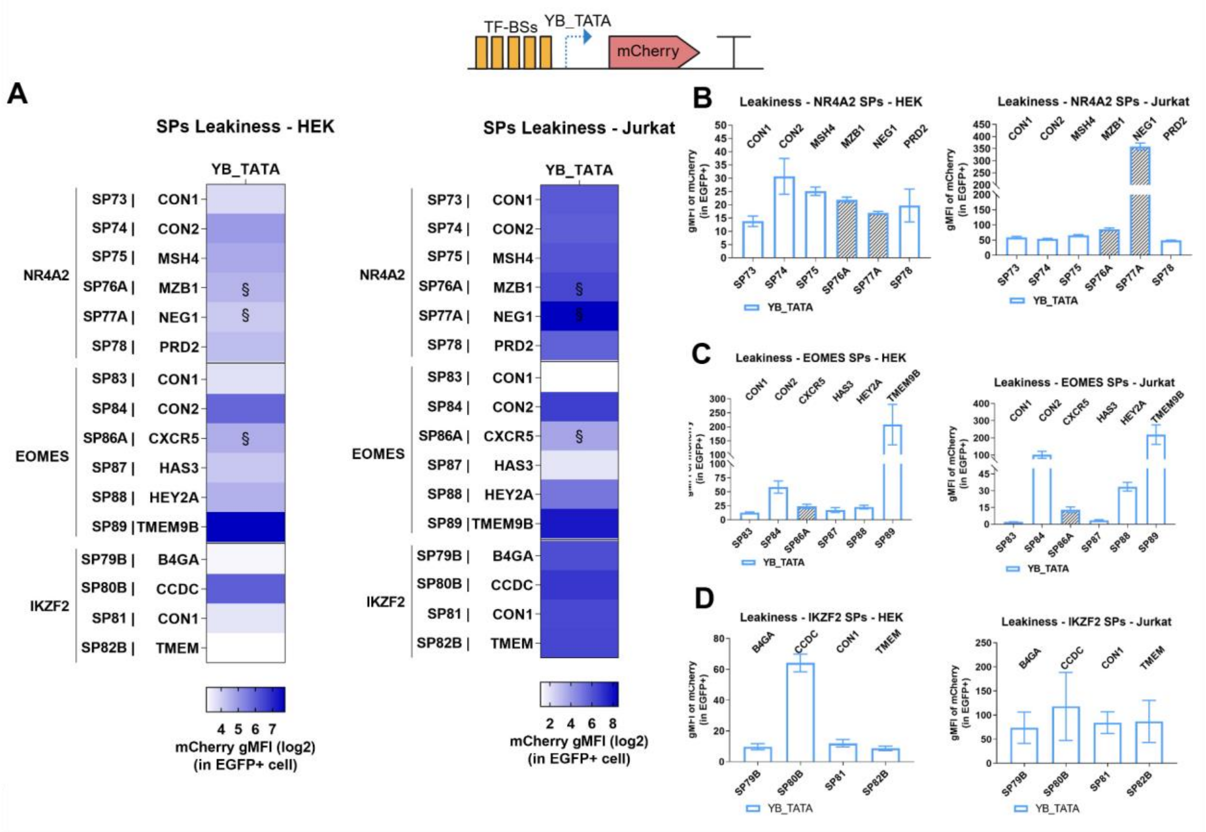

### Supplementary Figure S3.1

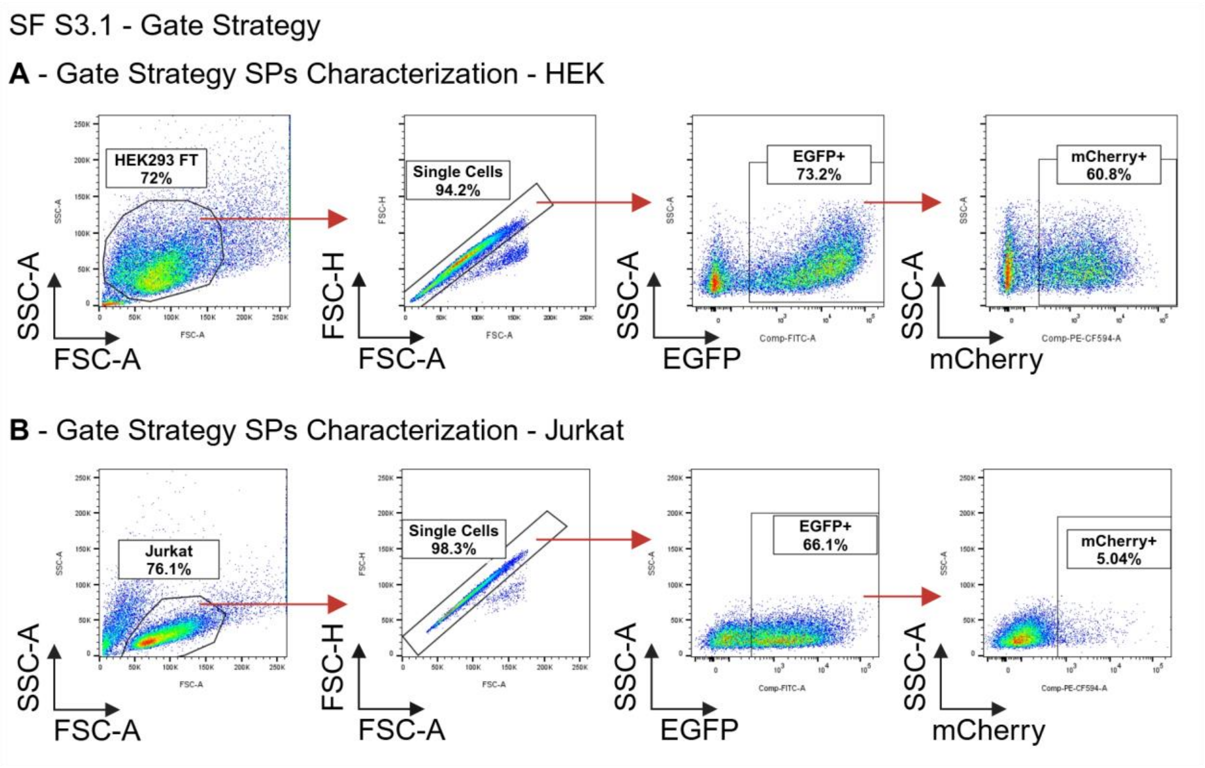

**Supplementary Figure S3.2**

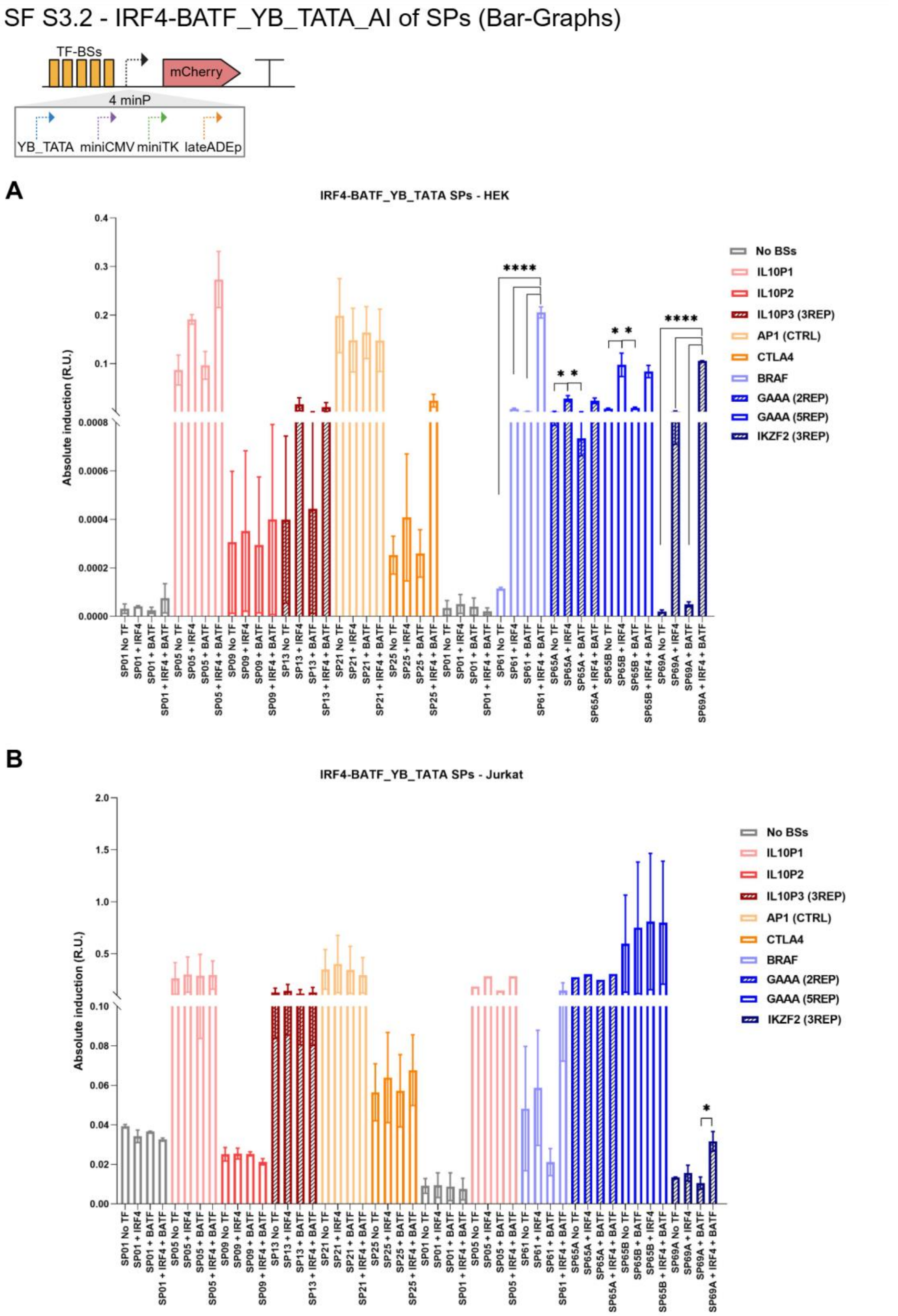

### Supplementary Figure S3.3

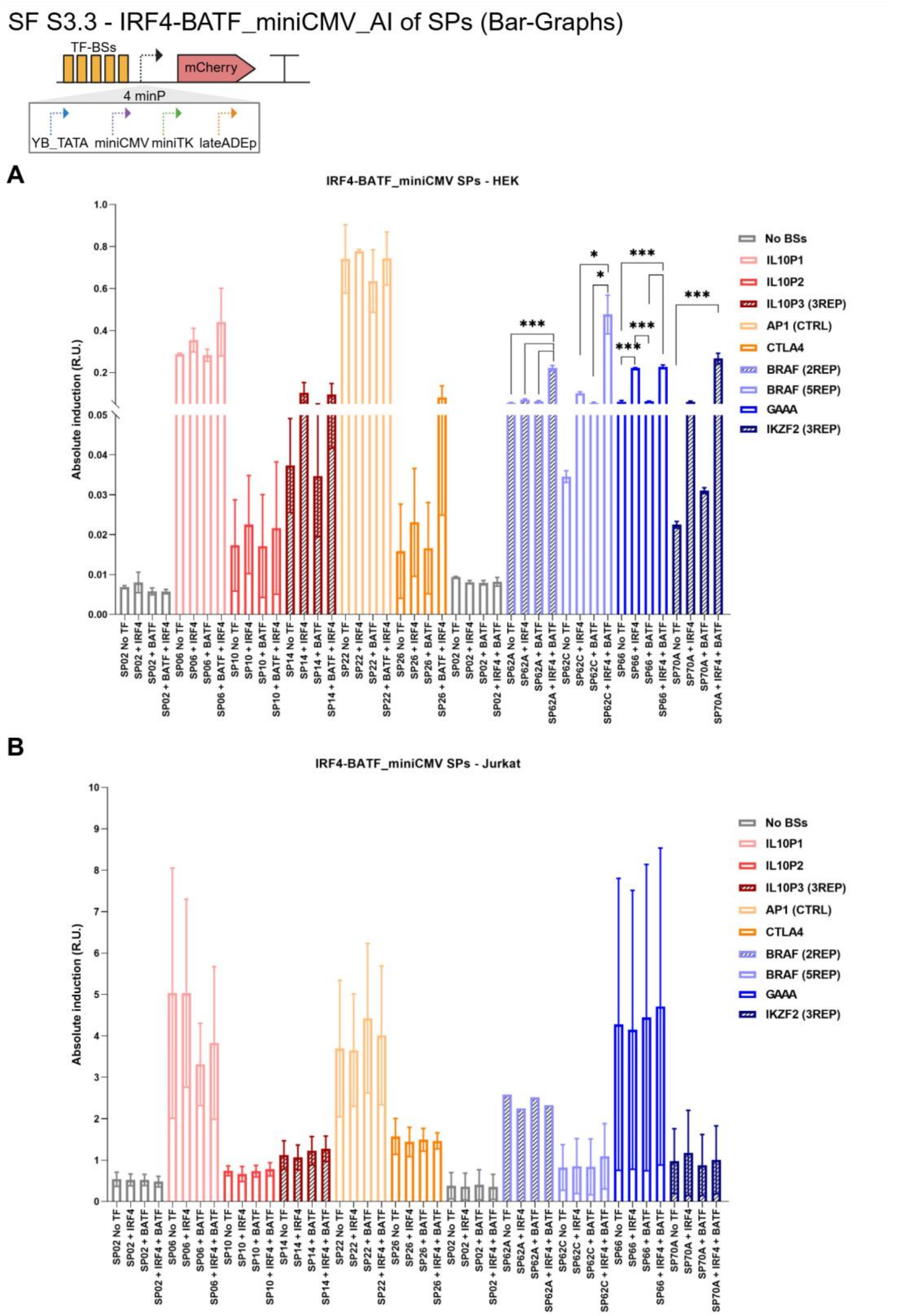

**Supplementary Figure S3.4**

SF S3.4_IRF4-BATF_miniTK_AI of SPs (Bar-Graphs)

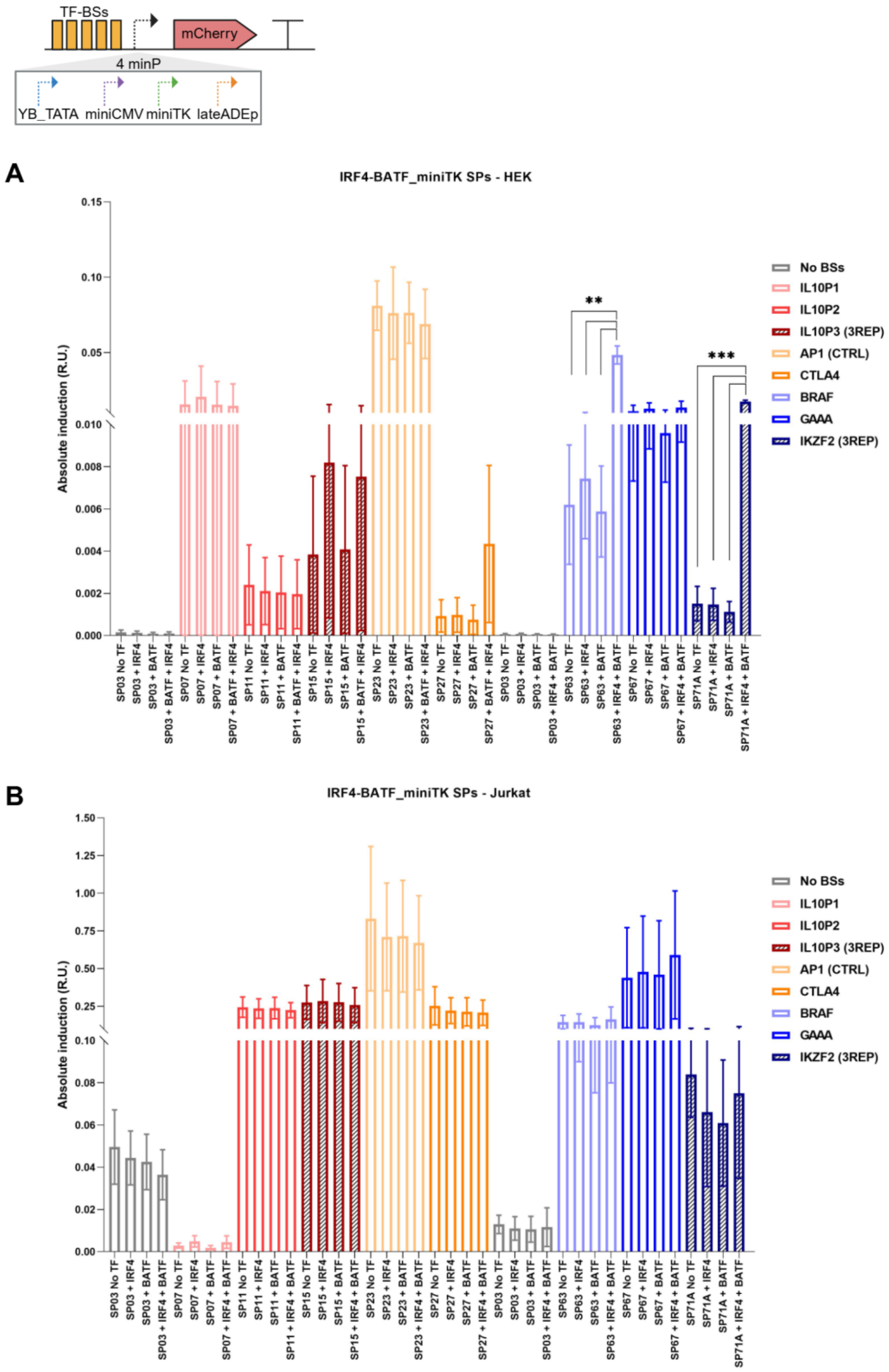

**Supplementary Figure S3.5**

SF S3.5_IRF4-BATF_lateADEp_AI of SPs (Bar-Graphs)

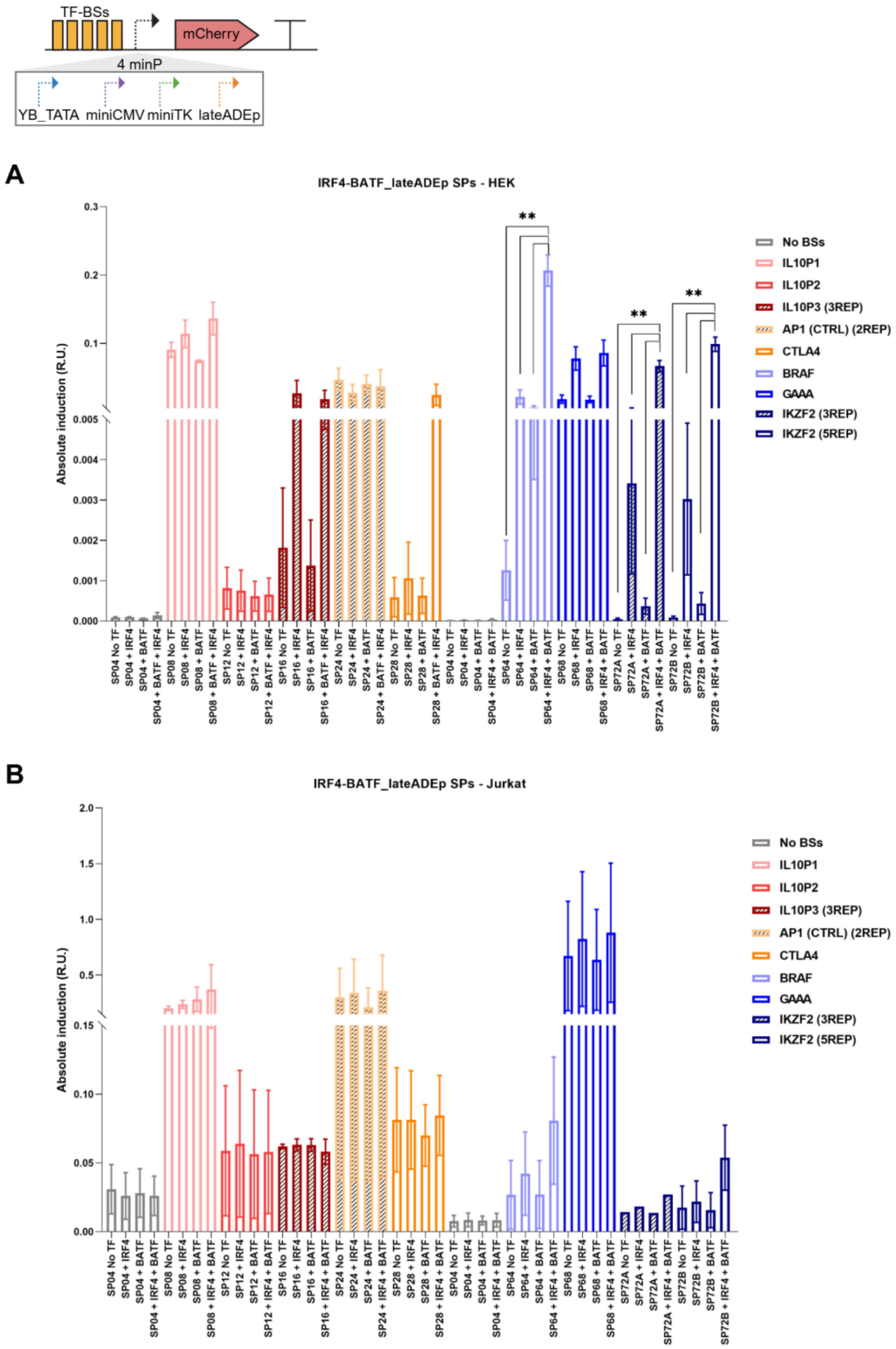

**Supplementary Figure S3.6**

SF S3.6 - Absolute Induction Profile of SPs (Bar-Graphs)

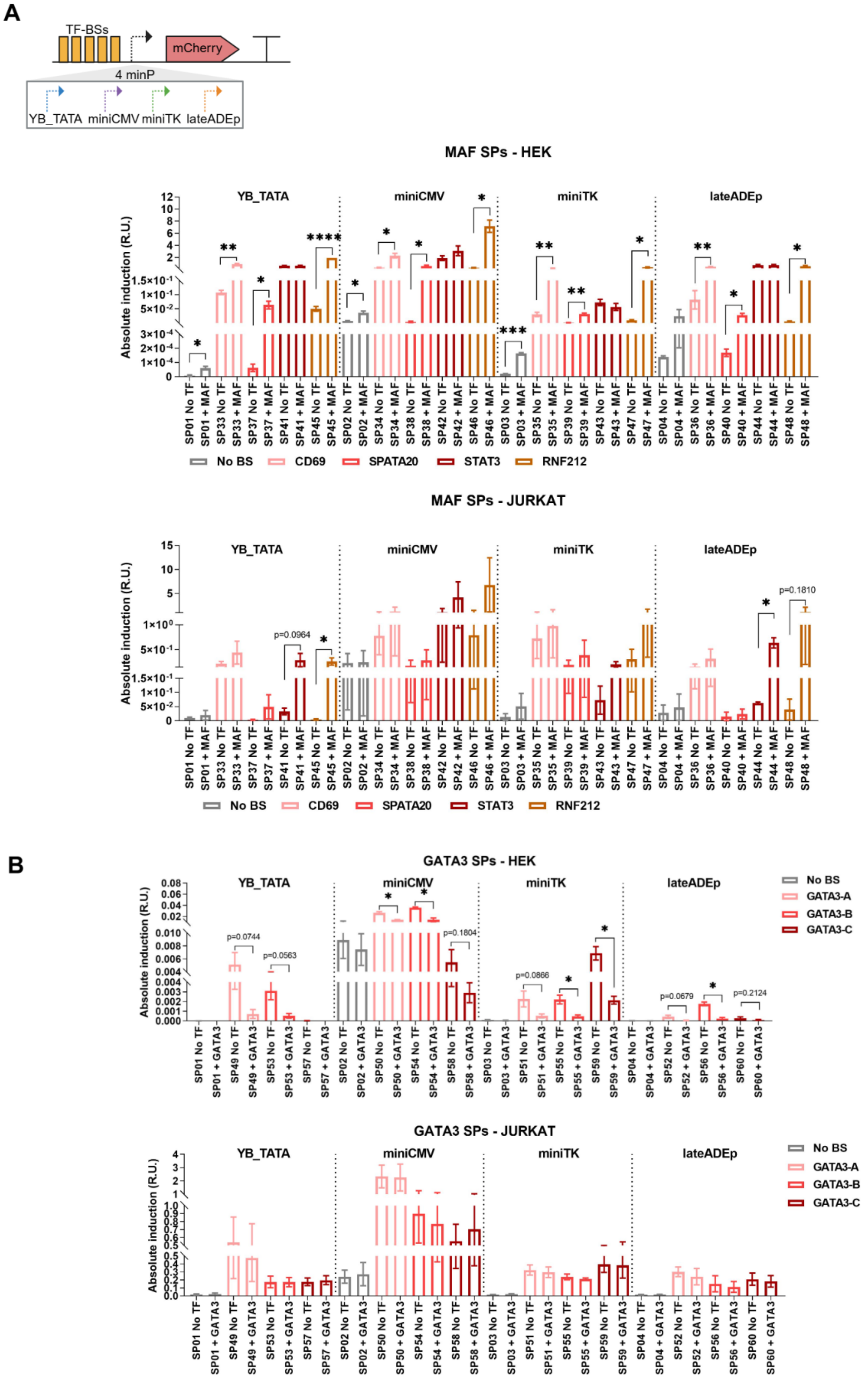

**Supplementary Figure S3.7**

Figure S3.7 - Absolute Induction Profile of SPs (Bar-Graphs)

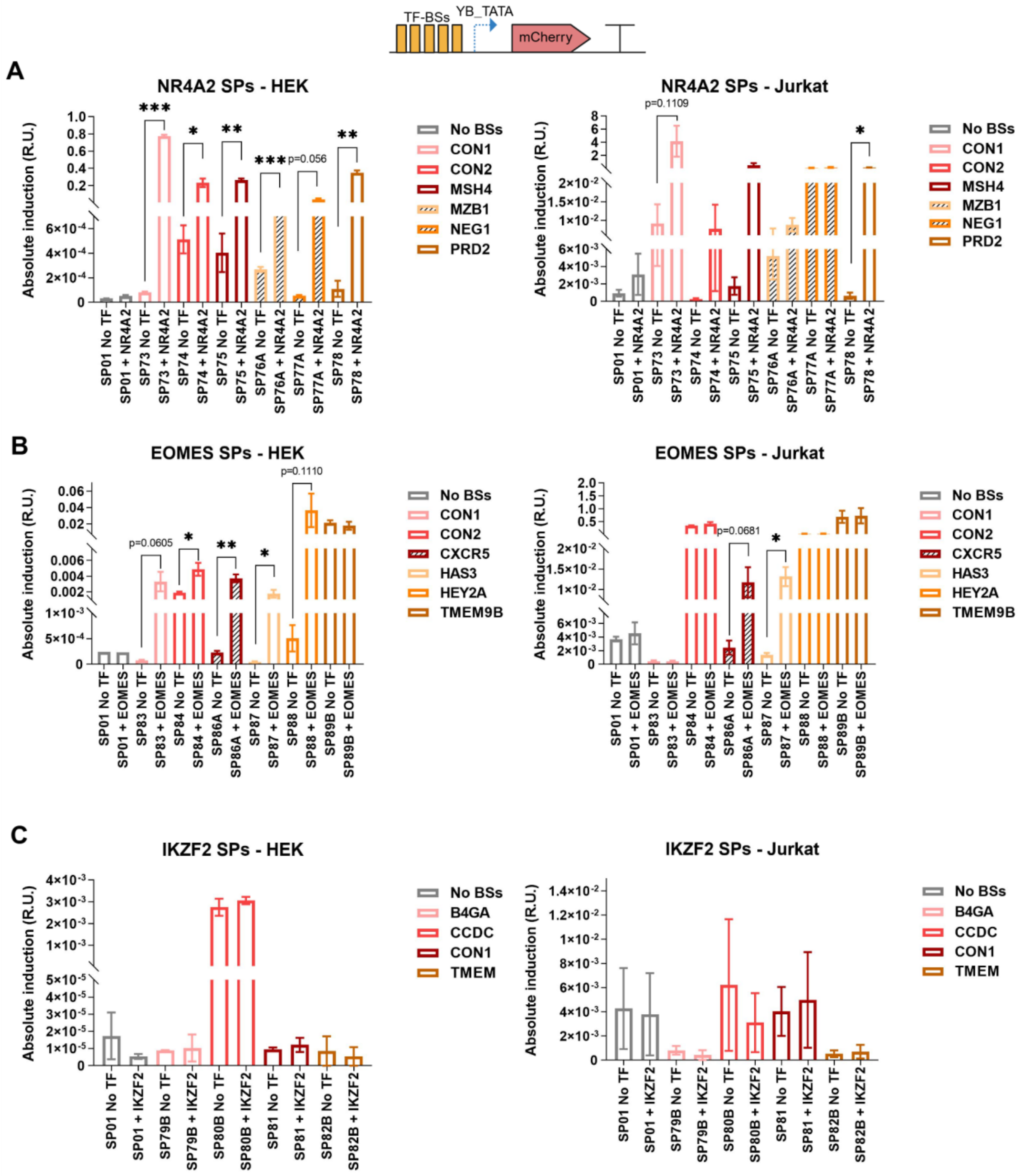

**Supplementary Figure S3.8**

SF S3.8_IRF4-BATF_YB_TATA - Fold Induction Profile of SPs (Bar-Graphs)

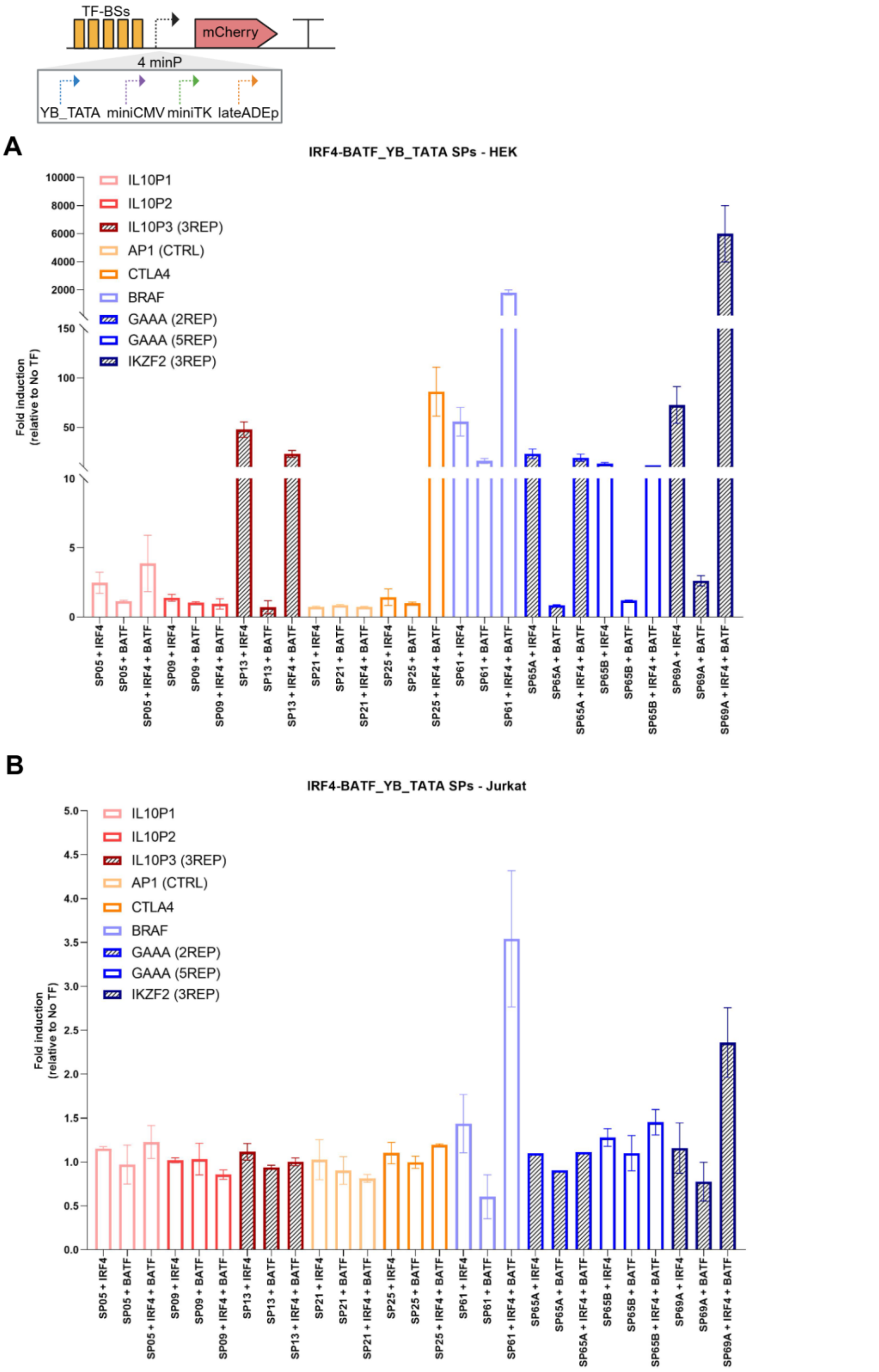

**Supplementary Figure S3.9**

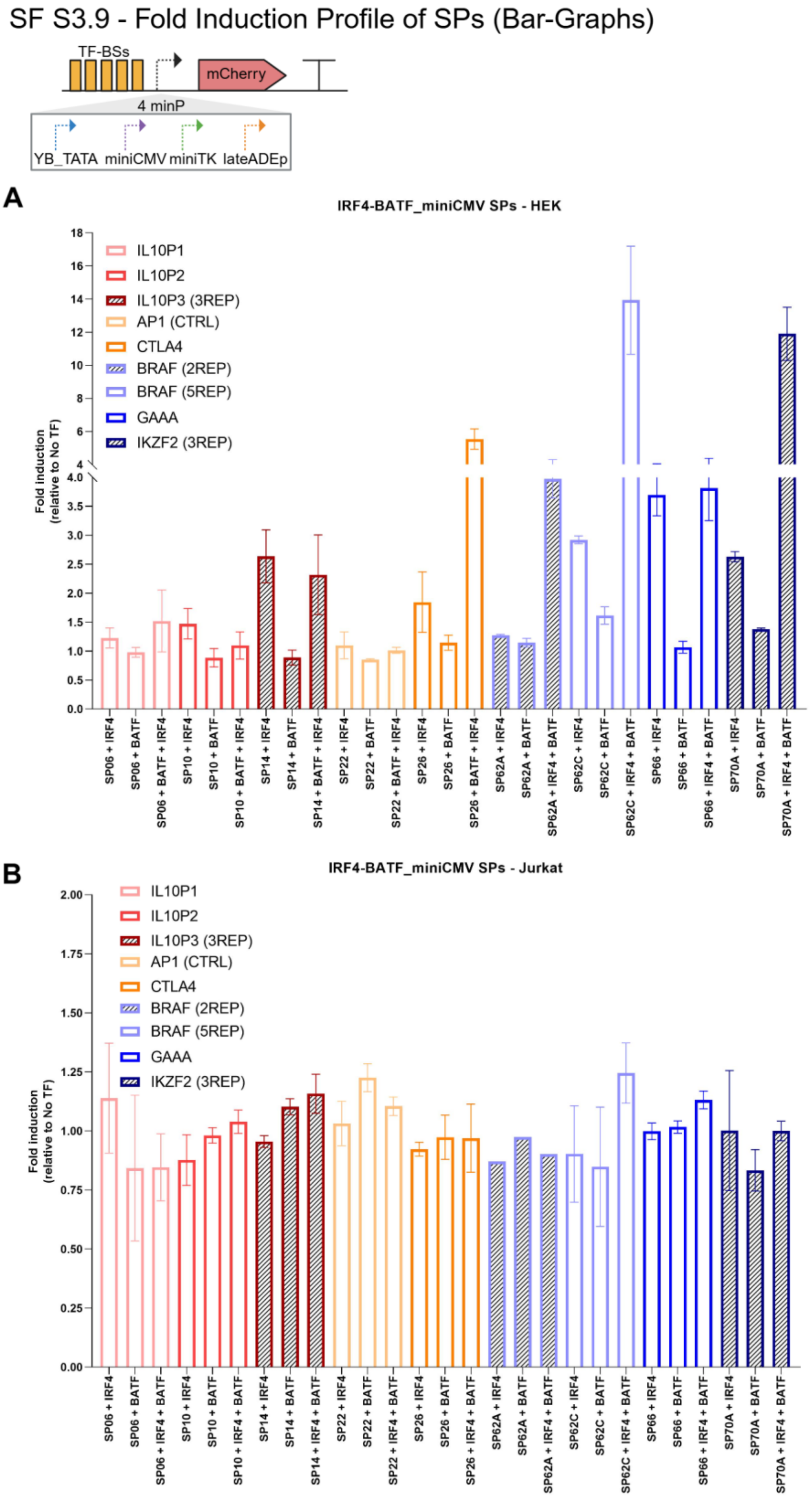

### Supplementary Figure S3.10

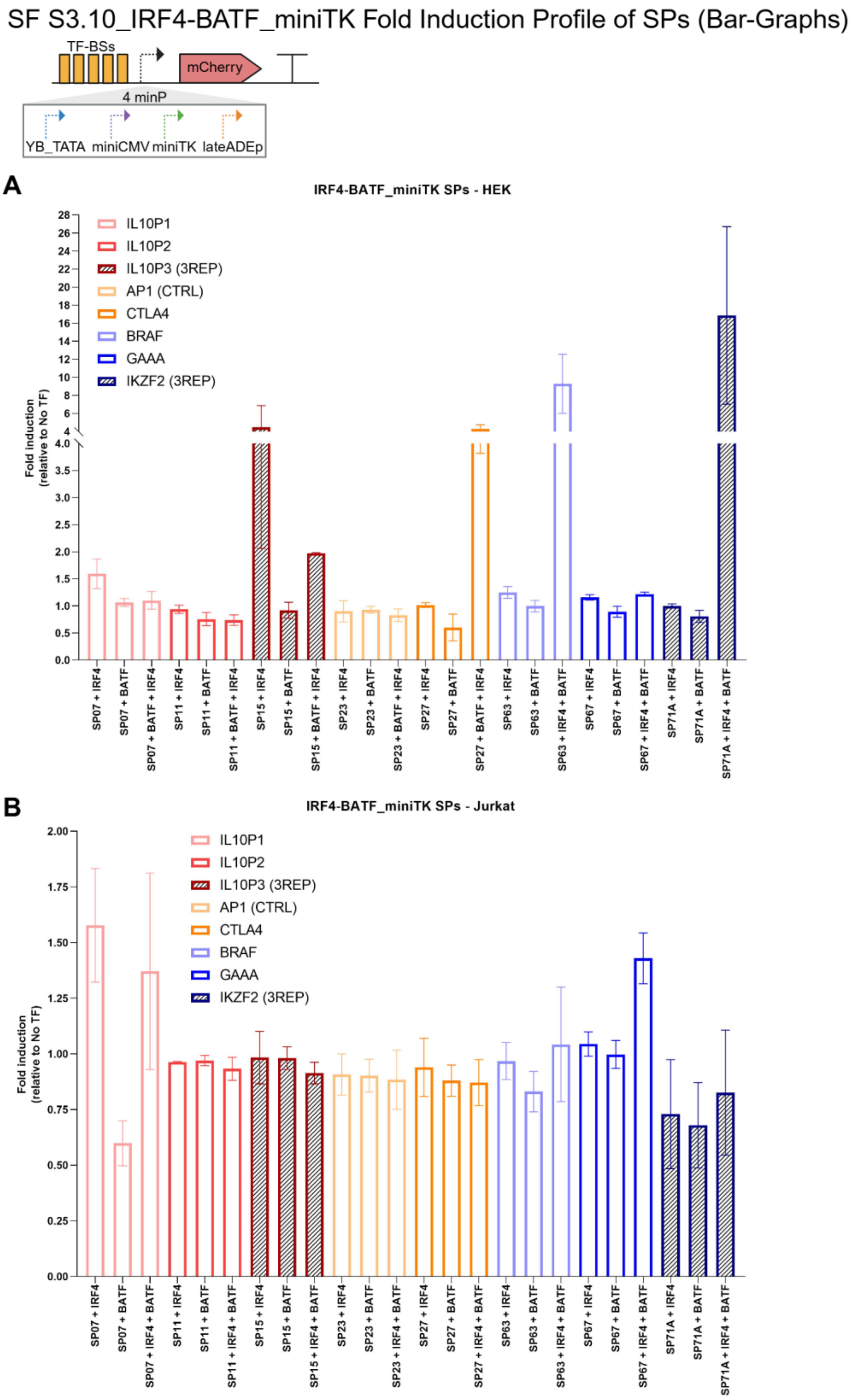

**Supplementary Figure S3.11**

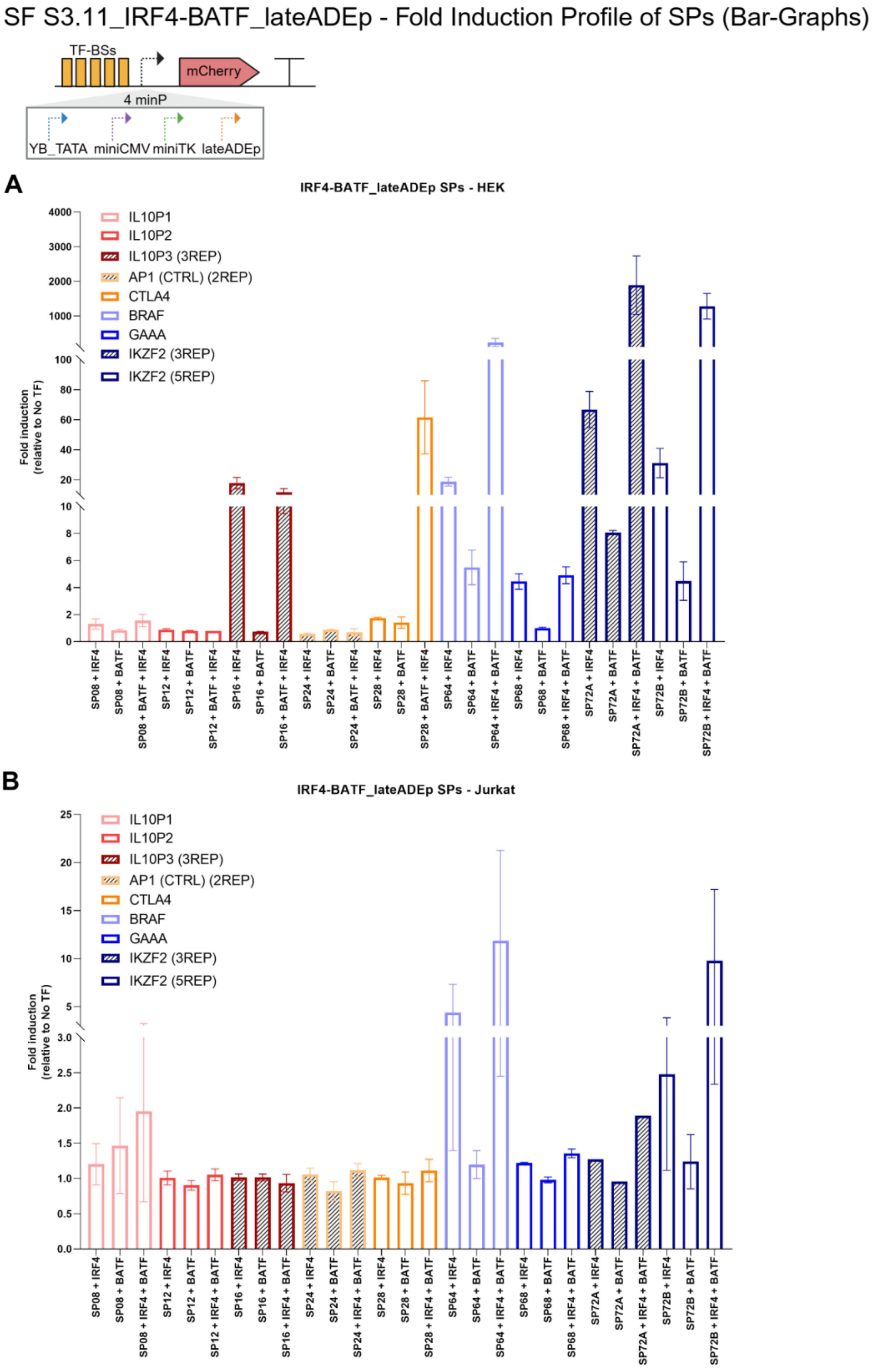

**Supplementary Figure S3.12**

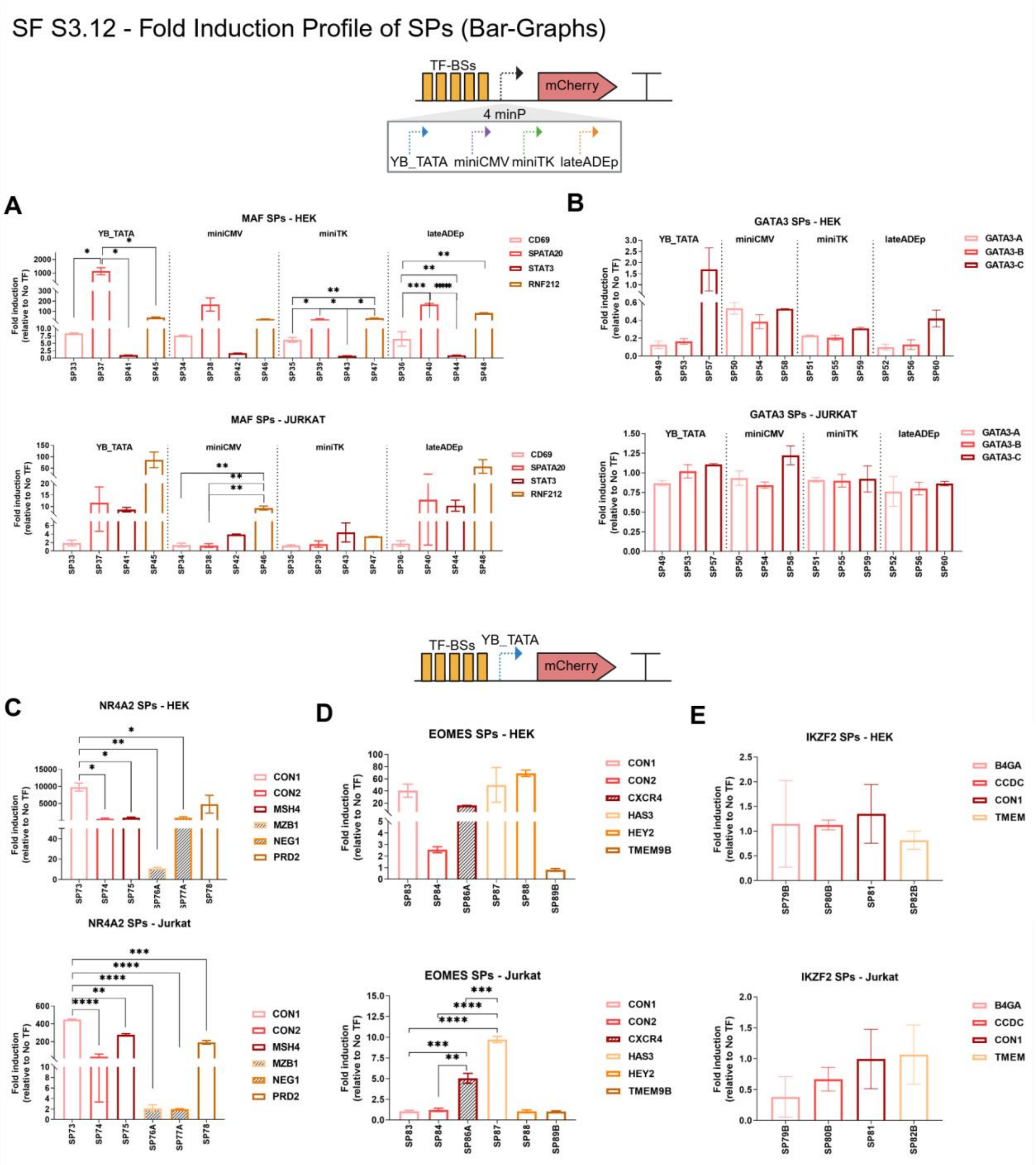

### Supplementary Figure S5

**Figure S5.1**

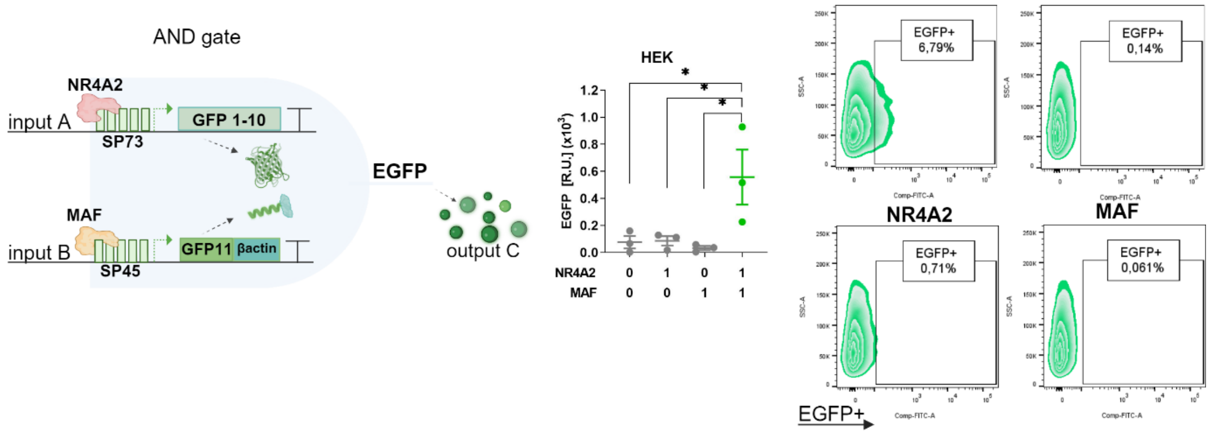

**Supplementary Figure S6.1**

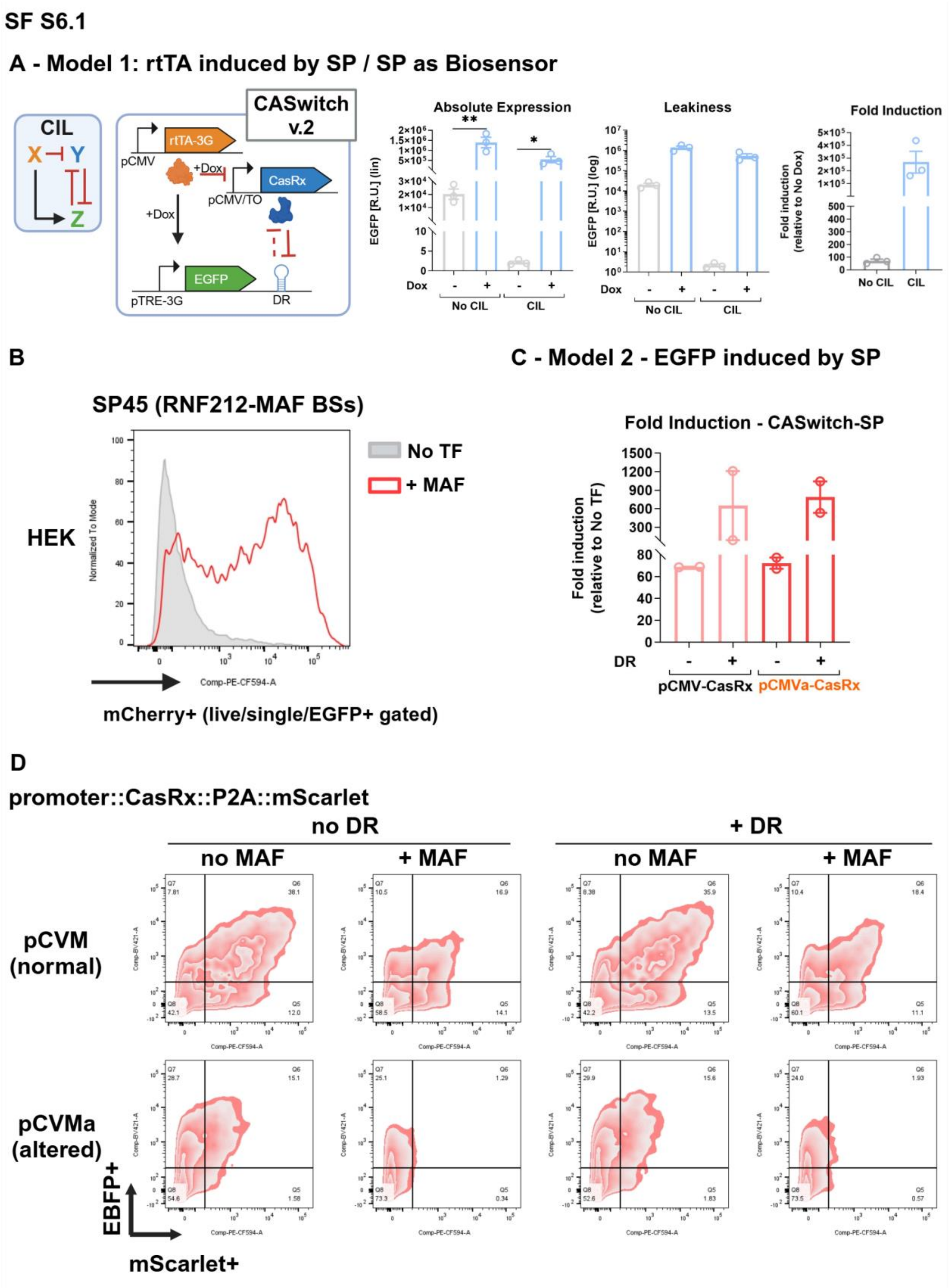

**Supplementary Figure S6.2**

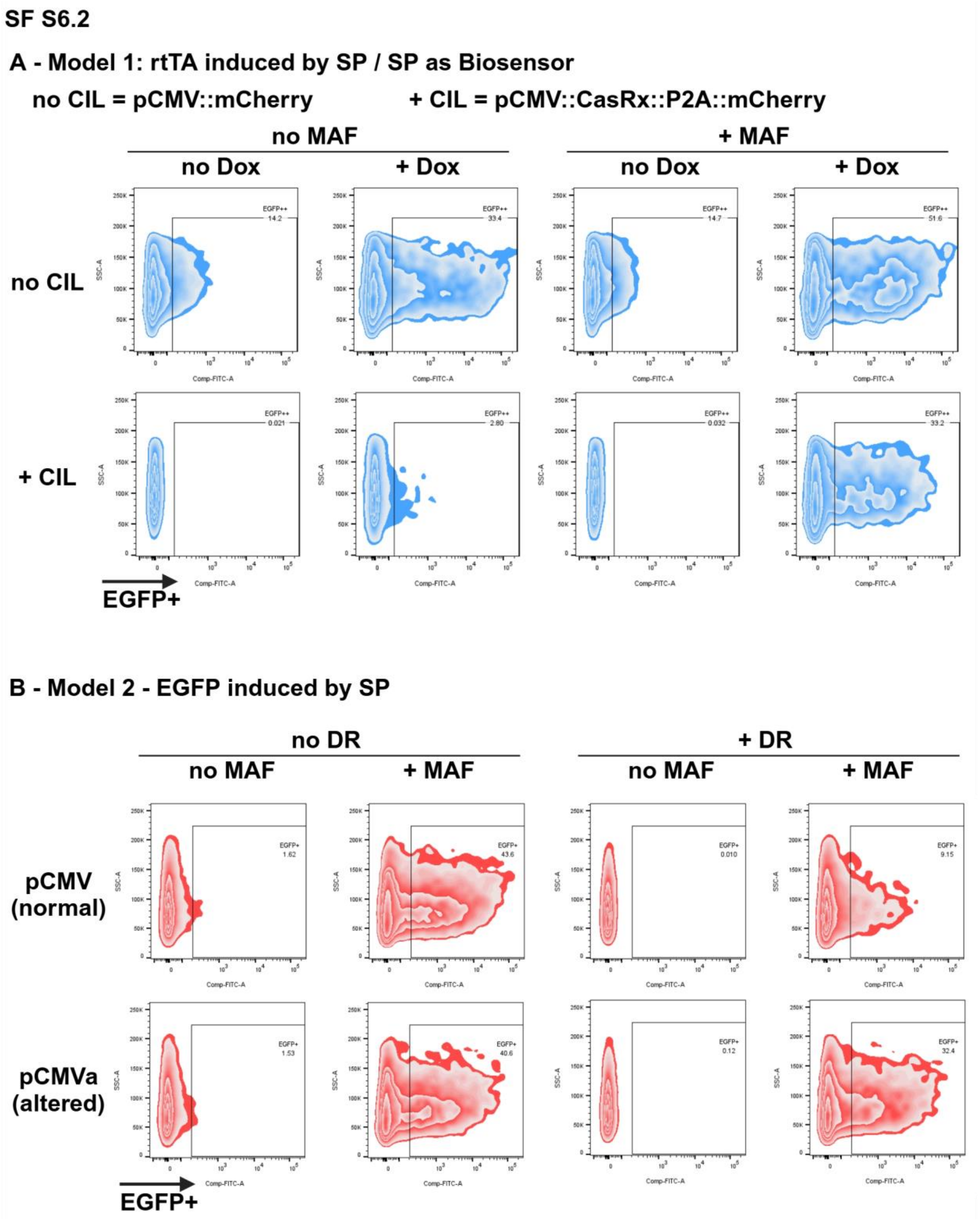

**Supplementary Figure S7**

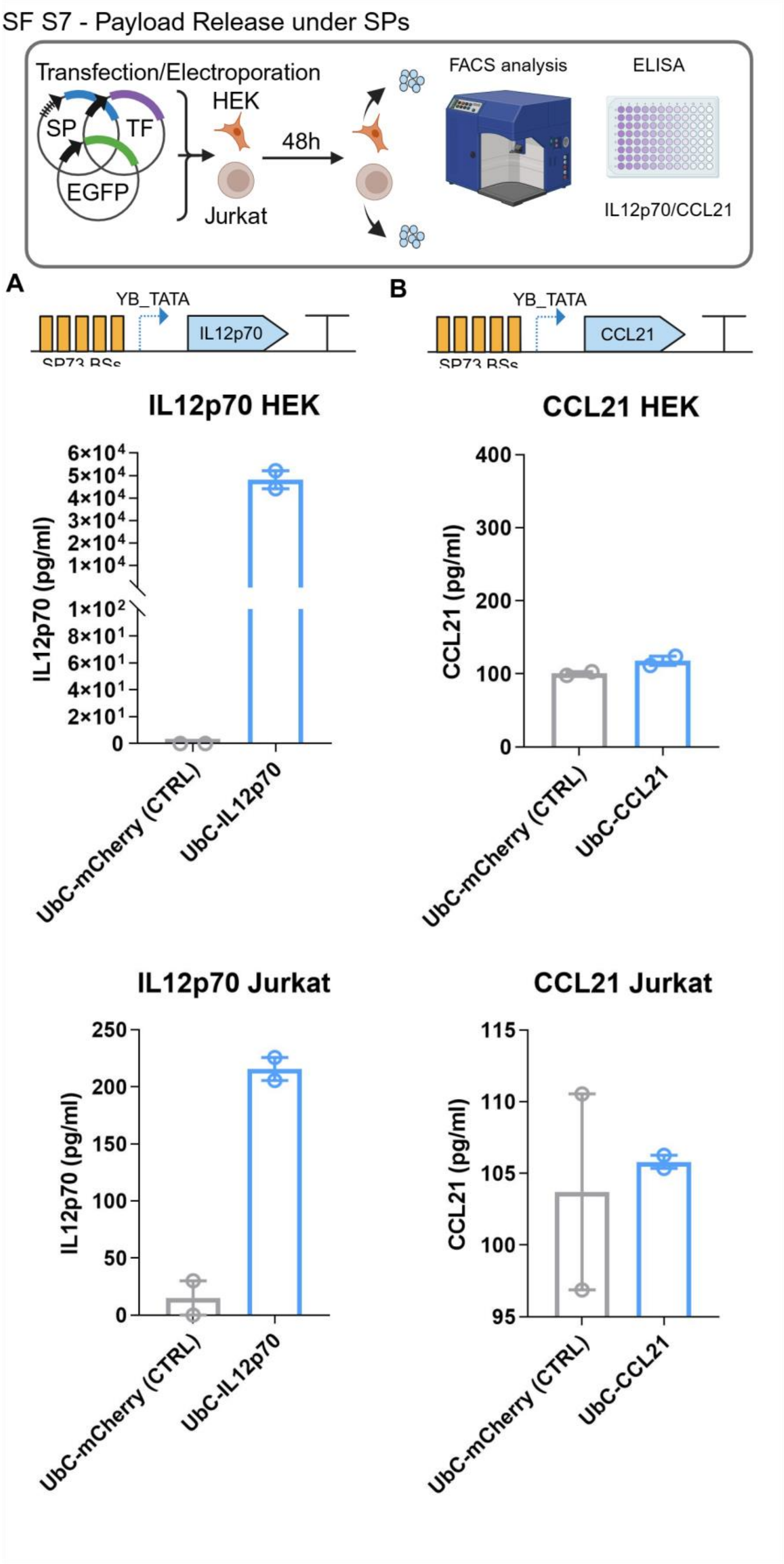
