## Supplementary Note 1 for "Design of novel synthetic promoters to tune gene expression in T cells"

**Transfer Learning and Promoter Ranking**

For the purposes of modelling exhaustion synthetic promoter ’strength’ we utilized two datasets in a transfer learning strategy. Our primary dataset of 82 exhaustion SPs is not numerous enough to directly apply a deep learning approach. Transfer learning is a technique where a pre-trained model on one task is adapted to a different but related task. To apply transfer learning to the exhaustion SPs task we use a dataset of approximately 25,000 human promoters from the Eukaryotic Promoter Database (EPD) [1].

Our secondary EPD dataset contains characterization information for each promoter in the form of the number of ChIP-seq tags matching the region 250bp to each side of the transcription start site. Our approach was to begin by training a deep neural network taking as input the EPD promoter sequences and predicting as output the expression in the form of ChIP-seq tag count.

Having trained a network on the EPD promoter task we must now approach how to use this network on the exhaustion SPs task. Our difficulty lies in the different characterization metrics used to quantify promoter ’strength’ in each dataset. To apply the EPD network to the exhaustion SPs we ranked order the SPs based on fluorescence. We attempted to recapitulate the CAR-T ’strength’ ranking using the EPD network.

**Preprocessing**:

Before use in training all EPD promoter sequences were represented as 200 bp regions and one-hot encoded into binary matrices of shape (200, 4). To account for the wide range of expression levels, we applied a Box-Cox transformation. The Box-Cox transformation is defined as:

$$y_i^{(\lambda)} = \begin{cases} \dfrac{y_i^{\lambda} - 1}{\lambda} & \text{if } \lambda \neq 0, \\ \ln y_i & \text{if } \lambda = 0, \end{cases}$$

where $y_i$ is the original data and $\lambda$ is the transformation parameter. Intuitively a Box-Cox transform attempts to move the data closer to a normal distribution. We follow the lead of Zrimec et al. [2] in this approach. Training on normally distributed data confers the benefit of maintaining balanced weight updates during backpropagation leading to faster convergence of the network.

To augment the EPD dataset, we applied a sliding window approach with a 5 bp stride over each 200 bp sequence. For each stride position, we added the original sequence, its complement, and its reverse complement. This approach has the benefit of providing us more training and test instances and attempting to train for translational invariance, allowing the network to identify predictive motifs even when their position on the strand is different.

Finally, we split the augmented EPD dataset for training and testing with an 80/20 split. After pre-processing and augmentation we have 351,540 training instances and 87,885 test instances.

Our EPDNet architecture comprises a hybrid approach combining convolutional layers and gated recurrent units (**Fig. 4A**). Convolutional layers slide a weighted window across the encoded sequence. Crucially convolutional layers allow the network to decide for itself which motifs are most predictive of promoter ’strength’. It is necessary to take this approach given the lack of knowledge about the structure of the promoter ’grammar’.

After each convolutional layer we apply batch normalization. This technique normalizes the input of each layer by adjusting and scaling the activations. It helps stabilize and accelerate training by reducing internal covariate shift, making the network less sensitive to initialization and learning rate. Max pooling is used to down-sample the content of each convolution window extracting only the most highly weighted feature. This helps to reduce computation and control overfitting.

Gated Recurrent Units (GRUs) are designed to efficiently capture sequential dependencies in data. They use gating mechanisms to control the flow of information, allowing the network to maintain and update relevant context over longer sequences. By combining convolutions and GRUs our network can leverage both short and long range motifs in its predictions. Finally we process the input through two dense layers to a final ReLU output which gives the predicted promoter ’strength’.

We train our network to minimize the Huber loss function using the Adam optimizer on an Nvidea L4 GPU. The Huber loss function is defined as:

$$L_\delta(y,\hat{y}) = \begin{cases} \frac{1}{2}\cdot(y-\hat{y})^2 & \text{if } |y-\hat{y}| \le \delta, \\ \delta\cdot\left(|y-\hat{y}| - \frac{1}{2}\delta\right) & \text{if } |y-\hat{y}| > \delta, \end{cases}$$

where $y$ is the true value, $\hat{y}$ is the predicted value, and $\delta$ is a threshold parameter. Intuitively the Huber loss attempts to be sensitive to outliers by switching between mean squared error for small errors and mean absolute error for large errors. We follow Kotopka et al. [3] in the use of the Huber loss function in a sequence-to-function network.

In **Figure 4A** we can see the progression during training of the EPD network. Training proceeds smoothly with no signs of over-fitting. Finally, we apply the trained network to the test data achieving a test loss of 0.03 and a mean absolute error or 0.25.

### Real vs Predicted Rank

After training on the EPD dataset, the neural network was applied to the 82 Synthetic Promoters. The fluorescence-based absolute mean induction of these promoters was Box-Cox transformed, and the model produced a predicted ranking of the sequences based on their expected ’strength’. In each case we use the mean absolute induction when the promoter is at its strongest expression to build the ranking. The correlation between the predicted and actual rankings of the Synthetic Promoters was 0.49, indicating a moderate but significant agreement. **Figure 4B** illuminates the exact differences in the correct and predicted ranking for each individual promoter. This result demonstrates that the transfer learning approach, leveraging a large dataset of human promoters, effectively captures relevant features for predicting gene expression in a distinct Synthetic Promoter context despite the limited dataset size.

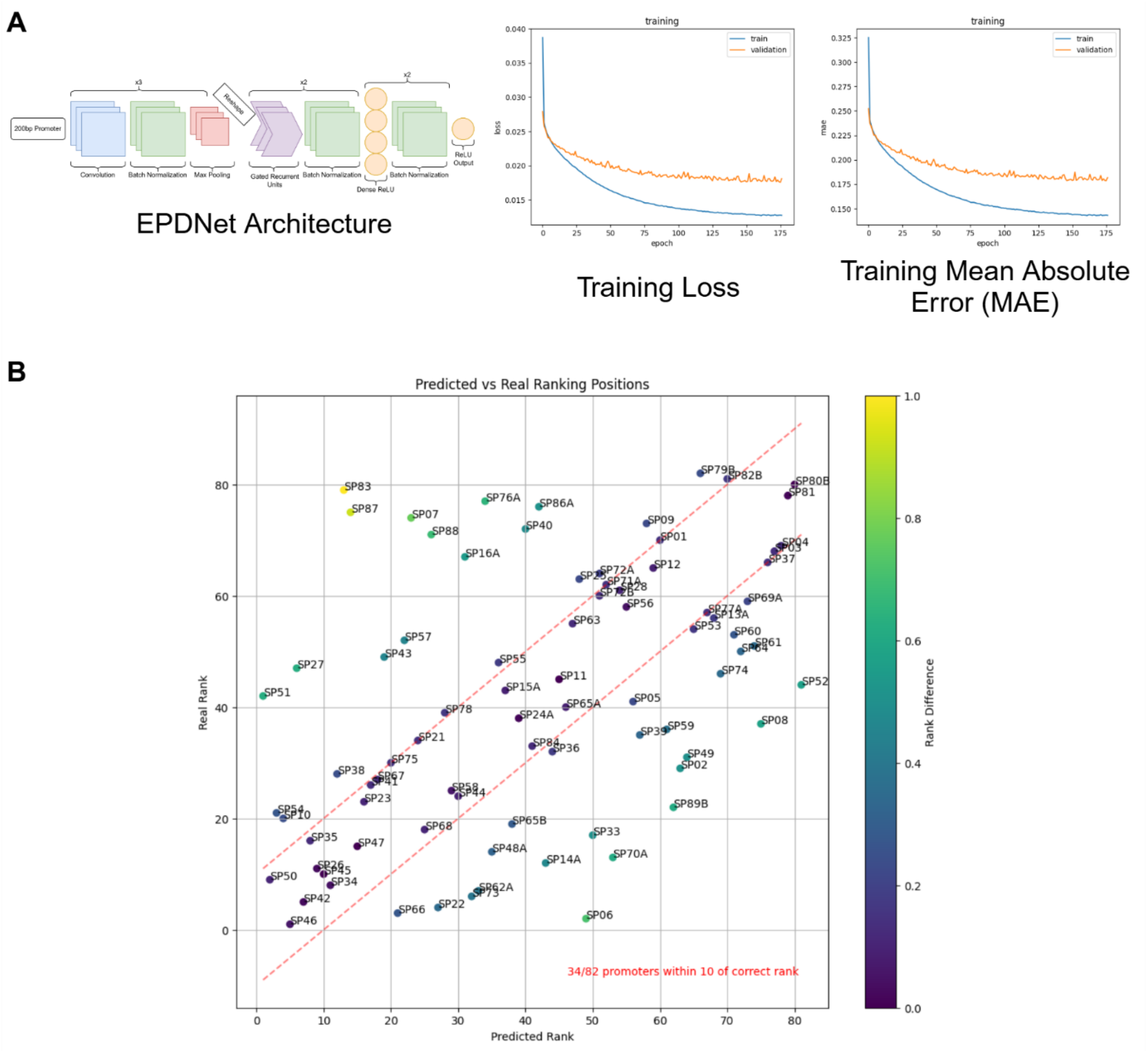


**Figure 4. Transfer learning model performance showing measured vs predicted ranking.** (**A**) Model architecture and training procedure. (**B)** The model produced a predicted ranking of the sequences based on their expected 'strength'. The mean absolute induction when the promoter is at its strongest expression was used to build the ranking to highlight the exact differences in the correct and predicted ranking for each individual promoter. Created with BioRender.com.
