## Supplementary Table S1 for "Design of novel synthetic promoters to tune gene expression in T cells"

**Supplementary Table S1 - TPM_TFs_exh_cond**

| TF | DICE | Liblau | June | Wherry |
|---|---|---|---|---|
| GTF2IRD1 | 1.10 | 0.01 | 1.74 | 0.03 |
| ZBTB32 | 51.81 | 1.25 | 0.28 | 2.86 |
| HMGB3 | 1.81 | 3.88 | 55.73 | 7.69 |
| IKZF2 | 26.23 | 0.99 | 6.03 | 2.34 |
| FOXP3 | 2.30 | 0.27 | 2.80 | 1.43 |
| PRDM1 | 2.63 | 7.17 | 15.72 | 9.29 |
| MXD1 | 13.20 | 4.15 | 3.98 | 2.24 |
| PMS1 | 17.06 | 0.72 | 14.76 | 1.03 |
| ZNF76 | 30.09 | 7.13 | 4.16 | 2.61 |
| SPI1 | 0.47 | 0.29 | 0.00 | 2.83 |
| RORA | 33.72 | 3.09 | 5.66 | 1.62 |
| TBX21 | 521.71 | 51.21 | 6.25 | 27.46 |
| FOSL2 | 301.86 | 95.45 | 1.06 | 35.28 |
| SP140 | 22.32 | 1.42 | 61.13 | 4.86 |
| TCF7 | 288.48 | 36.77 | 0.34 | 76.34 |
| MEF2C | 0.18 | 0.01 | 0.02 | 0.13 |
| ZBTB25 | 21.60 | 3.72 | 6.97 | 4.45 |
| LRRFIP2 | 11.97 | 1.26 | 9.79 | 1.45 |
| CREM | 225.79 | 18.59 | 244.81 | 1.91 |
| HIVEP1 | 98.15 | 2.10 | 8.41 | 2.69 |
| XBP1 | 328.77 | 454.56 | 71.29 | 84.19 |
| HIF1A | 440.03 | 23.59 | 11.49 | 8.05 |
| MYBL2 | 0.18 | 0.11 | 1.69 | 2.49 |
| GMEB2 | 29.65 | 8.14 | 0.49 | 4.24 |
| KLF8 | 0.42 | 0.04 | 0.50 | 0.16 |
| TSC22D1 | 10.11 | 0.64 | 9.23 | 1.73 |
| NFAT5 | 128.21 | 1.80 | 5.51 | 4.40 |
| TRPS1 | 4.47 | 0.22 | 2.52 | 0.85 |
| MEIS3 | 0.01 | 0.01 | 0.00 | 0.55 |
| PBX4 | 67.97 | 4.10 | 1.20 | 0.96 |
| ERF | 59.36 | 8.47 | 0.13 | 11.31 |
| AHR | 55.77 | 0.54 | 10.98 | 0.81 |
| GATA3 | 21.97 | 40.78 | 46.04 | 23.31 |
| TFAM | 119.33 | 37.07 | 29.15 | 16.29 |
| ELK3 | 88.47 | 1.43 | 9.63 | 3.69 |
| FOXM1 | 1.05 | 0.16 | 7.40 | 1.96 |
| VDR | 2.73 | 0.08 | 11.34 | 0.17 |
| BACH2 | 431.65 | 0.76 | 2.80 | 0.20 |
| E2F3 | 32.34 | 0.56 | 5.08 | 2.76 |
| NR3C1 | 94.49 | 9.29 | 26.34 | 6.19 |
| HMGXB3 | 85.09 | 7.34 | 3.11 | 3.29 |
| BCL6 | 30.57 | 8.85 | 1.35 | 4.77 |
| TFDP2 | 5.86 | 2.21 | 5.69 | 1.26 |
| FOXP1 | 129.36 | 3.71 | 5.53 | 2.01 |
| OTX1 | 0.00 | 0.04 | 0.10 | 0.58 |
| ID2 | 132.06 | 768.96 | 1135.92 | 75.29 |
| PLEK | 11.31 | 14.31 | 3.10 | 24.89 |
| EPAS1 | 1.54 | 0.49 | 14.36 | 0.34 |
| ARID3A | 1.41 | 0.17 | 0.42 | 1.30 |
| VAX2 | 0.00 | 0.00 | 0.00 | 0.04 |
| NFE2L2 | 213.48 | 4.51 | 64.79 | 1.35 |
| PRRX1 | 0.00 | 0.00 | 0.00 | 0.07 |
| MEF2D | 45.16 | 17.55 | 0.82 | 14.84 |

| | | | | |
|---|---|---|---|---|
| ETV3 | 66.27 | 14.15 | 1.52 | 19.42 |
| ID3 | 87.20 | 172.13 | 14.29 | 7.15 |
| IRF6 | 0.97 | 0.24 | 0.38 | 0.23 |
| KLF7 | 33.73 | 1.08 | 0.64 | 0.93 |
| PLAGL1 | 3.36 | 0.15 | 6.73 | 0.12 |
| MYB | 205.13 | 0.15 | 6.62 | 0.95 |
| KLF9 | 123.66 | 12.14 | 2.76 | 18.19 |
| NR4A3 | 1101.03 | 3.94 | 2.42 | 4.40 |
| BCL11A | 0.25 | 0.01 | 0.00 | 0.14 |
| MXI1 | 34.69 | 5.58 | 4.61 | 2.14 |
| ETF1 | 485.27 | 33.98 | 34.63 | 14.95 |
| EGR1 | 914.76 | 17.55 | 7.17 | 41.05 |
| TSHZ3 | 0.42 | 0.01 | 0.19 | 0.08 |
| EGR2 | 706.47 | 2.68 | 5.13 | 4.95 |
| BHLHE41 | 0.04 | 0.01 | 1.46 | 24.67 |
| NR4A1 | 529.98 | 3.22 | 0.78 | 2.13 |
| NFE2 | 0.04 | 0.39 | 0.00 | 0.41 |
| IKZF4 | 23.35 | 0.59 | 6.98 | 0.60 |
| TOX2 | 0.50 | 0.09 | 6.03 | 0.33 |
| TRERF1 | 5.18 | 0.67 | 1.66 | 2.05 |
| SOX4 | 2.09 | 22.92 | 59.97 | 10.64 |
| IRF1 | 470.31 | 67.08 | 2.30 | 38.21 |
| OVOL2 | 0.00 | 0.00 | 0.01 | 0.03 |
| THRA | 9.21 | 2.88 | 0.52 | 3.45 |
| STAT5A | 729.39 | 43.75 | 14.78 | 21.22 |
| HIVEP3 | 46.78 | 0.14 | 0.52 | 0.54 |
| KLF2 | 63.43 | 338.66 | 1.36 | 398.57 |
| ATF4 | 612.04 | 733.50 | 136.68 | 291.10 |
| IRF5 | 0.54 | 2.36 | 0.68 | 1.89 |
| E2F8 | 0.01 | 0.01 | 1.52 | 0.53 |
| MIS18BP1 | 10.93 | 4.68 | 39.90 | 7.98 |
| TULP4 | 13.68 | 1.11 | 0.74 | 2.15 |
| CASZ1 | 4.96 | 1.68 | 0.04 | 1.10 |
| NFATC1 | 88.77 | 2.05 | 1.60 | 2.26 |
| ZBED3 | 11.29 | 1.00 | 0.05 | 0.89 |
| BHLHE40 | 618.70 | 401.19 | 179.68 | 131.75 |
| GRHL1 | 0.42 | 0.04 | 0.14 | 0.09 |
| SETDB2 | 22.07 | 1.08 | 0.58 | 2.65 |
| BRIP1 | 0.41 | 0.04 | 4.50 | 0.24 |
| MYC | 2295.21 | 69.18 | 5.26 | 29.76 |
| IRF4 | 2730.74 | 4.70 | 4.23 | 9.35 |
| TCF19 | 3.81 | 3.27 | 0.35 | 11.45 |
| CARF | 5.75 | 0.25 | 1.90 | 0.87 |
| PRDM5 | 0.38 | 0.01 | 0.01 | 0.02 |
| LEF1 | 361.23 | 2.77 | 2.16 | 5.81 |
| JDP2 | 0.23 | 0.04 | 0.24 | 0.33 |
| TCF12 | 33.52 | 0.40 | 11.02 | 0.89 |
| ZFHX3 | 0.34 | 0.00 | 0.68 | 0.18 |
| IRF8 | 450.29 | 2.60 | 0.99 | 1.84 |
| ZNF750 | 0.62 | 0.05 | 0.00 | 0.32 |
| PRDM15 | 20.47 | 0.49 | 0.37 | 1.30 |
| RERE | 14.67 | 0.35 | 1.09 | 1.31 |
| PRDM16 | 0.02 | 0.00 | 0.00 | 0.03 |
| AFF3 | 0.27 | 0.04 | 0.10 | 0.05 |
| CSRNP1 | 504.64 | 220.33 | 1.21 | 59.33 |

| | | | | |
|---|---|---|---|---|
| ZEB1 | 74.36 | 3.65 | 22.17 | 2.63 |
| TCF7L2 | 5.84 | 0.43 | 2.65 | 0.39 |
| FOXO1 | 76.57 | 6.42 | 9.70 | 5.35 |
| THRB | 0.06 | 0.01 | 0.45 | 0.01 |
| SETBP1 | 0.08 | 0.05 | 1.73 | 0.38 |
| TCF7L1 | 0.70 | 0.03 | 0.05 | 0.08 |
| CARHSP1 | 15.90 | 12.76 | 4.40 | 7.11 |
| NR4A2 | 1336.80 | 207.98 | 14.16 | 29.96 |
| JAZF1 | 8.75 | 0.18 | 7.68 | 0.31 |
| KLF10 | 438.02 | 87.96 | 21.32 | 44.49 |
| BATF | 53.60 | 2.94 | 12.06 | 5.29 |
| BACH1 | 52.32 | 0.71 | 13.44 | 0.91 |
| ZSCAN12 | 13.08 | 2.70 | 2.53 | 2.93 |
| ELK4 | 98.51 | 5.64 | 5.75 | 36.14 |
| NR2F6 | 3.08 | 0.31 | 1.76 | 0.82 |
| ZNF208 | 0.89 | 0.03 | 0.00 | 0.08 |
| ZNF382 | 4.04 | 0.26 | 0.61 | 1.03 |
| IKZF3 | 157.91 | 4.39 | 16.52 | 10.50 |
| BCL6B | 0.24 | 0.01 | 0.00 | 0.56 |
| NFIA | 0.32 | 0.11 | 0.49 | 0.07 |
| GFI1 | 390.21 | 17.55 | 20.00 | 11.46 |
| EOMES | 24.98 | 64.92 | 0.58 | 49.03 |
| HEYL | 0.11 | 0.00 | 0.00 | 0.29 |
| HMGB2 | 26.79 | 746.48 | 2340.35 | 686.47 |
| HEY1 | 0.20 | 0.01 | 0.00 | 0.71 |
| OSR2 | 0.01 | 0.78 | 0.53 | 0.32 |
| NFIL3 | 47.25 | 5.97 | 19.22 | 8.32 |
| ZNF22 | 75.87 | 114.46 | 156.02 | 14.90 |
| E2F7 | 0.01 | 0.02 | 0.84 | 0.32 |
| ZMAT1 | 19.55 | 0.48 | 0.00 | 1.04 |
| ATMIN | 49.19 | 26.04 | 6.92 | 31.71 |
| SMAD3 | 30.39 | 1.34 | 8.24 | 3.73 |
| PBX3 | 8.37 | 0.25 | 3.82 | 0.21 |
| ZNF23 | 1.84 | 3.33 | 5.78 | 0.53 |
| RBPJ | 84.89 | 1.82 | 465.17 | 0.81 |
| CDKN2AIP | 58.75 | 164.16 | 8.63 | 48.60 |
| STAT3 | 329.47 | 29.36 | 16.82 | 11.32 |
| ATF5 | 13.37 | 7.18 | 1.21 | 5.19 |
| ZEB2 | 3.73 | 6.93 | 2.34 | 7.82 |
| FOS | 393.19 | 4090.64 | 47.23 | 1028.59 |
| SMAD1 | 0.81 | 0.03 | 1.18 | 0.06 |
| NPAS2 | 5.05 | 0.02 | 2.30 | 0.04 |
| ZIK1 | 12.47 | 3.22 | 0.47 | 1.43 |
| ARNT2 | 0.05 | 0.00 | 0.30 | 0.04 |
| AFF1 | 56.10 | 0.56 | 3.41 | 3.17 |
| RARG | 10.16 | 4.08 | 0.74 | 1.94 |
| GLIS1 | 0.01 | 0.00 | 0.00 | 0.01 |
| NR1D2 | 61.71 | 22.48 | 2.71 | 12.75 |
| DMRTA1 | 0.01 | 0.09 | 0.12 | 0.47 |
| SIX5 | 0.05 | 0.37 | 0.00 | 1.24 |
| ZBTB38 | 4.25 | 4.78 | 5.55 | 9.69 |
| HIC1 | 2.75 | 1.54 | 0.49 | 6.32 |
| MAF | 0.32 | 10.84 | 4.67 | 47.56 |
| MSC | 0.14 | 34.38 | 0.04 | 24.34 |
| EGR3 | 548.99 | 0.71 | 3.24 | 5.61 |

| | | | | |
|---|---|---|---|---|
| ZNF467 | 0.26 | 1.06 | 0.13 | 1.74 |
| ZFP41 | 3.27 | 1.12 | 0.01 | 1.14 |
| CREB3L2 | 3.61 | 0.88 | 5.69 | 1.64 |
| SATB1 | 109.34 | 13.30 | 15.13 | 4.14 |
| HOXB4 | 0.69 | 0.81 | 3.47 | 8.06 |
| KCNH8 | 0.44 | 0.00 | 0.00 | 0.01 |
| POU6F1 | 10.47 | 1.34 | 1.70 | 0.90 |
| ZFP1 | 22.79 | 3.82 | 2.85 | 3.90 |
| ZFP90 | 20.42 | 1.97 | 3.67 | 3.27 |
| IRF7 | 18.66 | 39.28 | 0.82 | 24.75 |
| MYBL1 | 2.94 | 3.77 | 1.57 | 6.47 |
| RXRA | 2.03 | 0.46 | 0.30 | 0.57 |
| FOXD2 | 0.43 | 1.63 | 0.08 | 5.99 |
| TEAD1 | 0.88 | 0.00 | 1.52 | 0.02 |
| PLSCR1 | 19.73 | 2.44 | 107.73 | 2.13 |
| LITAF | 132.69 | 99.35 | 160.92 | 14.55 |
| NCOR2 | 21.51 | 1.01 | 0.38 | 4.61 |
| TCF4 | 0.31 | 0.01 | 4.46 | 0.03 |
| ARID5A | 140.53 | 311.53 | 1.25 | 28.66 |
| MAFG | 35.63 | 5.26 | 0.89 | 2.75 |
| TFDP1 | 54.33 | 1.40 | 38.07 | 6.06 |
| ZXDB | 16.82 | 19.95 | 0.32 | 23.01 |
| ZNF511 | 9.46 | 63.33 | 4.63 | 15.24 |
| WDHD1 | 5.31 | 0.17 | 11.89 | 0.65 |
| PAX9 | 0.00 | 0.03 | 0.00 | 0.27 |
| TOX | 1.98 | 0.57 | 19.60 | 1.62 |
| L3MBTL3 | 42.38 | 0.89 | 2.71 | 2.55 |
| ZBTB10 | 79.74 | 5.10 | 0.74 | 5.62 |
| MXD3 | 2.41 | 6.71 | 0.32 | 2.00 |
| REPIN1 | 42.84 | 29.06 | 2.27 | 15.89 |
| CEBPD | 0.03 | 145.66 | 2.42 | 51.82 |
| ETV5 | 0.06 | 0.01 | 1.51 | 0.06 |
| ZNF260 | 19.52 | 1.80 | 3.63 | 3.37 |
| SPIB | 0.79 | 0.13 | 0.00 | 0.20 |
