## Supplementary Table S2 for "Design of novel synthetic promoters to tune gene expression in T cells"

**General SP Library Structure and Supplementary Table S2A and B**

**A)** General structure of SP containing plasmids: **SP ([2-5xTFBS]-minP)-GOI-SV40pA**

SP General Design (5x TFBS):

GATCTTTGTTTTGATTAA(**TFBS**)CGCGATGTACGTAAG(**TFBS**)TGCTGATAAG(**TFBS**)AGACGTGGCAGTTAG(**TFBS**)GGGGATTAAG(**TFBS**)GCGATTAAGCTTTCCAATATGC(**minP**)CTAGTACTACTACCAGATAGCTTGGTACCGAGCTCG[G/A]ATCCAGCCACCGTTTAATTAAACC-**GOI-SV40pA.**

Unbold: spacers; [G/A]: YB_TATA minP=G, others minP=A.

SP Control General Design (No TFBS):

GCTAGAGCTTTCCAATATGC(**minP**)CTAGTACTACTACCAGATAGCTTGGTACCGAGCTCG[G/A]ATCCAGCCACCGTTTAATTAAACC-**GOI-SV40pA.**

Unbold: spacers, [G/A]: YB_TATA minP=G, others minP=A.

**Supplementary Table S2A. Library of SP sensor plasmids used in this study**

| Plasmid - SPs Characterization | TF Target | TFBS (rep) (Table S3) | minP (Table S4) | GOI (Table S4) |
|---|---|---|---|---|
| pSP01 | NA | No TFBS | YB_TATA | mCherry |
| pSP02 | NA | No TFBS | miniCMV | mCherry |
| pSP03 | NA | No TFBS | miniTK | mCherry |
| pSP04 | NA | No TFBS | lateADEp | mCherry |
| pSP05 | IRF4/BATF | IL10P1 (5x) | YB_TATA | mCherry |
| pSP06 | IRF4/BATF | IL10P1 (5x) | miniCMV | mCherry |
| pSP07 | IRF4/BATF | IL10P1 (5x) | miniTK | mCherry |
| pSP08 | IRF4/BATF | IL10P1 (5x) | lateADEp | mCherry |
| pSP09 | IRF4/BATF | IL10P2 (5x) | YB_TATA | mCherry |
| pSP10 | IRF4/BATF | IL10P2 (5x) | miniCMV | mCherry |
| pSP11 | IRF4/BATF | IL10P2 (5x) | miniTK | mCherry |
| pSP12 | IRF4/BATF | IL10P2 (5x) | lateADEp | mCherry |
| pSP13A | IRF4/BATF | IL10P3 (3x) | YB_TATA | mCherry |
| pSP14A | IRF4/BATF | IL10P3 (3x) | miniCMV | mCherry |
| pSP15A | IRF4/BATF | IL10P3 (3x) | miniTK | mCherry |
| pSP16A | IRF4/BATF | IL10P3 (3x) | lateADEp | mCherry |
| pSP21 | IRF4/BATF | AP-1 (5x) | YB_TATA | mCherry |
| pSP22 | IRF4/BATF | AP-1 (5x) | miniCMV | mCherry |

| | | | | |
|---|---|---|---|---|
| pSP23 | IRF4/BATF | AP-1 (5x) | miniTK | mCherry |
| pSP24A | IRF4/BATF | AP-1 (2x) | lateADEp | mCherry |
| pSP25 | IRF4/BATF | CTLA4 (5x) | YB_TATA | mCherry |
| pSP26 | IRF4/BATF | CTLA4 (5x) | miniCMV | mCherry |
| pSP27 | IRF4/BATF | CTLA4 (5x) | miniTK | mCherry |
| pSP28 | IRF4/BATF | CTLA4 (5x) | lateADEp | mCherry |
| pSP33 | MAF | CD69 (5x) | YB_TATA | mCherry |
| pSP34 | MAF | CD69 (5x) | miniCMV | mCherry |
| pSP35 | MAF | CD69 (5x) | miniTK | mCherry |
| pSP36 | MAF | CD69 (5x) | lateADEp | mCherry |
| pSP37 | MAF | SPATA20 (5x) | YB_TATA | mCherry |
| pSP38 | MAF | SPATA20 (5x) | miniCMV | mCherry |
| pSP39 | MAF | SPATA20 (5x) | miniTK | mCherry |
| pSP40 | MAF | SPATA20 (5x) | lateADEp | mCherry |
| pSP41 | MAF | STAT3 (5x) | YB_TATA | mCherry |
| pSP42 | MAF | STAT3 (5x) | miniCMV | mCherry |
| pSP43 | MAF | STAT3 (5x) | miniTK | mCherry |
| pSP44 | MAF | STAT3 (5x) | lateADEp | mCherry |
| pSP45 | MAF | RNF212 (5x) | YB_TATA | mCherry |
| pSP46 | MAF | RNF212 (5x) | miniCMV | mCherry |
| pSP47 | MAF | RNF212 (5x) | miniTK | mCherry |
| pSP48A | MAF | RNF212 (4x) | lateADEp | mCherry |
| pSP49 | GATA3 | GATA3-A (5x) | YB_TATA | mCherry |
| pSP50 | GATA3 | GATA3-A (5x) | miniCMV | mCherry |
| pSP51 | GATA3 | GATA3-A (5x) | miniTK | mCherry |
| pSP52 | GATA3 | GATA3-A (5x) | lateADEp | mCherry |
| pSP53 | GATA3 | GATA3-B (5x) | YB_TATA | mCherry |
| pSP54 | GATA3 | GATA3-B (5x) | miniCMV | mCherry |
| pSP55 | GATA3 | GATA3-B (5x) | miniTK | mCherry |
| pSP56 | GATA3 | GATA3-B (5x) | lateADEp | mCherry |
| pSP57 | GATA3 | GATA3-C (5x) | YB_TATA | mCherry |
| pSP58 | GATA3 | GATA3-C (5x) | miniCMV | mCherry |
| pSP59 | GATA3 | GATA3-C (5x) | miniTK | mCherry |
| pSP60 | GATA3 | GATA3-C (5x) | lateADEp | mCherry |

| | | | | |
|---|---|---|---|---|
| pSP61 | IRF4/BATF | BRAF (5x) | YB_TATA | mCherry |
| pSP62A | IRF4/BATF | BRAF (2x) | miniCMV | mCherry |
| pSP62C | IRF4/BATF | BRAF (5x) | miniCMV | mCherry |
| pSP63 | IRF4/BATF | BRAF (5x) | miniTK | mCherry |
| pSP64 | IRF4/BATF | BRAF (5x) | lateADEp | mCherry |
| pSP65A | IRF4 | GAAA (2x) | YB_TATA | mCherry |
| pSP65B | IRF4 | GAAA (5x) | YB_TATA | mCherry |
| pSP66 | IRF4 | GAAA (5x) | miniCMV | mCherry |
| pSP67 | IRF4 | GAAA (5x) | miniTK | mCherry |
| pSP68 | IRF4 | GAAA (5x) | lateADEp | mCherry |
| pSP69A | IRF4/BATF | IKZF2 (3x) | YB_TATA | mCherry |
| pSP70A | IRF4/BATF | IKZF2 (3x) | miniCMV | mCherry |
| pSP71A | IRF4/BATF | IKZF2 (3x) | miniTK | mCherry |
| pSP72A | IRF4/BATF | IKZF2 (3x) | lateADEp | mCherry |
| pSP72B | IRF4/BATF | IKZF2 (5x) | lateADEp | mCherry |
| pSP73 | NR4A2 | CON1 (5x) | YB_TATA | mCherry |
| pSP74 | NR4A2 | CON2 (5x) | YB_TATA | mCherry |
| pSP75 | NR4A2 | MSH4 (5x) | YB_TATA | mCherry |
| pSP76A | NR4A2 | MZB1 (3x) | YB_TATA | mCherry |
| pSP77A | NR4A2 | NEG1 (3x) | YB_TATA | mCherry |
| pSP78 | NR4A2 | PRD2 (5x) | YB_TATA | mCherry |
| pSP79A | IKZF2 | B4GA (3x) | YB_TATA | mCherry |
| pSP79B | IKZF2 | B4GA (5x) | YB_TATA | mCherry |
| pSP80A | IKZF2 | CCDC (3x) | YB_TATA | mCherry |
| pSP80B | IKZF2 | CCDC (5x) | YB_TATA | mCherry |
| pSP80C | IKZF2 | CCDC (2x) | YB_TATA | mCherry |
| pSP81 | IKZF2 | CON1 (5x) | YB_TATA | mCherry |
| pSP82A | IKZF2 | TMEM (3x) | YB_TATA | mCherry |
| pSP82B | IKZF2 | TMEM (5x) | YB_TATA | mCherry |
| pSP83 | EOMES | CON1 (5x) | YB_TATA | mCherry |
| pSP84 | EOMES | CON2 (5x) | YB_TATA | mCherry |
| pSP86A | EOMES | CXCR5 (4x) | YB_TATA | mCherry |
| pSP87 | EOMES | HAS3 (5x) | YB_TATA | mCherry |
| pSP88 | EOMES | HEY2A (5x) | YB_TATA | mCherry |

| | | | | |
|---|---|---|---|---|
| pSP89A | EOMES | TMEM9B (4x) | YB_TATA | mCherry |
| pSP89B | EOMES | TMEM9B (5x) | YB_TATA | mCherry |

**B)** General structure of other plasmids containing SPs cloned using MTK: **SP ([5xTFBS]-minP)-GOI-SV40pA**

SP General Design (5x TFBS):

GATCTTTGTTTTGATTAA(**TFBS**)CGCGATGTACGTAAG(**TFBS**)TGCTGATAAG(**TFBS**)AGACGTGGCAGTTAG(**TFBS**)GGGGATTAAG(**TFBS**)GCGATTAAGCTTTCCAACGCGT(**minP**)CTAGTACTACTACCAGATAGCTTGGTACCGAGCTCGGATCCAGCCACT-**GOI-SV40pA.**

Unbold: spacers; Green: MTK2a (5x TF-BSs); Blue: MTK2b (minP YB_TATA+spacer) (**Supplementary Table S4**).

SP Control General Design (No TFBS):

GATTAAGCTTTCCAACGCGT(**minP**)CTAGTACTACTACCAGATAGCTTGGTACCGAGCTCGGATCCAGCCACT-**GOI-SV40pA.**

Unbold: spacers; Green: MTK2a (No TF-BSs); Blue: MTK2b (minP YB_TATA+spacer) (**Supplementary Table S4**).

**Supplementary Table S2B. Library of SP sensor plasmids used in this study**

| **Plasmid - Logic AND/OR Gates** | **TF Target** | **TFBS (rep) (Table S3)** | **minP (Table S4)** | **GOI (Table S4)** |
|---|---|---|---|---|
| TU1-TU6 | NR4A2 (SP73) | CON1 (5x) (MTK2a) | YB_TATA (MTK2b) | N7-VP16 |
| TU2 | MAF (SP45) | RNF212 (5x) (MTK2a) | YB_TATA (MTK2b) | GAL4-N8 |
| TU8 | MAF (SP45) | RNF212 (5x) (MTK2a) | YB_TATA (MTK2b) | N7-VP16 |
| TU3 | NR4A2 (SP73) | CON1 (5x) (MTK2a) | YB_TATA (MTK2b) | GFP1-10 |
| TU4 | MAF (SP45) | RNF212 (5x) (MTK2a) | YB_TATA (MTK2b) | GFP11-βActin |
| **Plasmid - CASwitch Applications** | **TF Target** | **TFBS (rep) (Table S3)** | **minP (Table S4)** | **GOI (Table S4)** |
| pGB_063 | NA | No TF-BSs (MTK2a) | YB_TATA (MTK2b) | rtTA3G |
| pGB_064 | MAF (SP45) | RNF212 (5x) (MTK2a) | YB_TATA (MTK2b) | rtTA3G |
| pGB_100 | MAF (SP45) | RNF212 (5x) (MTK2a) | YB_TATA (MTK2b) | EGFP |

| | | | | |
|---|---|---|---|---|
| pGB_101 | MAF (SP45) | RNF212 (5x)<br>(MTK2a) | YB_TATA<br>(MTK2b) | EGFP-DR |

| **Plasmid -<br>Actuator Release** | **TF Target** | **TFBS (rep)<br>(Table S3)** | **minP<br>(Table S4)** | **GOI<br>(Table S4)** |
|---|---|---|---|---|
| pGB_090 | NA | No TF-BSs<br>(MTK2a) | YB_TATA<br>(MTK2b) | IL12p70 |
| pGB_091 | NR4A2<br>(SP73) | CON1 (5x)<br>(MTK2a) | YB_TATA<br>(MTK2b) | IL12p70 |
| pGB_092 | NA | No TF-BSs<br>(MTK2a) | YB_TATA<br>(MTK2b) | CCL21 |
| pGB_093 | NR4A2<br>(SP73) | CON1 (5x)<br>(MTK2a) | YB_TATA<br>(MTK2b) | CCL21 |
