## Supplementary Table S3 for "Design of novel synthetic promoters to tune gene expression in T cells"

**Supplementary Table S3. Transcription factors binding sites (TF-BS) used for the SP sensors.**

| TF Target | TF-BS Names | TFBS Sequences (5' to 3') | Reference/Repository Databank |
|---|---|---|---|
| **IRF4/BATF** | IL10P1[a] (IL10) | TGACTCACAACGAAA | Li et al., 2012 |
| | IL10P2[a] (IL10) | GCAGTCAATTTAGAAA | Li et al., 2012 |
| | IL10P3 (IL10) | ACGAAACTGAGACAT | SELEX |
| | AP1 | TGAGTCA | Li et al., 2012 |
| | CTLA4[a] | TGTACTCAACTTGAAA | Li et al., 2012 |
| | BRAF | TGAGTCAAAATGAGA | Li et al., 2012 |
| | CON1[b] (GAAA) | CGAAACCGAAACT | JASPAR |
| | IKZF2[a] | ACAGTCAGAATGAAA | Li et al., 2012 |
| **MAF** | CD69 | ATAATTGCTGATGTAATGT | TRANSFAC |
| | SPATA20 | ATATTTGCTGAATTAAATC | TRANSFAC |
| | STAT3 | AAATGTGCTGACTCAGAGA | TRANSFAC |
| | RNF212 | AATAGTGCTGATGCTGTGT | TRANSFAC |
| **GATA3** | GATA3-A[b] | AGATAAGA | SELEX |
| | GATA3-B[b] | AAAGATAAGA | TRANSFAC |
| | GATA3-C[b] | ACTTAGAGATTTTATCCGCGTAAC | TRANSFAC |
| **NR4A2** | CON1[b] | TGACCTTTAAAGGTCA | SELEX |
| | CON2[b] | AGGTCAACTTGTGACCT | SELEX |
| | MSH4 | TCTAAAGGTCA | SELEX |
| | MZB1 (MZB1 - A SLC23A1 – A) | AGGTCACAAGGTGACCT | SELEX |
| | NEG1 (NEGR1) | TTTAAAGGTCA | SELEX |
| | PRD2 (PRDM2-A) | TGACCTTTAAAGATCA | SELEX |
| **EOMES** | CON1[b] | TAAAAGGTGTGAAAATT | TRANSFAC |
| | CON2[b] | CCGGAGGTGTCGCCTC | TRANSFAC |
| | CXCR5 | TGAAAGGTGTGAAAACA | TRANSFAC |

| | | | |
|---|---|---|---|
| | HAS3 | AAGGTGTGAAAAA | SELEX |
| | HEY2A (HEY2-A) | TAACGCCTTAAAATGTGTGA | SELEX |
| | TMEM9B | GGGGAGGTGTTGCCTA | TRANSFAC |
| **IKZF2** | B4GA (B4GALT3) | TTTAGGGATAA | TRANSFAC |
| | CCDC (CCDC91) | ATAAGGAAAAA | TRANSFAC |
| | CON1[b] | TTAAGGAAAAA | TRANSFAC |
| | TMEM (TMEM17) | TATAGGGATAA | TRANSFAC |

a: AICE - AP1–IRF composite elements (Li et al., 2012).
b: Consensus sequence for those specific TFs.
