## Supplementary Table S4 for "Design of novel synthetic promoters to tune gene expression in T cells"

| Supplementary Table S4. Sequences of minimal promoters (minP) and GOI modules used in this study. | |
|---|---|
| **Module** | **Sequence** |
| **minP** | |
| YB_TATA | TCTAGAGGGTATATAATGGGGGCCA |
| miniCMV | GTAGGCGTGTACGGTGGGAGGTCTATATAAGCAGAGCTCGTTT<br>AGTGAACCGTCAGATC |
| miniTK | TTCGCATATTAAGGTGACGCGTGTGGCCTCGAACACCGAGCGA<br>CCCTGCAGCGACCCGCTTAA |
| lateADEp-**5’UTR** | AGACGCTAGCGGGGGGCTATAAAAGGGGGTGGGGGCGTTCGT<br>CCTCACTCT**AGATCTGCGATCTAAGTACAGCTTGGCATTCCGG**<br>**TACTGTTGGTAAA** |
| **GOI** | |
| mCherry | ATGGTGAGCAAGGGCGAGGAGGATAACATGGCCATCATCAAGG<br>AGTTCATGCGCTTCAAGGTGCACATGGAGGGCTCCGTGAACGG<br>CCACGAGTTCGAGATCGAGGGCGAGGGCGAGGGCCGCCCCTA<br>CGAGGGCACCCAGACCGCCAAGCTGAAGGTGACCAAGGGTGG<br>CCCCCTGCCCTTCGCCTGGGACATCCTGTCCCCTCAGTTCATG<br>TACGGCTCCAAGGCCTACGTGAAGCACCCCGCCGACATCCCCG<br>ACTACTTGAAGCTGTCCTTCCCCGAGGGCTTCAAGTGGGAGCG<br>CGTGATGAACTTCGAGGACGGCGGCGTGGTGACCGTGACCCA<br>GGACTCCTCCCTGCAGGACGGCGAGTTCATCTACAAGGTGAAG<br>CTGCGCGGCACCAACTTCCCCTCCGACGGCCCCGTAATGCAGA<br>AGAAGACCATGGGCTGGGAGGCCTCCTCCGAGCGGATGTACC<br>CCGAGGACGGCGCCCTGAAGGGCGAGATCAAGCAGAGGCTGA<br>AGCTGAAGGACGGCGGCCACTACGACGCTGAGGTCAAGACCA<br>CCTACAAGGCCAAGAAGCCCGTGCAGCTGCCCGGCGCCTACAA<br>CGTCAACATCAAGTTGGACATCACCTCCCACAACGAGGACTACA<br>CCATCGTGGAACAGTACGAACGCGCCGAGGGCCGCCACTCCA<br>CCGGCGGCATGGACGAGCTGTACAAGTAG |
| N7-VP16 | ATGGGGGAAATTGCTGCTCTGGAAGCCAAAAATGCGGCGTTGA<br>AAGCCGAGATTGCGGCCTTGGAAGCTAAGATCGCTGCTTTAAA<br>GGCCGGATACGGCGGTTCTGGAGGTGGATCTGGCGGTTCTGAT<br>CCAAAAAAGAAGAGAAAGGTAGCCCCCCCGACCGATGTCAGCC<br>TGGGGGACGAGCTCCACTTAGACGGCGAGGACGTGGCGATGG<br>CGCATGCCGACGCGCTAGACGATTTCGATCTGGACATGTTGGG<br>GGACGGGGATTCCCCGGGTCCGGGATTTACCCCCCACGACTC<br>CGCCCCCTACGGCGCTCTGGATATGGCCGACTTCGAGTTTGAG<br>CAGATGTTTACCGATGCCCTTGGAATTGACGAGTACGGTGGGG<br>GATCCTAA |
| GAL4-N8 | ATGCACCATCACCATCACCATAAGCTACTGTCTTCTATCGAACA<br>AGCATGCGATATTTGCCGACTTAAAAAGCTCAAGTGCTCCAAAG<br>AAAAACCGAAGTGCGCCAAGTGTCTGAAGAACAACTGGGAGTG<br>TCGCTACTCTCCCAAAACCAAAAGCTCTCCGCTGACTAGGGCAC<br>ATCTGACAGAAGTGGAATCAAGGCTAGAAAGACTGGAACAGCT<br>ATTTCTACTGATTTTTCCTCGAGAAGACCTTGACATGATTTTGAA<br>AATGGATTCTTTACAGGATATAAAAGCATTGTTAACAGGATTATT |

| | |
|---|---|
| | TGTACAAGATAATGTGAATAAAGATGCCGTCACAGATAGATTGG<br>CTTCAGTGGAGACTGATATGCCTCTAACATTGAGACAGCATAGA<br>ATAAGTGCGACATCATCATCGGAAGAGAGTAGTAACAAAGGTCA<br>AAGACAGTTGACTGTATCGGGTGGTTCTGGAGGTGGATCTGGC<br>GGATCGTACGGGAAAATCGCGGCATTAAAGGCGGAGAACGCA<br>GCTCTGGAAGCCAAGATTGCAGCCTTAAAAGCGGAGATTGCTG<br>CGTTAGAGGCAGGCTACGGATCCTAA |
| GFP1-10 | ATGTCCAAAGGAGAAGAACTGTTTACCGGTGTTGTGCCAATTTT<br>GGTTGAACTCGATGGTGATGTCAACGGACATAAGTTCTCAGTGA<br>GAGGCGAAGGAGAAGGTGACGCCACCATTGGAAAATTGACTCT<br>TAAATTCATCTGTACTACTGGTAAACTTCCTGTACCATGGCCGA<br>CTCTCGTAACAACGCTTACGTACGGAGTTCAGTGCTTTTCGAGA<br>TACCCAGACCATATGAAAAGACATGACTTTTTTAAGTCGGCTAT<br>GCCTGAAGGTTACGTGCAAGAAAGAACAATTTCGTTCAAAGATG<br>ATGGAAAATATAAAACTAGAGCAGTTGTTAAATTTGAAGGAGATA<br>CTTTGGTTAACCGCATTGAACTGAAAGGAACAGATTTTAAAGAA<br>GATGGTAATATTCTTGGACACAAACTCGAATACAATTTTAATAGT<br>CATAACGTATACATCACTGCTGATAAGCAAAAGAACGGAATTAA<br>AGCGAATTTCACAGTACGCCATAATGTAGAAGATGGCAGTGTTC<br>AACTTGCCGACCATTACCAACAAAACACCCCTATTGGTGACGGT<br>CCGGTACTTCTTCCTGATAATCACTACCTCTCAACACAAACAGT<br>CCTGAGCAAAGATCCAAATGAAAAAGGAACAGGTGGCGGCGGA<br>TCCTAA |
| GFP11-βActin | ATGCGTGACCACATGGTCCTTCATGAGTATGTAAATGCTGCTGG<br>GATTACAGGTGGCGGCGGTTCTGATGATGATATCGCCGCGCTC<br>GTCGTTGACAACGGCTCCGGCATGTGCAAGGCCGGCTTCGCG<br>GGCGACGATGCCCCCCGGGCCGTCTTCCCCTCCATCGTGGGG<br>CGCCCCAGGCACCAGGGCGTGATGGTGGGCATGGGTCAGAAG<br>GATTCCTATGTGGGCGACGAGGCCCAGAGCAAGAGAGGCATCC<br>TCACCCTGAAGTACCCCATCGAGCACGGCATCGTCACCAACTG<br>GGACGACATGGAGAAAATCTGGCACCACACCTTCTACAATGAG<br>CTGCGTGTGGCTCCCGAGGAGCACCCCGTGCTGCTGACCGAG<br>GCCCCCCTGAACCCCAAGGCCAACCGCGAGAAGATGACCCAG<br>ATCATGTTTGAAACCTTCAACACCCCAGCCATGTACGTTGCTAT<br>CCAGGCTGTGCTATCCCTGTACGCCTCTGGCCGTACCACTGGC<br>ATCGTGATGGACTCCGGTGACGGGGTCACCCACACTGTGCCCA<br>TCTACGAGGGGTATGCCCTCCCCCATGCCATCCTGCGTCTGGA<br>CCTGGCTGGCCGGGACCTGACTGACTACCTCATGAAGATCCTC<br>ACCGAGCGCGGCTACAGCTTCACCACCACGGCCGAGCGGGAA<br>ATCGTGCGTGACATTAAGGAGAAGCTGTGCTACGTCGCCCTGG<br>ACTTCGAGCAAGAGATGGCCACGGCTGCTTCCAGCTCCTCCCT<br>GGAGAAGAGTTACGAGCTGCCTGACGGCCAGGTCATCACCATT<br>GGCAATGAGCGGTTCCGCTGCCCTGAGGCACTCTTCCAGCCTT<br>CCTTCCTGGGCATGGAGTCCTGTGGCATCCACGAAACTACCTT<br>CAACTCCATCATGAAGTGTGACGTGGACATCCGCAAAGACCTGT<br>ACGCCAACACAGTGCTGTCTGGCGGCACCACCATGTACCCTGG<br>CATTGCCGACAGGATGCAGAAGGAGATCACTGCCCTGGCACCC<br>AGCACAATGAAGATCAAGATCATTGCTCCTCCTGAGCGCAAGTA<br>CTCCGTGTGGATCGGCGGCTCCATCCTGGCCTCGCTGTCCACC |

| | |
|---|---|
| | TTCCAGCAGATGTGGATCAGCAAGCAGGAGTATGACGAGTCCG<br>GCCCCTCCATCGTCCACCGCAAATGCTTCGGATCCTAA |
| rtTA-3G | ATGTCTAGACTGGACAAGAGCAAAGTCATAAACTCTGCTCTGGA<br>ATTACTCAATGGAGTCGGTATCGAAGGCCTGACGACAAGGAAA<br>CTCGCTCAAAAGCTGGGAGTTGAGCAGCCTACCCTGTACTGGC<br>ACGTGAAGAACAAGCGGGCCCTGCTCGATGCCCTGCCAATCGA<br>GATGCTGGACAGGCATCATACCCACTCCTGCCCCCTGGAAGGC<br>GAGTCATGGCAAGACTTTCTGCGGAACAACGCCAAGTCATACC<br>GCTGTGCTCTCCTCTCACATCGCGACGGGGCTAAAGTGCATCT<br>CGGCACCCGCCCAACAGAGAAACAGTACGAAACCCTGGAAAAT<br>CAGCTCGCGTTCCTGTGTCAGCAAGGCTTCTCCCTGGAGAACG<br>CACTGTACGCTCTGTCCGCCGTGGGCCACTTTACACTGGGCTG<br>CGTATTGGAGGAACAGGAGCATCAAGTAGCAAAAGAGGAAAGA<br>GAGACACCTACCACCGATTCTATGCCCCCACTTCTGAAACAAGC<br>AATTGAGCTGTTCGACCGGCAGGGAGCCGAACCTGCCTTCCTT<br>TTCGGCCTGGAACTAATCATATGTGGCCTGGAGAAACAGCTAAA<br>GTGCGAAAGCGGCGGGCCGACCGACGCCCTTGACGATTTTGA<br>CTTAGACATGCTCCCAGCCGATGCCCTTGACGACTTTGACCTTG<br>ATATGCTGCCTGCTGACGCTCTTGACGATTTTGACCTTGACATG<br>CTCCCCGGGGGATCCTAA |
| EGFP | ATGGTGAGCAAGGGCGAGGAGCTGTTCACCGGGGTGGTGCCC<br>ATCCTGGTCGAGCTGGACGGCGACGTAAACGGCCACAAGTTCA<br>GCGTGTCCGGCGAGGGCGAGGGCGATGCCACCTACGGCAAGC<br>TGACCCTGAAGTTCATCTGCACCACCGGCAAGCTGCCCGTGCC<br>CTGGCCCACCCTCGTGACCACCCTGACCTACGGCGTGCAGTGC<br>TTCAGCCGCTACCCCGACCACATGAAGCAGCACGACTTCTTCAA<br>GTCCGCCATGCCCGAAGGCTACGTCCAGGAGCGCACCATCTTC<br>TTCAAGGACGACGGCAACTACAAGACCCGCGCCGAGGTGAAGT<br>TCGAGGGCGACACCCTGGTGAACCGCATCGAGCTGAAGGGCA<br>TCGACTTCAAGGAGGACGGCAACATCCTGGGGCACAAGCTGGA<br>GTACAACTACAACAGCCACAACGTCTATATCATGGCCGACAAGC<br>AGAAGAACGGCATCAAGGTGAACTTCAAGATCCGCCACAACAT<br>CGAGGACGGCAGCGTGCAGCTCGCCGACCACTACCAGCAGAA<br>CACCCCCATCGGCGACGGCCCCGTGCTGCTGCCCGACAACCA<br>CTACCTGAGCACCCAGTCCGCCCTGAGCAAAGACCCCAACGAG<br>AAGCGCGATCACATGGTCCTGCTGGAGTTCGTGACCGCCGCCG<br>GGATCACTCTCGGCATGGACGAGCTGTACAAGGGTTCTCCCAG<br>CCGGCTGGAGGAGGAGCTGAGAAGAAGACTGACCGAACCCGG<br>ATCCTAA |
| EGFP-**DR** | ATGGTGAGCAAGGGCGAGGAGCTGTTCACCGGGGTGGTGCCC<br>ATCCTGGTCGAGCTGGACGGCGACGTAAACGGCCACAAGTTCA<br>GCGTGTCCGGCGAGGGCGAGGGCGATGCCACCTACGGCAAGC<br>TGACCCTGAAGTTCATCTGCACCACCGGCAAGCTGCCCGTGCC<br>CTGGCCCACCCTCGTGACCACCCTGACCTACGGCGTGCAGTGC<br>TTCAGCCGCTACCCCGACCACATGAAGCAGCACGACTTCTTCAA<br>GTCCGCCATGCCCGAAGGCTACGTCCAGGAGCGCACCATCTTC<br>TTCAAGGACGACGGCAACTACAAGACCCGCGCCGAGGTGAAGT<br>TCGAGGGCGACACCCTGGTGAACCGCATCGAGCTGAAGGGCA<br>TCGACTTCAAGGAGGACGGCAACATCCTGGGGCACAAGCTGGA<br>GTACAACTACAACAGCCACAACGTCTATATCATGGCCGACAAGC<br>AGAAGAACGGCATCAAGGTGAACTTCAAGATCCGCCACAACAT |

| | |
|---|---|
| | CGAGGACGGCAGCGTGCAGCTCGCCGACCACTACCAGCAGAA<br>CACCCCCATCGGCGACGGCCCCGTGCTGCTGCCCGACAACCA<br>CTACCTGAGCACCCAGTCCGCCCTGAGCAAAGACCCCAACGAG<br>AAGCGCGATCACATGGTCCTGCTGGAGTTCGTGACCGCCGCCG<br>GGATCACTCTCGGCATGGACGAGCTGTACAAGGGTTCTCCCAG<br>CCGGCTGGAGGAGGAGCTGAGAAGAAGACTGACCGAACCCGG<br>ATCCTAAACTAGAGATCTATGTGAGGATC**CAAGTAAACCCCTAC**<br>**CAACTGGTCGGGGTTTGAAAC**TCGAGATCGAGCTG |
| IL12p70 | ATGTGTCACCAACAGTTGGTTATCTCTTGGTTCAGTCTTGTCTTC<br>CTGGCTTCCCCCCTGGTAGCCATCTGGGAGTTGAAGAAAGATG<br>TGTACGTGGTGGAATTGGACTGGTATCCGGACGCACCCGGTGA<br>GATGGTGGTACTTACCTGCGACACCCCCGAGGAGGACGGAATC<br>ACCTGGACTCTTGATCAGTCTAGCGAGGTCTTGGGTAGTGGAA<br>AGACCCTGACCATTCAGGTGAAGGAATTCGGAGATGCAGGGCA<br>GTACACATGTCACAAGGGTGGGGAAGTGCTCTCTCATTCCCTTC<br>TTCTCCTCCACAAAAAGGAGGACGGAATTTGGTCAACCGATATT<br>CTGAAGGATCAGAAAGAGCCAAAAAACAAAACATTCCTGAGGTG<br>TGAAGCCAAGAACTACAGCGGACGGTTTACGTGTTGGTGGCTG<br>ACTACAATTTCAACCGATCTGACTTTCAGTGTGAAGTCCTCCCG<br>CGGCAGCTCAGATCCCCAGGGGGTGACATGTGGCGCAGCGAC<br>GCTCTCCGCAGAGCGAGTGAGGGGGGATAACAAAGAATACGAA<br>TATAGCGTGGAGTGCCAGGAGGATTCCGCCTGCCCAGCCGCC<br>GAAGAGTCCCTGCCCATCGAGGTTATGGTCGACGCAGTGCATA<br>AACTCAAATACGAGAACTATACGTCCTCCTTTTTCATACGCGACA<br>TCATCAAACCTGACCCCCCAAAAAACCTTCAACTCAAACCCCTG<br>AAAAATAGTCGGCAGGTGGAGGTCAGTTGGGAATACCCGGATA<br>CTTGGAGTACTCCGCACTCCTATTTTTCCTTGACTTTTTGTGTCC<br>AGGTACAGGGAAAGAGCAAAAGGGAAAAGAAGGACCGCGTCTT<br>CACTGACAAGACATCAGCTACAGTGATCTGTCGAAAGAATGCAA<br>GCATTAGCGTGAGAGCCCAGGACAGGTACTATAGTTCTAGCTG<br>GTCAGAATGGGCAAGCGTACCTTGCTCTGGCGGTGGAGGCTCC<br>GGCGGGGGGGGGAGCGGCGGGGGAGGATCTCGGAATCTCCC<br>TGTTGCCACTCCAGACCCAGGTATGTTTCCATGCCTGCACCACT<br>CTCAGAACTTGCTCCGGGCTGTCTCTAACATGCTGCAGAAGGC<br>CAGGCAGACATTGGAGTTTTACCCCTGTACGTCTGAAGAGATCG<br>ATCATGAGGACATTACAAAGGATAAAACATCAACGGTCGAAGCC<br>TGCCTTCCCCTTGAGCTTACCAAAAATGAAAGCTGTCTGAATAG<br>CCGCGAAACATCCTTTATTACTAATGGTTCATGCCTCGCCTCCA<br>GGAAAACATCTTTTATGATGGCGCTCTGTCTCAGTTCTATTTATG<br>AGGATCTCAAAATGTATCAAGTTGAGTTCAAGACCATGAACGCC<br>AAACTGCTGATGGATCCTAAACGCCAGATCTTTTTGGATCAAAA<br>TATGCTTGCTGTCATCGACGAACTCATGCAGGCCCTGAATTTCA<br>ACTCCGAGACTGTTCCCCAAAAATCATCCCTGGAAGAGCCCGA<br>CTTTTACAAAACAAAGATCAAGCTCTGCATTCTTCTGCACGCTTT<br>TCGGATCCGAGCTGTGACTATAGACAGAGTGATGTCCTACCTGA<br>ATGCTAGTGGTTCTGGCGGTGGAGGCTCCGAGCAGAAGCTGAT<br>CAGCGAGGAGGACCTGGGATCCTAA |
| CCL21 | ATGGCTCAATCCTTGGCGCTGTCCTTGCTGATCTTGGTGCTGGC<br>ATTTGGCATTCCCCGCACACAAGGAAGCGATGGTGGCGCACAG<br>GACTGTTGCCTGAAGTATTCACAGCGGAAAATCCCAGCTAAAGT<br>GGTTCGGAGCTATCGCAAGCAGGAACCTTCCTTGGGGTGTTCA |

| | |
|---|---|
| | ATCCCAGCTATCCTCTTTCTTCCAAGAAAGCGCTCTCAGGCCGA<br>GCTGTGTGCTGACCCAAAAGAGCTGTGGGTCCAGCAGTTGATG<br>CAGCACTTGGACAAGACGCCCTCTCCCCAGAAACCCGCCCAGG<br>GCTGCAGAAAAGATAGGGGCGCCTCAAAAACAGGCAAAAAAGG<br>GAAAGGTAGCAAAGGTTGCAAACGCACCGAACGAAGCCAGACG<br>CCAAAAGGGCCAGGTTCTGGCGGTGGAGGCTCCGAGCAGAAG<br>CTGATCAGCGAGGAGGACCTGGGATCCTAA |
